# Bacterial degradation of a plant toxin and nutrient competition with commensals trade off to constrain pathogen growth

**DOI:** 10.1101/2025.11.17.688815

**Authors:** Kerstin Unger, Marco Mauri, Jonathan Gershenzon, Rosalind J. Allen, Matthew T. Agler

**Author notes:** co-corresponding authors: Matthew T. Agler, Institute of Microbiology, Plant Microbiosis Group, Friedrich Schiller University Jena, Neugasse 23, 07743 Jena, Germany, Rosalind J. Allen, Institute of Microbiology, Theoretical Microbial Ecology Group, Friedrich Schiller University Jena, Buchaer Str. 6, 07745 Jena, Germany. contributed equally (Kerstin Unger, Marco Mauri). <u>Author contributions</u>: <u>KU:</u> Conceptualization, Data Curation, Investigation, Methodology, Project Management, Visualization, Writing – Original draft, Writing – Review & Editing, <u>MM:</u> Conceptualization, Data Curation, Formal Analysis, Methodology, Project Management, Visualization, Writing – Original draft, Writing – Review & Editing, <u>JG:</u> Funding Acquisition, Resources, Writing – Review & Editing, <u>RJA:</u> Conceptualization, Funding Acquisition, Resources, Supervision, Writing – Review & Editing, <u>MTA:</u> Conceptualization, Funding Acquisition, Resources, Supervision, Writing – Original draft, Writing – Review & Editing.

## Abstract

Healthy plant leaves host both commensal bacteria, which usually do not cause harm, and opportunistic pathogens, which under the right circumstances can cause disease. Microbial and plant-derived factors can potentially govern the balance between commensals and pathogens; understanding this dynamic is crucial for developing effective biocontrol strategies. In *Arabidopsis thaliana*, isothiocyanates (ITCs) are toxic defense metabolites that suppress most bacteria. An important virulence factor for bacterial and fungal pathogens of *A. thaliana* is the ITC hydrolase SaxA, which detoxifies ITCs. To investigate microbial interactions based on SaxA-mediated ITC degradation, we used five ITC-sensitive bacterial commensals and the opportunistic pathogen *Pseudomonas viridiflava* 3D9 (PS). All strains were isolated from healthy *A. thaliana* leaves and PS degrades 4-methylsulfinylbutyl-ITC (4MSOB-ITC) with SaxA. We examined their growth in the presence of 4MSOB-ITC, both in monoculture and in coculture with PS or a *saxA*-deficient mutant (PSKO). Based on experimental growth data, we developed a generalizable consumer-resource mathematical model incorporating ITC toxicity, ITC degradation, and nutrient use. We predicted conditions and confirmed them experimentally under which SaxA not only benefits the pathogen but also indirectly favors commensal growth, which then can limit pathogen proliferation by competing for nutrients. In addition, we tested *in silico* how commensal ITC susceptibility, pathogen ITC degradation rates, and growth parameters affect the trade-off between SaxA-mediated virulence (strong pathogen growth) and high commensal rescue (commensal growth). Our findings suggest that the effects of microbial traits - traditionally viewed as either virulence or plant-beneficial factors - are context-dependent. This underscores the need to reconsider how such traits are classified in the context of plant-microbiome interactions.

**Author Summary:** Healthy plant leaves host a variety of bacteria; these can be beneficial, but some (opportunistic pathogens) can also be harmful under certain conditions. To design effective biocontrol strategies to sustainably protect plants, it is important to understand how opportunistic pathogens thrive as part of a healthy, balanced leaf microbiome. Plant defense metabolites such as isothiocyanates (ITCs) and bacterial ITC resistance mechanisms such as the ITC hydrolase SaxA may play key roles in maintaining this balance. In this study, we explore how SaxA-mediated ITC degradation by a pathogen also benefits diverse ITC-sensitive commensals and how this in turn could shape microbiome stability and plant health. Using mathematical modeling based on growth data from PS with diverse commensals, we find that interaction dynamics can be explained by ITC detoxification and nutrient competition. We predict and experimentally confirm that conditions exist under which SaxA favors commensal growth so strongly that the pathogen is outcompeted for resources, thus not benefiting from its own virulence factor. Our findings suggest that the effects of microbial traits are context-dependent, especially when functioning as public good in a community context like SaxA. Taken together, quantitative modeling of these interactions may inform strategies to maintain healthy plant microbiomes and control disease.

## Introduction

Pathogens have traditionally been defined based on Koch’s postulates, which identify a specific microbe as the causal agent of a specific disease; and the disease-causing microbe is postulated to be always present in sick but never in healthy individuals. In this framework, disease would only depend on a pathogen’s genetic capacity to produce virulence factors and evade host defenses to proliferate in a host (1). However, this view neglects how non-host or environmental factors can influence the outbreak of a disease. In plant pathology, the so-called disease triangle rationalizes this context-dependence and postulates that disease is not solely dependent on the presence of a pathogen but rather results from the interplay between plant host traits, opportunistic pathogens, and the environment. The concept has recently been extended to include interactions between pathogens and the residual plant microbiome to underline the importance of co-colonizing microbes (2). In this context, a balanced microbiome where “beneficial” microbes keep pathogens in check is associated with health and a dysbalanced microbiome with disease (3). The development of concepts of health and disease reflects a shift in research in plant pathology from the one-pathogen-one-disease model toward understanding microbial and plant traits that help sustain a balanced microbiome and promote plant health.

One important plant trait that shapes microbial colonization of roots and leaves is the presence of antimicrobial plant specialized metabolites. Microbial detoxification of antimicrobial metabolites is therefore a key microbial process that contributes to the virulence of plant pathogens (4–7). An important trait of the plant microbiome is to compete for nutrients with plant pathogens to suppress them (8,9). However, the interplay of these traits - plant defense metabolites, detoxification, and nutrient competition between pathogens and commensals - especially in the context of complex plant microbiomes is still poorly understood. In bacterial communities from the human gut, the degradation of antimicrobial drugs such as antibiotics can protect co-colonizing antibiotic-susceptible bacteria (10) and we propose the effects of degradation of antimicrobial plant defense metabolites on the plant microbiome are similar.

The plant microbiome, specifically the microbiome of healthy leaves, consists of mainly Proteobacteria, Actinobacteria and Bacteroidetes (11,12). These include both non-pathogenic commensal taxa that are often sensitive to plant defense metabolites, and opportunistic pathogens some of which can degrade plant defense metabolites. The presence of opportunistic pathogens even in healthy leaf microbiomes (13) raises questions about their interactions with commensals which may cause pathogens to shift from harmless leaf colonizers to disease-causing agents.

In the model plant *Arabidopsis thaliana,* aliphatic glucosinolates (GLS) are major leaf defense metabolites which are converted into antimicrobial isothiocyanates (ITCs) when plant cells are damaged: the so-called “mustard-oil bomb” (14). The release of ITCs can be caused for example by insect herbivory (15) or by disruption of plant cells by microbial pathogens (16). Both bacterial and fungal pathogens express virulence factors encoded by *sax* genes (<u>s</u>urvival in *<u>A</u>rabidopsis* e<u>x</u>tracts) that confer ITC resistance. These include the efflux pumps SaxF, SaxG and SaxD, as well as the ITC hydrolase SaxA (17). SaxA hydrolyzes various ITCs into their corresponding amines, rendering them non-toxic (18) and promoting virulence in various pathosystems (5,17,19).

Bacteria that do not have a specialized ITC hydrolase, including commensal leaf colonizers and non-adapted pathogens, have varying levels of ITC susceptibility ranging from highly susceptible to almost resistant (17,20). These bacteria can face very sudden exposure to ITC defense metabolites when the plant host comes under attack, and they probably also need to cope with low constitutive levels of ITCs arising from leaky transport of GLS (21), constant cycling between GLS and ITCs in intact plant tissues (22), and/or release of ITCs by GLS-utilizing bacteria (20). We previously observed that ITC-sensitive commensal bacteria can benefit from SaxA-mediated ITC degradation (23). However, it is still unclear how this public good affects commensals and pathogens in a more complex community context.

Here, we hypothesize that SaxA-mediated ITC degradation by the opportunistic plant pathogen *Pseudomonas viridiflava* not only increases its virulence but also acts as a public good to protect ITC-sensitive commensals creating a trade-off in pathogenicity by enhancing competition of the pathogen with the rescued commensals for limited nutrients. To investigate this proposition, we used the ITC-degrading opportunistic pathogen *Pseudomonas viridiflava* 3D9 (PS) and five commensals that are sensitive to 4-methylbutylsulfinyl ITC (4MSOB-ITC), a major GLS-derived defense metabolite in the widely used reference genotype *A. thaliana* Col-0 (17). All strains were previously isolated from healthy leaves of wild *A. thaliana* plants, making this a good model system for taxa that are likely to co-occur and interact in natural systems. We built a mathematical model based on our experimental measurements of microbial interactions mediated by the degradation of 4MSOB-ITC. We then used the model to predict when a tradeoff occurs that is likely to be unfavorable to the pathogen but benefits the commensals and experimentally tested these predictions. This work reveals that the benefit of the “virulence factor” SaxA for pathogen growth is highly dependent on pathogen and commensal traits and the ITC concentration supplied by the plant.

## Material and Methods

### Bacterial strains used in this study

Commensal bacterial strains used in this study were *Stenotrophomonas* sp. SrG (hereafter: E), *Plantibacter* sp. 2H11-2 (hereafter: G), *Janthinobacterium* J4 (hereafter: K), *Brevundimonas* sp. 7B5 (hereafter: M) and *Rhodococcus* sp. 6G8 (hereafter: R) and the opportunistic pathogen *Pseudomonas viridiflava* 3D9 (hereafter: PS) (Suppl. Tab. 1). A *saxA*-knock-out mutant in the PS background (hereafter: PSKO) was generated in a previous study (23). All bacteria were grown on R2A agar plates or in R2A broth (yeast extract 0.5 g/L, peptone 0.5g/L, casein hydrolysate 0.5 g/L, glucose 0.5 g/L, soluble starch 0.5 g/L, K_2_HPO_4_ 0.3 g/L, MgSO_4_ 0.024 g/L, sodium pyruvate 0.3 g/L, for plates: 15 g agar/L; pH=7.2±0.2 (24)) at 28°C, unless stated otherwise.

### Fluorescent tagging of PS and PSKO with mScarlet-I

Wildtype PS and PSKO were fluorescently labelled according to Schlechter et al. (25). Briefly, the pMRE-Tn7-145 plasmid containing the gene for the red fluorescent protein mScarlet-I as well as chloramphenicol and gentamicin resistance cassettes was conjugated from *E. coli* ST18 into both PS and PSKO strains. The plasmid was integrated into the genome by a Tn-7 transposase and bacterial cells which did not integrate the plasmid were counter selected by incubation at 35°C to prevent replication of the temperature-sensitive plasmid. Absence of the delivery plasmid and the fluorescent protein, and absence of *E. coli* contamination was checked via PCR as described in (25). The constitutive expression of mScarlet-I in colonies on agar plates and in liquid cultures was confirmed via fluorescence microscopy. Expression of the fluorescent protein did not significantly influence the overall growth of either strain at any 4MSOB-ITC concentration (Suppl. Fig. 1). The labelled strains were therefore used throughout the experiments.

### Growth curves of mono- and cocultures of bacteria

All strains were streaked from glycerol stocks on R2A agar and incubated at 28°C. Individual colonies were used to inoculate R2A broth and shaken overnight at 28°C at 200 rpm. mScarlet-I-tagged PS and PSKO strains were pre-grown with appropriate antibiotics. On the next day the cultures were normalized to OD_600_ = 0.2 (monocultures) or 0.4 (cocultures). For pairwise cocultures, equal volumes of the commensal and PS or PSKO were combined, diluting each strain to a final OD_600_ = 0.2. For pairwise cocultures with different PS(KO):commensal ratios, the total OD_600_ was kept at 0.4, and the OD_600_ of PS(KO) and a commensal partner were mixed in different ratios (PS(KO):commensal = 100:1, 10:1, 1:1, 1:10, 1:100). In a 96-well plate three replicates per 4MSOB-ITC concentration were inoculated with 10 µL normalized culture in a total volume of 100 µL supplemented with 4MSOB-ITC ranging from 0 (pure DMSO) to 60 µg/mL. Negative growth controls (blanks) were inoculated with sterile R2A broth instead of bacteria. The plate was incubated at 28°C in a TECAN Infinite M Plex plate reader (i-control 2.0 software) and the OD_600_ and red fluorescence (excitation: 540 nm, emission: 595 nm, gain: 120) were measured every 30 min after 1 min of orbital shaking.

### Test of SaxA public good effect in a commensal bacterial community

Overnight cultures of all five commensals, PS and PSKO (both tagged with mScarlet-I) were normalized to OD_600_ = 0.6 and combined in equal amounts to form synthetic communities (SynCom): all five commensals together (SynCom5), SynCom5+PS (hereafter: SynCom5+SaxA) and SynCom5+PSKO (hereafter: SynCom5-SaxA). Instead of PS or PSKO, sterile R2A broth was added to dilute SynCom5 such that in all SynComs each strain was present at abundance corresponding to an OD_600_ of 0.1. 1350 µL R2A was inoculated with 150 µL of the SynCom of interest in a 5 mL tube supplemented with a final concentration of 60 µg/mL 4MSOB-ITC, with five replicates per SynCom. The cultures were incubated on the shaker at 150 rpm at 28°C for 24 h. After 0, 3, 5, 7, 9, 11 and 24 h 150 µL of the cultures were sampled, transferred first to a 96-well plate to measure the OD_600_ and red fluorescence on the plate reader and then they were frozen at -20°C. The OD_600_ of the non-inoculated medium was monitored throughout; and, as expected, there was no increase in OD_600_ over time, so samples for DNA extraction (as sterility control) were only taken after 24 h. Samples for 4MSOB-ITC and 4MSOB-amine quantification were taken after 24 h as well, to confirm the degradation of the ITC in SynCom+SaxA. To enable absolute quantification of bacterial growth, before the DNA extraction, 2 µL of internal standard (ZymoBiomics, Spike-In Control, High Bacterial Loads, Zymo Research, Freiburg, Germany) which contains a defined cell number of the gram-negative bacterium *Imtechella halotolerans* and the gram-positive bacterium *Allobacillus halotolerans* were added to each sample, except for samples from timepoint 24 h and the inocula, which received 8.5 µL. The DNA was extracted using bead beating and a SDS-buffer protocol, as described in (20).

The V3-V4 region of the 16S rRNA gene was amplified with primers 341F/799R (26) and amplicon sequencing libraries were prepared in a two-step PCR as described in (20). Pooled libraries were sequenced on an Illumina MiSeq instrument using a MiSeq Reagent Micro Kit v2 (300-cycles). The quality of raw reads was checked and forward and reverse reads were combined using dada2 package (27). Only reads with a quality score greater than 15 and no more than 1 expected error were included. ASVs were assigned using the SILVA 16S rRNA gene database (v 138.1) (28).

The data was analyzed and plotted using the phyloseq package (29) in R (version 4.4.2). The non-inoculated medium mainly amplified the added internal standard taxa, confirming both its sterility and the successful amplification of both the internal standard taxa (Suppl. Fig. 2A). Positive controls (ZymoBIOMICS Microbial Community DNA Standard, Zymo Research, Freiburg, Germany) produced the expected results and negative controls (nuclease-free water) had less than 250 reads and were not analyzed further (Suppl. Fig. 2B). *Allobacillus halotolerans* was identified using all Bacillaceae reads, *Imtechella halotolerans* was identified using all Flavobacteriaceae reads, since neither family was part of the SynComs. Before further analysis the addition of 8.5 instead of 2 µL internal standard in the 24 h-samples was corrected by dividing the reads of the two internal standard taxa by 4.25. Though both internal standard taxa were added in similar cell counts, the gram-positive *Allobacillus halotolerans* was underrepresented (Suppl. Fig. 2A), suggesting lower efficiency of DNA extraction for gram-positive bacteria. Thus, gram-positive SynCom members (G, R) were normalized to Bacillaceae reads and gram-negative members (PS, PSKO, E, K, M) were normalized to Flavobacteriaceae reads. The normalzed results showed a clear increase in read counts over time in all three communities, reflecting growth (Suppl. Fig. 2C). The increase in read counts was highest in SynCom5+SaxA and lowest in SynCom5, in line with the OD_600_ measurements (Suppl. Fig. 3A). The resulting “absolute” abundances were plotted using ggplot2 package in R (30).

### Quantification of 4MSOB-ITC and its degradation products via LC-MS

To quantify 4MSOB-ITC, 4MSOB-amine and 4MSOB-ITC conjugated to glutathione (4MSOB-ITC-GSH) in supernatants of bacterial monocultures (n=3) or 4MSOB-ITC and 4MSOB-amine in supernatants of the SynComs after 24 h (n=5), we harvested 20 µL of the supernatant and diluted it in 180 µL MiliQ water. The diluted samples were frozen at -20°C until the analysis was carried out on an LC-MS as described (20). Briefly, external standard curves of 4MSOB-ITC (L-Sulforaphane, CAS 142825-10-3, Sigma Aldrich) and 4MSOB-amine (4-methanesulfinylbutan-1-amine, Enamine Germany GmbH, Frankfurt a.M., Germany) were measured together with the samples. 4MSOB-ITC-GSH was quantified relatively, and the data is shown in peak area units. 4MSOB-ITC, 4MSOB-amine and 4MSOB-ITC-GSH were analyzed on an Agilent 1200 HPLC system (Agilent, Santa Clara, CA, United States) coupled to an API3200 tandem mass spectrometer (AB SCIEX, Darmstadt, Germany). All compounds were separated on an Agilent XDB-C18 column (5 cm × 4.6 mm, 1.8 μm, Agilent, Waldbronn, Germany). The mobile phase consisted of 0.05% (v/v) formic acid in ultrapure water as solvent A and acetonitrile as solvent B, at a flow rate of 1.1 mL/min. The elution gradient was: 0-0.5 min, 3-15% B; 0.5-2.5 min, 15-85% B; 2.5-2.52 min, 85-100% B; 2.25-3.5 min, 100% B; 3.5-3.51 min, 100-3% B, 3.51-6 min, 3% B. The ion spray voltage was maintained at 5500 eV in positive mode. The turbo gas temperature was set to 500 °C, nebulizing gas to 60 PSi, drying gas to 60 PSi, curtain gas to 35 PSi, and collision gas to 3 PSi. Details of multiple reaction monitoring (MRM) can be found in Suppl. Tab. 2. Analyst Software 1.6 Build 3773 (AB SCIEX) was used to acquire and process the data.

### Model development

We developed a dynamical model based on ordinary differential equations to describe bacterial growth via nutrient assimilation, suppression of growth by the ITC, and degradation of the ITC by the opportunistic pathogen (Fig. 1A, 1B). In the absence of ITC, the ITC-degrading and non-degrading strains of PS are assumed to grow equally well, and nutrient utilization is modelled by decreasing nutrient concentration proportional to bacterial growth (blue curve, Fig. 1C, 1D). However, increasing the ITC concentration affects the degrader and non-degrader strains differently.

**Figure 1:**
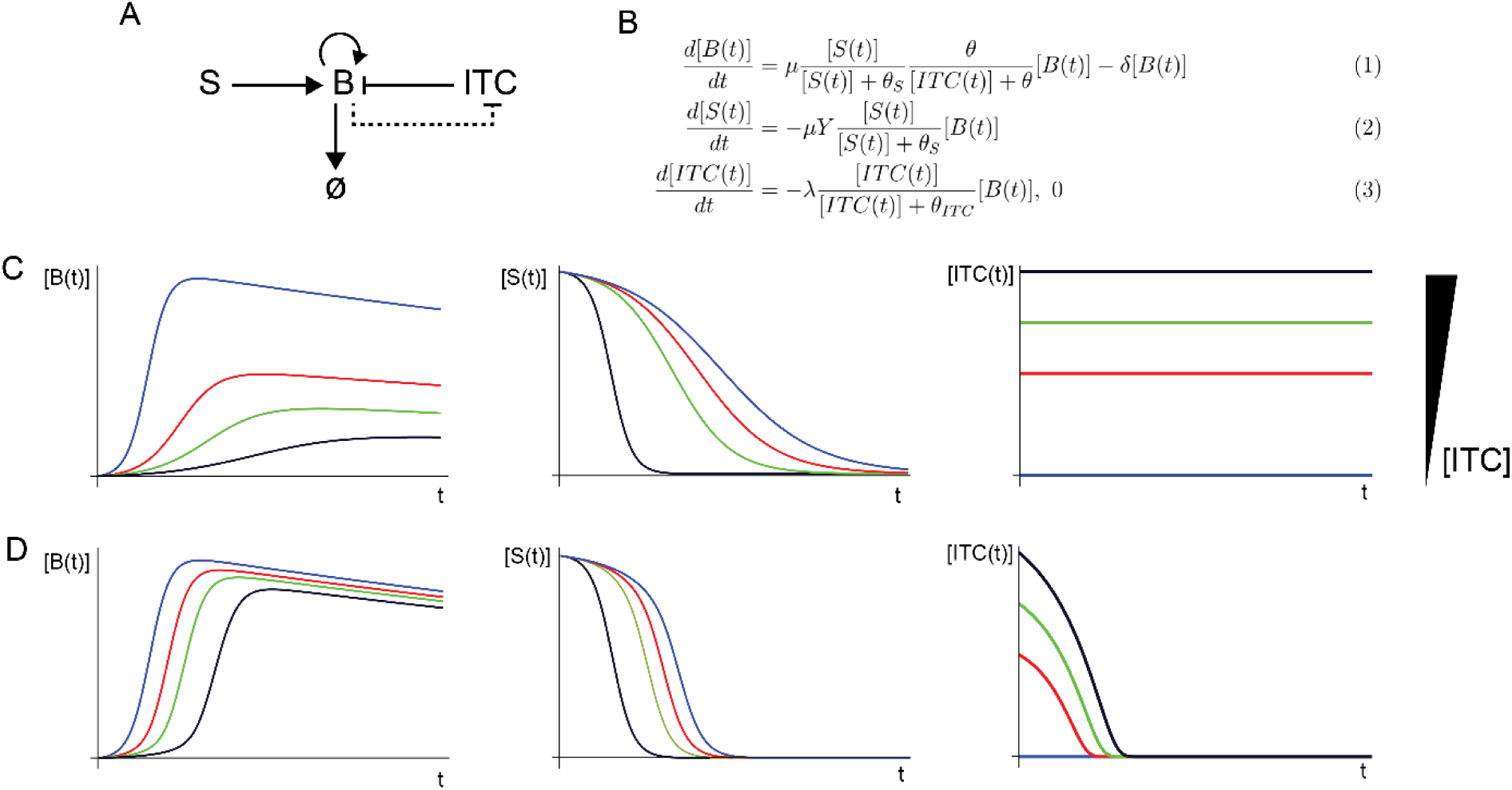
The mathematical model. **(A)** Schematic description of the model: the rate of change of bacterial biomass B is governed by a balance between growth due to consumption of the substrate S and death. ITC inhibits growth (and, effectively also decreases the biomass yield; see Suppl. Text 1), but ITC can be degraded by the degrader-strain PS (dotted line). **(B)** System of ordinary differential equations governing the growth dynamics of a given strain, nutrient consumption and ITC inhibition and degradation, in the case of a single substrate. Biomass B increases by assimilating the substrate S via a Monod term with maximal growth rate µ and half-maximal concentration θ_S_, while ITC impairs growth via a repression term with half maximal ITC concentration θ. Bacteria die with a constant death rate δ (Equation 1). The substrate S is consumed via a corresponding Monod term and transformed into biomass; the constant Y converts between the units of biomass and substrate (Equation 2). The degrader PS strain degrades ITC with maximal degradation rate λ and half-maximal ITC concentration θ_ITC_, while non-degrading strains are characterized by a null ITC degradation rate (Equation 3). These equations correspond to a single substrate version of the model; the experimental data was fitted using a slightly more complex, 2-nutrient version of the model for the PS and PSKO strains, see Suppl. Text 2 **(C, D)** Qualitative illustration of the model dynamics for the biomass concentration [B(t)], substrate concentration [S(t)], and ITC concentration [ITC(t)] for the non-degrader strain PSKO (C) and degrader strain PS (D) for a range of values of the initial ITC concentration. For these simulations, we used the equations in (B) and parameter values in Suppl. Tab. 3.

For the non-degrader strain (PSKO), ITC toxicity is modelled by slowing growth and decreasing the maximal biomass (i.e. the yield) at higher ITC concentrations (Fig. 1C). Since this strain does not degrade ITC, the ITC concentration remains constant in time (Fig. 1C). These model features are consistent with our experimental growth curves in which both the exponential growth rate and the maximal OD_600_ decrease with increasing ITC concentration.

The ITC degrader is modelled identically to the non-degrader (with the same ITC susceptibility), the only exception being that this strain degrades ITC with dynamics which were tested experimentally previously (23). Running the model for this strain, we observe that growth is delayed by the presence of ITC, until the strain later achieves a growth rate and maximal biomass that are almost the same as in the absence of any ITC – consistent with the fact that the ITC has been degraded (Fig. 1D).

The mathematical model is described, in its simplest form, by Equations 1-3 (Fig. 1B). The biomass growth rate in Equation 1 (Fig. 1B) depends on nutrient concentration via a standard Monod function (31) in which nutrient S is consumed with maximal growth rate µ and half-maximal concentration θ_S_. The Monod function is modulated by a term that describes inhibition by the ITC, with half-maximal repression concentration θ (32). A smaller value of θ corresponds to greater ITC sensitivity. Bacteria are also assumed to die at a constant rate δ. To account for the observed decrease in maximal OD_600_ with increasing ITC, the rate of nutrient consumption (Equation 2, Fig. 1B) depends on the nutrient concentration S via the same Monod function, multiplied by a yield coefficient Y (that converts units from nutrient to biomass concentration), but it does not depend explicitly on the ITC concentration. This is equivalent to having an effective growth yield that decreases with ITC concentration (Suppl. Text 1). The ITC-degrading and non-degrading strains are modelled with the same equations and parameter values, except that the degrading strain degrades the ITC with maximal rate λ and half-maximal concentration θ_ITC_ (Equation 3, Fig. 1B). The equations describing the non-degrader strains are also used to model the commensals that are unable to degrade ITC, but with parameter values derived from their experimental measurements of ITC susceptibility and growth (Suppl. Text 2). Within this model framework, the interaction between the pathogen and commensal strains is mediated only by ITC degradation and competition for nutrients. Our experimental growth curves for the ITC-degrader (PS) and non-degrader (PSKO) show an apparent change in growth dynamics (or “kink”) after 10-15 h that may correspond to utilization of a second nutrient within the complex R2A medium, i.e. a diauxic shift. To model this observed behavior we extended the model to include a diauxic shift to a second nutrient (Suppl. Text 2). This required three additional parameters describing the maximal growth on the second nutrient, a second Monod constant, and a diauxic repression coefficient (Suppl. Text 2). This resulted in an excellent fit to the data (Fig. 4, Suppl. Tab. 4)

The model results shown in Figures 4 and 6 include this diauxic shift and therefore corresponds to equations 4-7 and 8-12 of Suppl. Text 2, for monocultures and cocultures, respectively.

### Fitting the model to experimental data to obtain parameter values

To obtain the parameter values µ, θ_S_, Y, θ and δ that describe the growth curves of PS, PSKO and the commensals, the mathematical model was fitted to monoculture growth curves, assuming that the biomass density B in our model is proportional to the experimentally measured OD_600_. To perform the fit, we wrote a custom routine in MATLAB that first subtracts background (media only blank values) from the raw growth curve data and then computes averages S_observed,i_ and standard deviations σ of the experimental OD_600_ measurements at each time point i over multiple replicate growth curves. The fit is then performed by minimizing the error-weighted chi-square function

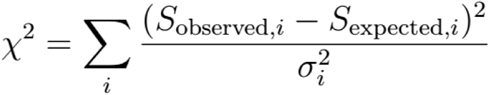

where S_observed,i_ and S_expected,i_ are the OD_600_ values, for a given time point i, as experimentally observed and as predicted by the model, respectively. The predictions for S_expected,i_ are obtained by numerical integration of the model equations for a test parameter set. The test parameter set is then systematically varied to find the parameter values which minimize the absolute difference between the model prediction and the experimental data, summed over all time points for that dataset. To achieve this, we used the MATLAB routine *fminsearch* and the system of ordinary differential equations was solved numerically via the function *ode45*. The standard deviations of the parameter values were computed by using the procedure described in (33). All the parameter values in this study were therefore obtained by fitting experimental data for monocultures of either PSP, PSKO, or commensals (Fig. 4, 5, Suppl. Tab. 4).

### Model validation

We validated the model (which had been parameterized on monoculture data) by predicting growth curves (OD_600_ as function of time) for 1:1 mixed cultures. We first verified the model by predicting a simple culture, where we mixed an equal amount of fluorescently tagged PS(KO) and non-tagged PS(KO) at different ITC concentrations and cultured them as described above (Suppl. Fig. 4A, 4B). In this case, where the two strains are described by the same equations (Suppl. Text 2) and the system differs only for the initial conditions compared to the monoculture, the model predictions were in good agreement with the data. Then, we predicted growth curves for 1:1 mixed cultures of the degrader (PS) and non-degrader (PSKO) strains with different ITC concentrations. For mixed cultures, the model outcome depends on two effects: the two strains compete for nutrients, and PS degrades the ITC, protecting PSKO (see Suppl. Text 2 for the equations corresponding to the mixed culture model). The model’s predictions were tested experimentally by measuring the dynamics of OD_600_ in mixed cultures of PS and PSKO. The model predictions for the OD_600_ of the mixed cultures were again in good agreement with the experimental data (Suppl. Fig. 4C, 4D). Moreover, the model also allows us to predict the dynamics of the PS and PSKO subpopulations within the mixed culture, revealing how PS promotes the growth of PSKO compared to the PSKO monoculture. Indeed, the model predicts that since SaxA functions as a public good, PS and PSKO grow equally well within the coculture (Suppl. Fig. 4E, 4F); however, since nutrients are shared between the two subpopulations, competition suppresses the maximal population size of each strain compared to monoculture (Suppl. Fig. 4D). The good agreement between our model predictions and measured growth curves suggests that the model, while conceptually simple, reproduces the essential biological mechanisms that are at play in our system, namely, protection of non-ITC-degraders by degraders, together with competition for nutrients.

## Results

### Commensal leaf bacteria are sensitive to 4MSOB-ITC to different extents

We screened a diverse set of commensal bacteria isolated from *A. thaliana* leaves (Suppl. Tab. 1) to find strains that may benefit from SaxA-mediated ITC degradation. First, we checked for SaxA-like activity, although to the best of our knowledge it is not described in non-pathogenic bacteria. To our surprise, 4 out of 13 tested commensal taxa degraded 4MSOB-ITC to 4MSOB-amine, which is the typical degradation product of the ITC hydrolase SaxA (Fig. 2C, Suppl. Fig. 5A). From the remaining nine taxa, we selected five (*Stenotrophomonas* sp. E, *Plantibacter* sp. G, *Janthinobacterium* sp. K, *Brevundimonas* sp. M and *Rhodococcus* sp. R (Fig. 2B)) that are phylogenetically diverse and represent the major leaf colonizing phyla Alphaproteobacteria, Gammaproteobacteria, and Actinobacteria (Suppl. Tab. 1). They all grew to detectable levels within 24-48 h in R2A medium and showed variable susceptibility to 4MSOB-ITC. Among the selected five commensals *Stenotrophomonas* sp. E was the most ITC-resistant, while the least resistant strain was *Janthinobacterium* sp. K (Fig. 2B, Suppl. Fig. 5B). To control for SaxA-independent interactions between PS and the commensals we used a *saxA* knock-out mutant (PSKO). As expected, PSKO was more sensitive to high concentrations of the ITC than the wildtype PS (Fig. 2A, 2C). Previous quantification of the kinetics of 4MSOB-ITC degradation by PS has shown that the dose of ITC supplied is fully degraded in 3-6 h under the same culture conditions (23).

**Figure 2:**
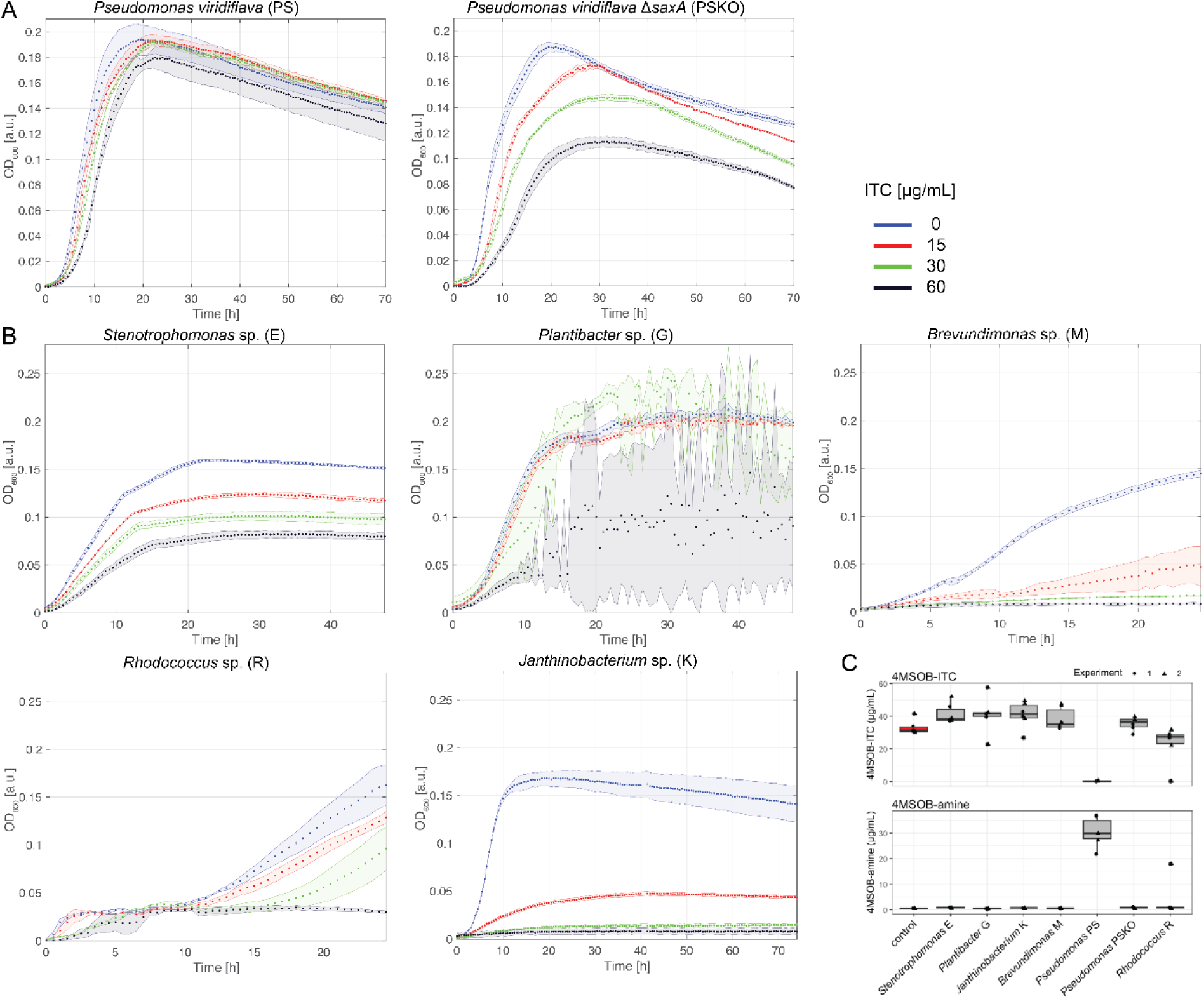
Leaf bacteria are sensitive to 4MSOB-ITC to different extents. Growth curves of PS and PSKO (A) and of the five selected commensals (B) in R2A broth supplemented with different 4MSOB-ITC concentrations at 28°C. Dots represent means, shaded areas depict standard deviations of one experiment with n=3 technical replicates. With higher ITC concentrations, *Plantibacter* sp. G formed cell aggregates resulting in high standard deviations. (C) Quantification of 4MSOB-ITC and its breakdown product 4MSOB-amine in the supernatant of monocultures of the five commensals, PS and PSKO after 16 h of incubation. The different symbols illustrate two independent experiments (n=3 each). Boxplots summarize the data of both experiments. Red boxplots show controls which were not inoculated with bacteria, grey boxplots show data for the bacterial strains.

### SaxA-mediated rescue of diverse commensals in a community is dependent on their ITC sensitivity

Previously, we had shown that the commensal G benefits in an ITC-concentration dependent manner from SaxA-mediated ITC degradation by PS (23). To assess the effect of SaxA within a community, we performed growth experiments with a synthetic community (SynCom) containing all five of our selected ITC-sensitive commensals (SynCom5), together with PS (SynCom5+SaxA) or with PSKO (SynCom5-SaxA) (Fig. 3A). The liquid medium was supplemented with a high 4MSOB-ITC concentration (60 µg/mL) which is sufficient to strongly inhibit the growth of most commensals (Fig. 2A). We took samples for 16S rRNA gene amplicon sequencing at regular intervals and at the end of the experiment. ITC quantification after 24 h confirmed its degradation only in SynCom5+SaxA (Suppl. Fig. 3B). The absolute microbial abundance, as measured by both normalized amplicon sequencing data and OD_600_, was, as expected, highest for the SynCom5+SaxA community. However, perhaps surprisingly, SynCom5 without PS or PSKO also grew significantly at this high ITC concentration (Suppl. Fig. 2A, Suppl. Fig. 3A). The most ITC-resistant commensal strain E dominated all three communities, regardless of the presence of *Pseudomonas* or SaxA (Fig. 3B, Suppl. Fig. 6). Commensals M, G and especially K benefited from SaxA, showing growth in SynCom+SaxA but not in SynCom-SaxA after 24 h, commensal R did not grow in the community setting within 24 h (Fig. 3B, Suppl. Fig. 6). Thus, in the presence of high levels of ITC, SaxA is a public good that can simultaneously benefit diverse sensitive commensals.

**Figure 3:**
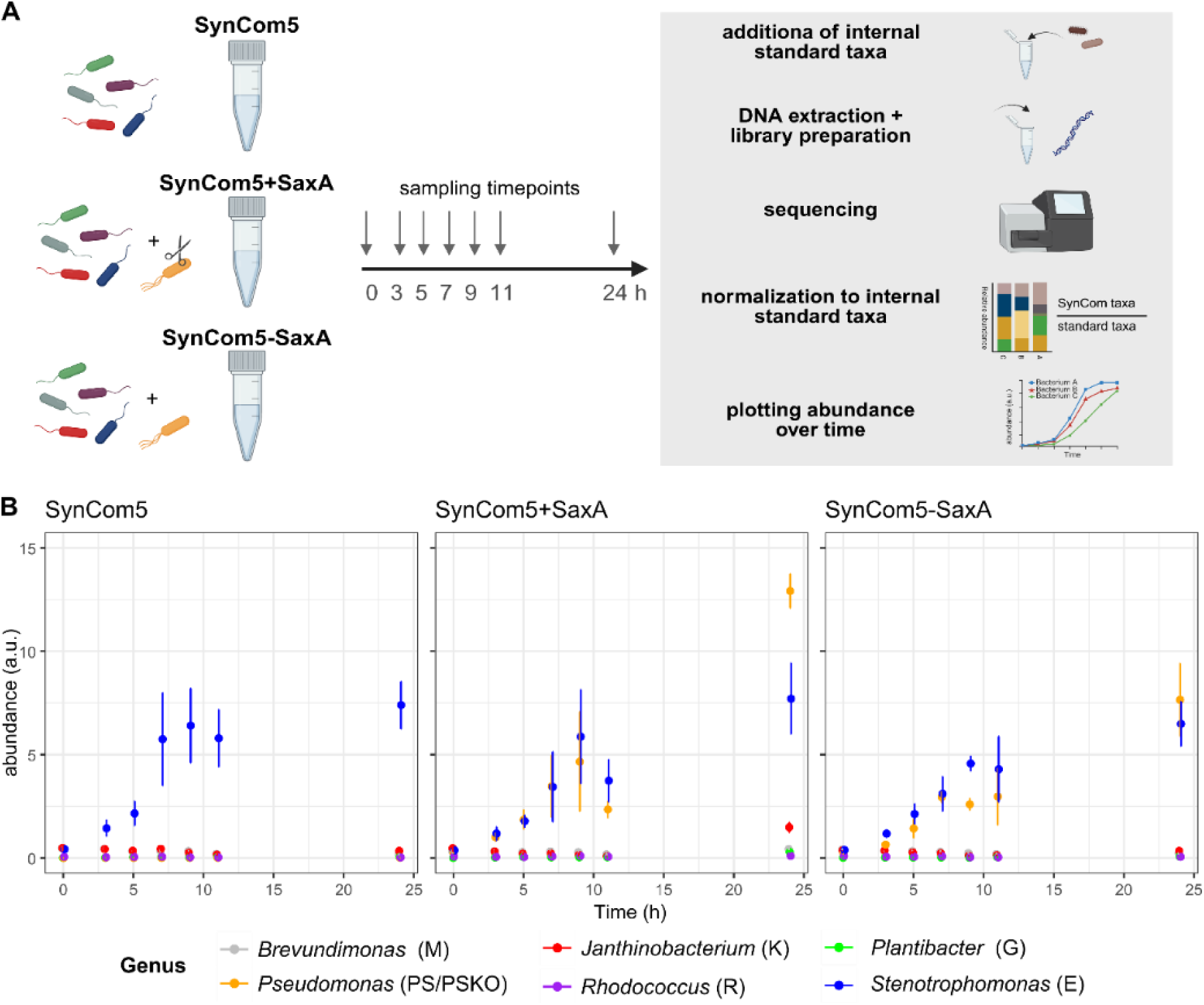
Growth of commensals in a synthetic community with or without SaxA. **(A)** Experimental design: all five commensals were mixed in equal amounts (SynCom5) and either PS (SynCom5+SaxA) or PSKO (SynCom5-SaxA) were added. The communities were inoculated in R2A broth supplemented with 60 µg/mL 4MSOB-ITC (n=5 technical replicates) and sampled over 24 h. Internal standard taxa were added before DNA extraction and subsequent library preparation. After sequencing the gram-positive SynCom taxa were normalized to the gram-positive standard taxa and the same for gram-negative ones. Created with Biorender.com. **(B)** Normalized absolute abundances (a.u. = artificial units) of each SynCom taxon over the course of 24 h. Each dot represents the mean of five replicates; standard deviations are depicted as whiskers.

### Commensal and pathogen growth can be quantitatively described by fitting model parameters to monoculture growth curves

To dissect the microbial interactions responsible for the observed public good function of SaxA in the SynCom, we developed a mathematical model, which we parameterized using growth data for monocultures of the five commensals, PS, and PSKO. The model consists of a set of ordinary differential equations describing bacterial growth, inhibition by ITC and degradation of ITC, and it makes qualitatively distinct predictions for the growth dynamics of ITC-degrading and non-degrading strains, in the presence of ITC (Fig. 1; see Methods). Model parameters for PS and PSKO are estimated by fitting to monoculture growth curves for a range of ITC concentrations (Fig. 4), taking into account a diauxic shift in nutrient usage on the complex R2A medium (see Methods, Suppl. Fig. 7, Suppl. Text 2); parameter values are assumed to be identical for PS and PSKO apart from the ITC degradation rate (λ in the model; see Fig. 1), which is zero for PSKO. The model produces an excellent fit to our monoculture data for PS and PSKO across the full range of ITC concentrations used in this study (Fig. 4, Suppl. Tab. 4).

**Figure 4:**
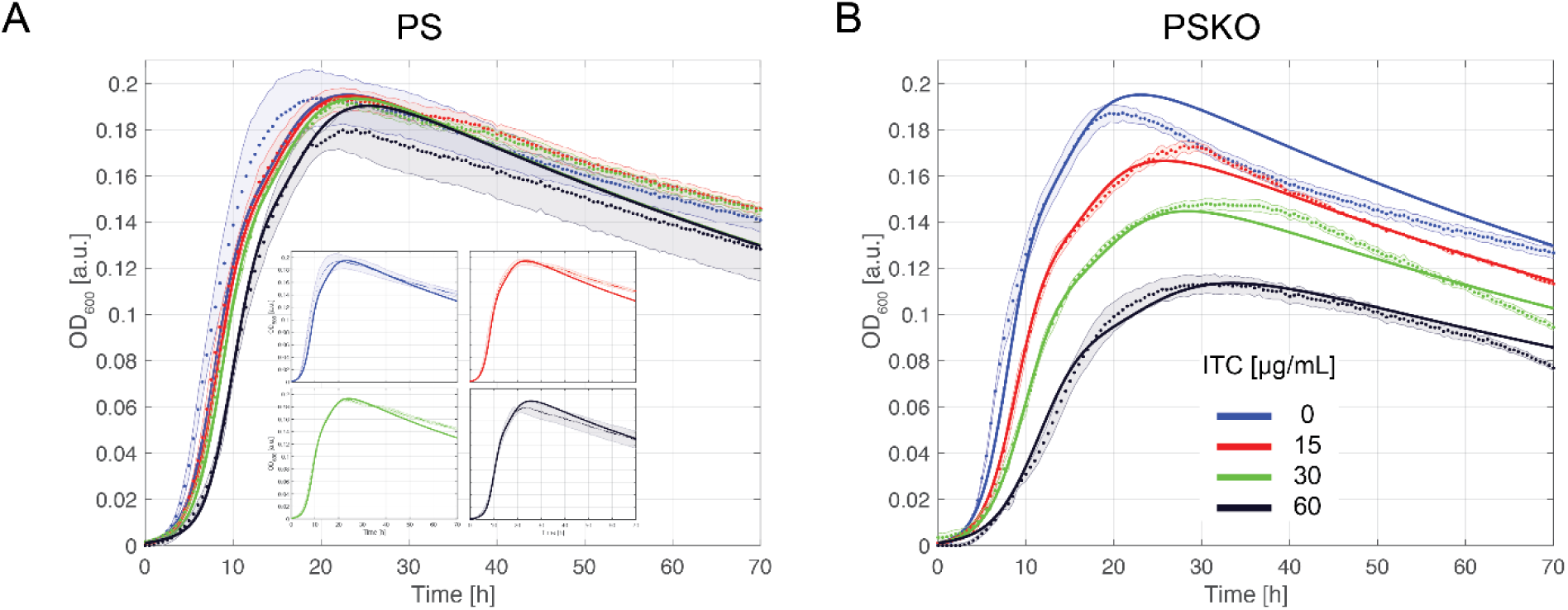
Fits of PS and PSKO monocultures. Fits of the mathematical model (including diauxic shift) to the experimental growth curves for PS **(A)** and PSKO **(B)** monocultures at increasing ITC concentrations. Experimental data points (dots) represent the mean of three technical replicates, with standard deviations indicated by shaded regions. Model predictions are depicted by solid lines. For clarity, small panels in A show separate fits for the different ITC concentrations of the PS monoculture.

We next used the same model, with some modifications, to describe the growth of each of the five commensal strains in monoculture, over a range of ITC concentrations. These strains were shown not to degrade ITC. The model parameters for each commensal strain were obtained by fitting to the respective monoculture growth curve. We tested whether including a diauxic shift (i.e. a two-nutrient model, as we used for PS and PSKO) improved the fit, but this was the case only for R (Fig. 5E, Suppl. Fig. 8); therefore, we used a single-nutrient model for all the commensals. For strains K and G, the growth rate depended non-linearly on ITC concentration (Figs. 5A, 5C); therefore we included a cooperative (Hill-type) inhibition function for these two commensals. We also noticed by visual inspection of our experiments that G and R showed biofilm and aggregate formation in the microplate wells, especially at later timepoints and for higher ITC concentrations (Fig. 5C, 5E; Suppl. Fig. 9). Since this likely interferes with the OD_600_ measurements, the ability of our model to predict the growth dynamics of these strains may be limited, especially for times later than 24 h. The full model used for each commensal strain, together with model fits and parameters are described in the Suppl. Text 2 and Suppl. Tab. 4. Thus, the simple model formulation of Fig. 1, with minor modifications, can describe monoculture growth of a range of commensals, with the exception of biofilm/aggregate formation. For the remainder of this study, we will focus on the most and the least ITC-sensitive strains, K and E, respectively (Fig. 5A, 5B).

**Figure 5:**
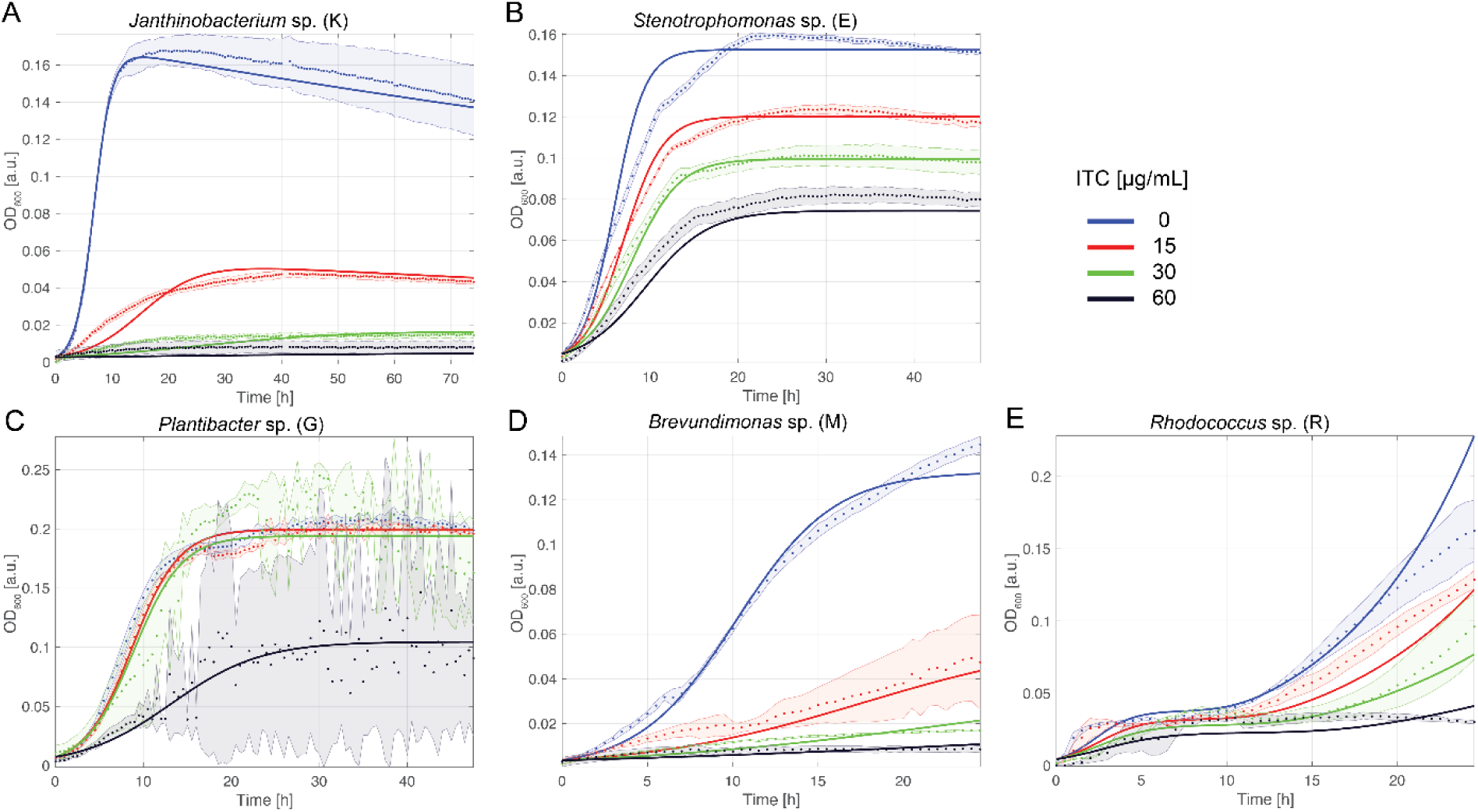
Model fits to monoculture growth data for the commensals. The single-nutrient model fitted to monoculture growth data for K **(A),** E **(B),** G **(C),** and M **(D)**, but a model with diauxic shift fitted better for R **(E)**. Parameter values resulting from these fits are given in Suppl. Tab. 4. Different ITC concentrations are represented by distinct colors. The data points and shaded areas denote the mean and standard deviation of the experimental data with three technical replicates while the solid lines represent the model fits.

### ITC detoxification and nutrient competition control the growth dynamics of pairwise cocultures

Next, we used the fitted model to predict SaxA-mediated interactions. Specifically, we predicted the growth of pairwise cocultures of commensals with either PS or PSKO, at different ITC concentrations. The model accounts for nutrient competition (that limits the total carrying capacity of the system) as well as SaxA-mediated ITC degradation; by comparing model predictions to coculture data we aimed to assess whether these two mechanisms could account for the observed coculture data.

Model predictions for the total OD_600_ of cocultures of PS/PSKO with either commensal K or E showed good agreement with our experimental data (Fig. 6A). The model also predicted that PS/PSKO is dominant while the commensal contributes up to one third of the total biomass (Fig. 6B). Comparing predictions for the most and least ITC-sensitive strains, the model shows that the more ITC-sensitive strain K benefits more from SaxA, whereas the more resistant strain E shows only a moderate increase in growth in the presence of PS versus PSKO, especially at low ITC levels. Despite the above-mentioned biofilm/aggregate formation by strains G and R (Suppl. Fig. 9), the model predictions for both mixed communities were in good agreement with the data, probably because of the dominant contribution of PS/PSKO to the biomass (Suppl. Fig. 10).

**Figure 6:**
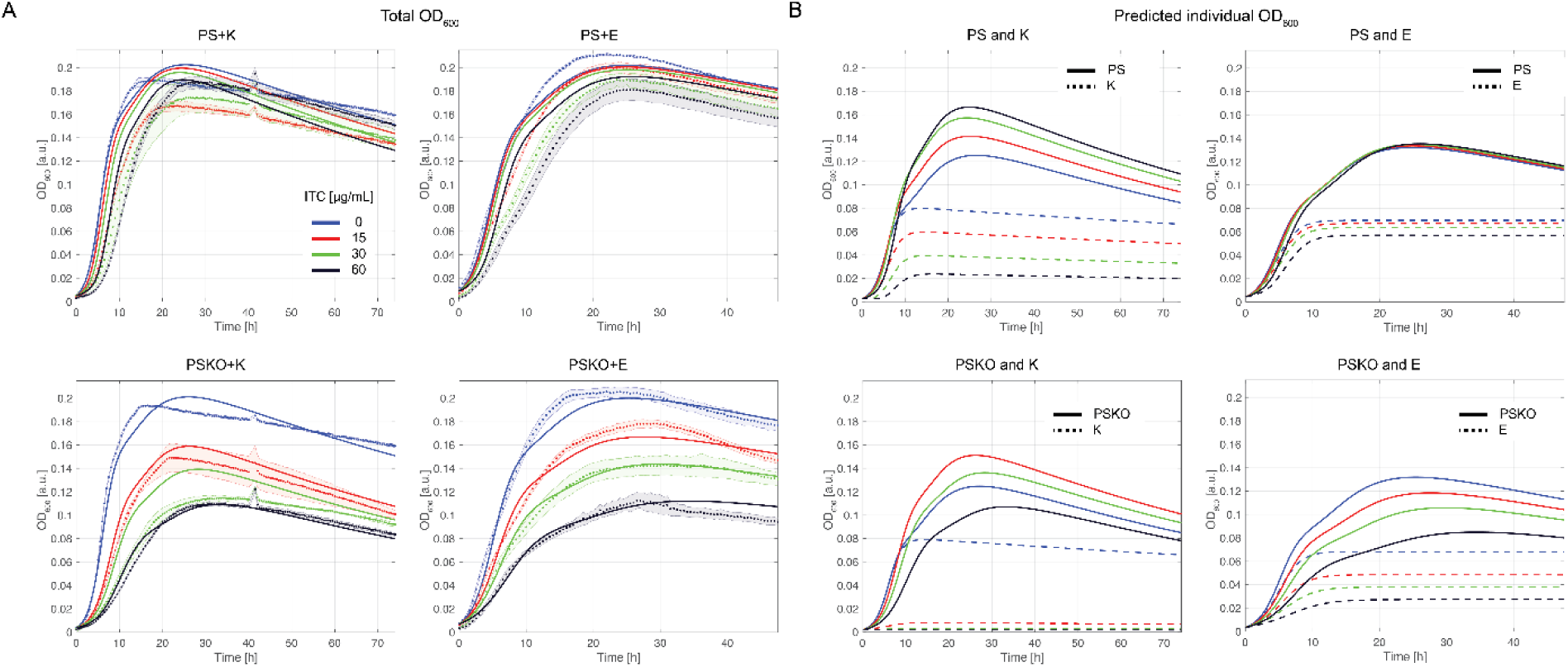
Model predictions for bacterial biomass in pairwise cocultures of PS or PSKO with commensal K or E. (A) The model prediction for the total biomass dynamics (OD_600_) of the pairwise coculture starting with the same amount of either PS or PSKO and the commensal for different ITC concentrations are shown by the solid lines and compared to the experimental data (dots, means of three technical replicates, standard deviations as shaded region). (B) Predicted separated growth dynamics of the biomass of either PS or PSKO and the commensal K or E in pairwise cocultures. Solid lines show the predicted OD_600_ of PS or PSKO, and dashed lines the OD_600_ of the commensal strain.

Thus, our mathematical model, that includes only nutrient competition and ITC inhibition and degradation, successfully predicts the growth dynamics of pairwise cocultures of commensals together with either PS or PSKO, and mixed communities. These results indicate that SaxA functions as a public good independently of the interaction partner, while also revealing that the rescued strain competes with the ITC-degrading strain for limiting nutrients. In coculture, nutrient resources are shared between the PS/PSKO and the commensal strain; consequently, a higher abundance of commensals restricts PS/PSKO growth, and vice versa. This reciprocal limitation is evident when comparing the maximal OD_600_ of the red curves representing PS and PSKO in Fig. 6B. At an intermediate ITC concentration (15 µg/mL), which only mildly inhibits PSKO growth and has no measurable effect on PS, the maximal OD_600_ of PS is further reduced in the presence of the rescued commensal strain K, demonstrating competitive interactions under nutrient limitation.

### A rescue index and a pathogen suppression index illustrate the trade-offs involved in SaxA-mediated biocontrol of the opportunistic pathogen

Up to now, we considered scenarios in which the opportunistic pathogen PS and the commensal strain were initially in equal abundance (Fig. 6B). However, in the context of plant leaf colonization, the abundance ratio between opportunistic pathogen may vary greatly: in a diseased plant an opportunistic pathogen may outnumber other colonizers while in a healthy plant it may be outnumbered by commensals. Additionally, ITC concentrations vary over a wide range, from low levels in healthy tissue to high levels immediately after tissue damage. To understand the trade-off between ITC degradation by the opportunistic pathogen, which benefits commensals, and competition for resources between the opportunistic pathogen and the commensals, which may limit pathogen abundance, we used our model to predict how varying both the ITC concentration and the initial ratio of PS(KO):K influences the growth of opportunistic pathogen and commensal in coculture.

Our model predicts that ITC degradation and nutrient competition shape pathogen-commensal cocultures. At 15 µg/mL ITC, the outcome for the ITC-sensitive commensal K depends on the initial mixing ratio (red lines, Fig. 7A; Suppl. Fig. 11). If PS is at least as abundant as K (1:1 or 100:1), it monopolizes nutrients and suppresses K. If PS is much rarer (1:100), it still degrades ITC, allowing K to grow, which in turn restricts PS through nutrient competition (Fig. 7A, Suppl. Fig. 11).

**Figure 7:**
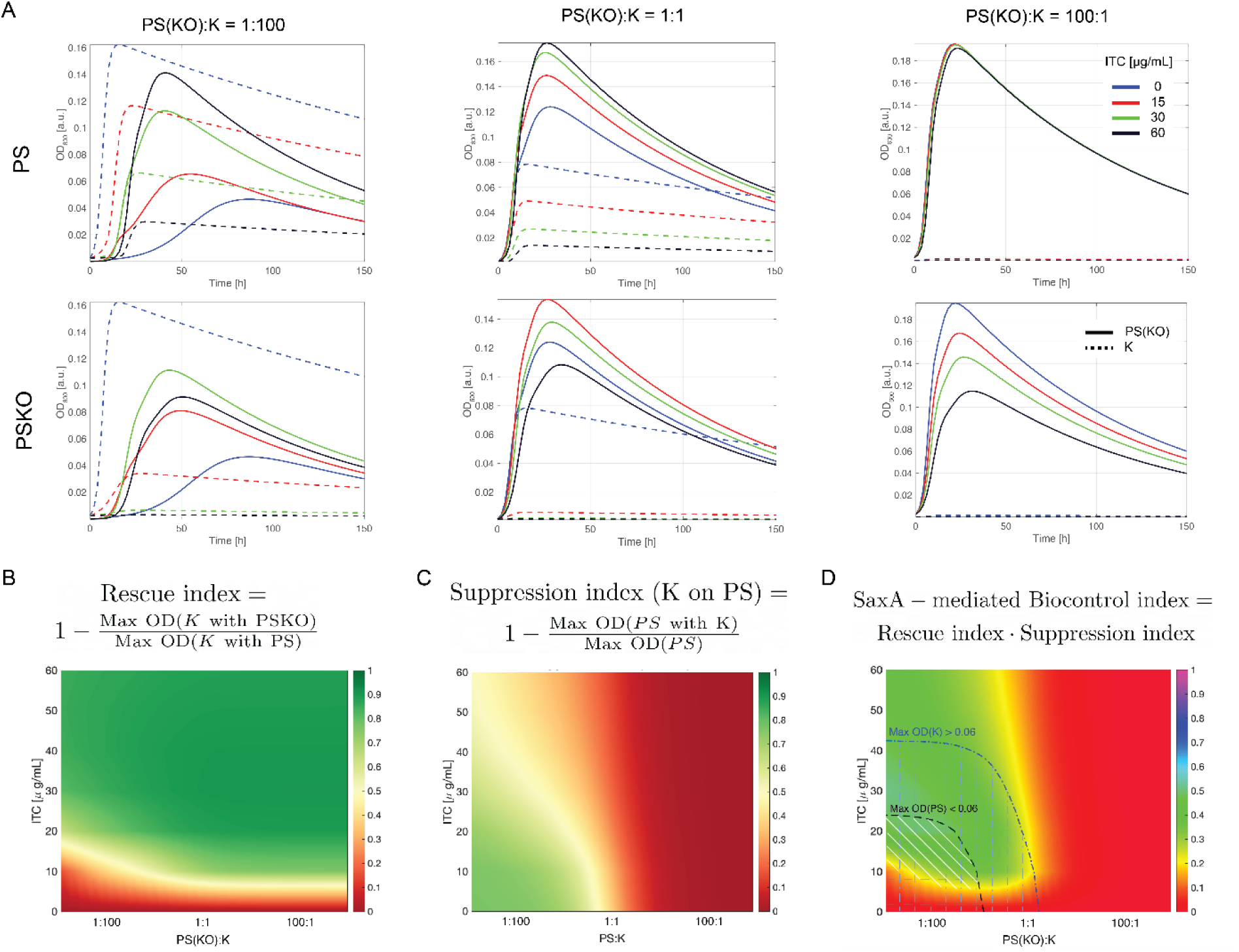
Three indexes to quantify the trade-off between commensal rescue and pathogen growth control. **(A)** Predicted growth dynamics of OD_600_ of PS and PSKO together with K at different initial ratios and ITC concentrations. Solid and dashed lines show the OD_600_ dynamics of the opportunistic pathogen and the commensal, respectively. **(B-D)** All three indices summarize the growth dynamics of PS(KO) and commensals at different ITC concentrations and PS(KO):commensal ratios. **(B)** The rescue index quantifies to which extent SaxA rescues growth of a commensal (in this case K) from ITC inhibition, detailed description in Suppl. Text 3. It compares the maximal OD_600_ (shown in heatmaps in Suppl. Fig. 12) of the commensal strain with and without SaxA-mediated ITC degradation and ranges from 0 (no rescue) to 1 (complete rescue). **(C)** The suppression index quantifies the extent to which growth of the commensal reduces the growth of PS. It compares the maximal OD_600_ of PS in the presence and absence of the commensal strain (in this case K; see also the heatmaps in Suppl. Fig. 12) and ranges from 0 (minimal suppression of PS) to 1 (maximal suppression of PS). **(D)** The SaxA-dependent biocontrol index combines the rescue index and the suppression index. The biocontrol index ranges from 0 (low K rescue and low PS suppression) to 1 (high K rescue and high PS suppression). This index is based on relative growth comparisons between PS(KO) and the commensal. To allow to compare to absolute abundances of the bacterial strains we introduce a black dashed line and blue dot-dashed line illustrating an absolute OD_600_ of 0.06 for both PS (black) and K (blue). The white hatched region defines the region where the SaxA-mediated biocontrol is maximal, PS cells are not too abundant, and K cells are present at the same number (arbitrarily set to OD_600_ = 0.06).

To characterize this complex and non-linear role of SaxA-mediated ITC degradation across different conditions (evident for instance from the maximal OD_600_ of the PSKO, solid lines in Fig. 7A, for PSKO:K = 1:100 and 1:1 that are not ordered following the increasing ITC concentrations), we therefore defined three indices to summarize the main features of the growth dynamics.

First, we defined a “rescue index” that quantifies the extent to which the commensal is rescued by SaxA from ITC inhibition (Fig. 7B, detailed description in Suppl. Text 3), ranging from 0 (minimal rescue) to 1 (maximal rescue). As expected, the rescue index is larger for high ITC concentrations where SaxA-mediated ITC degradation is more relevant (Fig. 7B) but is almost independent from the mixing ratio of PS:K, since even a small number of PS can fully rescue the commensal.

Next, we also summarized the effect of suppression of PS growth via nutrient competition with the commensal by defining a “suppression index” that quantifies the extent to which the growth of the commensal reduces PS growth (Fig. 7C), a number ranging from 0 (minimal suppression of PS) to 1 (maximal suppression of PS).The suppression index strongly depends on the initial PS:K ratio. When PS is initially present in excess (PS:K = 100:1), the commensal is not able to suppress the fast-growing opportunistic pathogen (Fig. 7C, red region, suppression index is low). However, if the initial ratio of PS is low and the ITC concentration is also low, nutrient competition becomes significant and PS is suppressed by K (Fig. 7C, green region, suppression index is high). Interestingly, the suppression index and the rescue index depend qualitatively differently on the conditions: maximal rescue of the commensal by PS occurs where the ITC concentration is high and PS:K is low (Fig. 7B); this region corresponds with intermediate values of the suppression index, where the growth of PS is only partially suppressed by nutrient competition with K (Fig 7C).

To capture this trade-off between SaxA-mediated rescue of the commensal and pathogen suppression via nutrient competition, we define a “SaxA-mediated biocontrol index” (Fig. 7C), that expresses the combined effect of the rescue and the suppression indexes. The SaxA-mediated biocontrol is maximal for low initial PS abundance and high ITC, depicted by the green region in Fig. 7D showing the intersection of the highest SaxA-mediated rescue and pathogen suppression (i.e. the green regions of Fig. 7B, 7C). Under these conditions, SaxA-mediated ITC degradation by PS is predicted to rescue commensal growth while at the same time PS growth is limited by commensals.

So far, the SaxA-mediated biocontrol index illustrates only relative changes of pathogen and commensal growth. However, for a healthy plant host the absolute numbers of pathogens and commensals matter. To illustrate that a high SaxA-mediated biocontrol index does not always correspond to low or high absolute pathogen or commensal abundances, we further refined the predicted regions of SaxA-mediated biocontrol by applying threshold criteria. For effective biocontrol, the absolute abundance of the pathogen must remain below a threshold value, while the absolute abundance of the commensal must remain above a threshold. In general, the two thresholds may have different values and need to be determined experimentally in future work; here, for illustration, we arbitrarily set them to OD_600_ = 0.06, producing the hatched blue and black regions in Fig. 7D. The white hatched region indicates where PS remains below and the commensal K exceeds this arbitrary value (see also heatmaps in Suppl. Fig. 12 and Suppl. Text 3).

### SaxA-mediated negative effect on PS growth is confirmed by experimental data of PS and K cocultures

We tested the model predictions by measuring the growth dynamics of PS(KO):K coculture at different mixing ratios (1:1, 1:10, 10:1, 1:100, and 100:1). We found excellent agreement between the model predictions and the measurements over the full range of ITC concentrations (Fig. 8), suggesting that nutrient competition and ITC degradation are indeed the key drivers of coculture dynamics.

**Figure 8:**
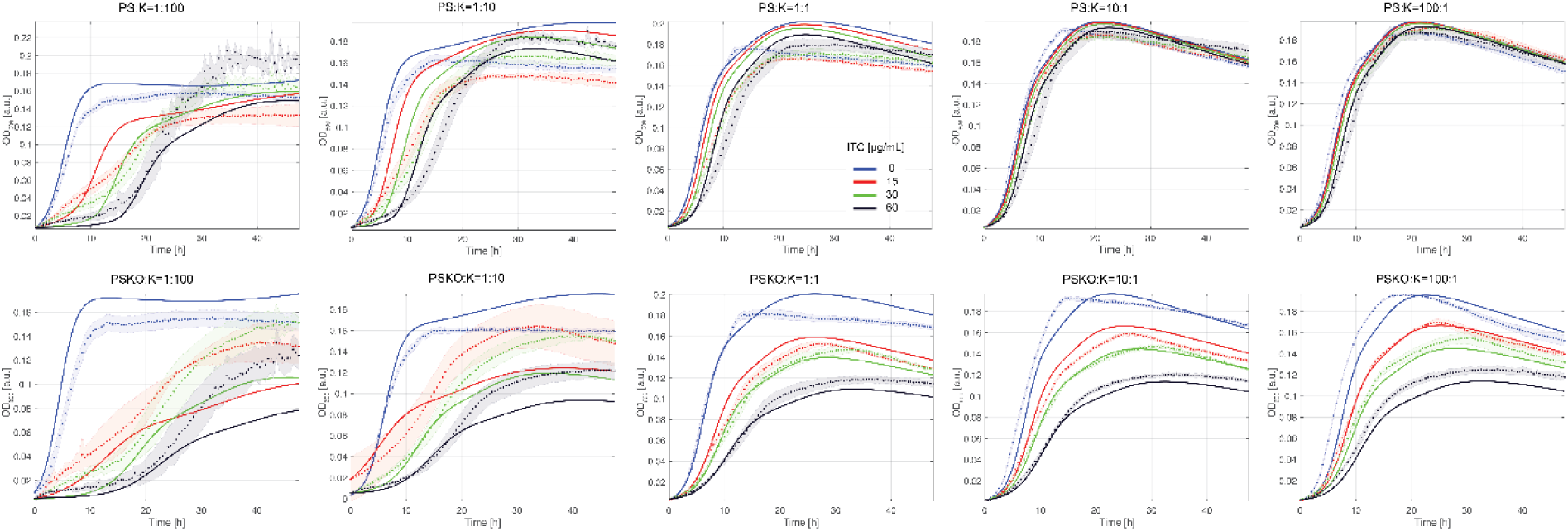
Comparison between model prediction and experimental data of pPS(KO):K at different initial ratios and ITC concentrations. Model predictions (solid lines) and experimental measurements (average as dots and shaded regions as standard deviations of three independent replicates) for the OD_600_ of the pairwise cultures PS:K and PSKO:K, in the first and second row, respectively, and for different PS(KO):K ratios (1:100, 1:10, 1:1, 10:1, 100:1) at a range of ITC concentrations.

**Figure 9:**
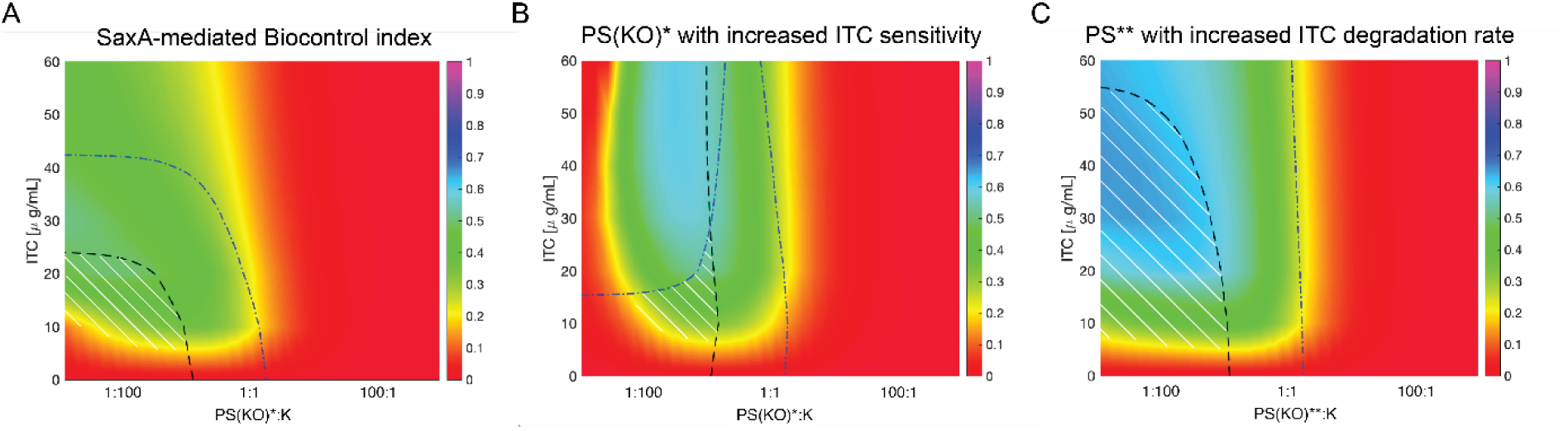
Biocontrol index maps predicted for a hypothetical pathogens PS* and or PS** compared to PS. **(A)** SaxA-dependent biocontrol for natural PS(KO) and K as shown in Fig. 7. **(B)** SaxA-mediated biocontrol index for hypothetical pathogen PS* with tenfold higher ITC sensitivity (tenfold smaller value of θ in the mathematical model, Equation 8 in Suppl. Text 2). **(C)** SaxA-mediated biocontrol index for PS** with a tenfold increased ITC degradation rate (tenfold larger value of λ in the mathematical model, Equation 12 of Suppl. Text 2). Red regions depict potential negative effects for the plant host (no rescue of commensals, no suppression of the pathogen), green/blue regions show potential positive effects which are mediated by SaxA (rescue of commensals, suppression of the pathogen). The black dashed and blue dot-dashed lines represent where the maximal OD_600_ of modified PS and K are at 0.06. The white hatched area defines the region where the SaxA-mediated biocontrol is maximal and satisfies the condition where max OD_600_(PS) < 0.06 and max OD_600_(K) > 0.06.

Our model predicted that the virulence factor SaxA would not just benefit the pathogen PS but at high ITC concentrations and high abundance of K, the ITC-sensitive commensal would indirectly benefit as well. This becomes clear as well when comparing the total OD_600_ (growth of both strains together, Suppl. Fig. 13A) to dynamics of red fluorescence (approximation for the growth of fluorescently tagged PS(KO), Suppl. Fig. 13B). Especially in cases with high ITC concentrations (30 and 60 µg/mL) the total OD_600_ clearly depends on SaxA, as it was roughly twice as high in PS:K vs. PSKO:K (Suppl. Fig. 13A). However, the pathogen’s red fluorescence was similar between PS and PSKO (Suppl. Fig. 13B), indicating that the increase in OD_600_ is likely mostly due to the rescue of K as predicted by the rescue index.

### Commensal ITC sensitivity influences the SaxA-mediated biocontrol effect on PS

To understand how the interplay between commensal rescue and nutrient competition functions for different commensals, we repeated our analysis for the least ITC-sensitive commensal strain, *Stenotrophomonas* sp. E. However, because E is only weakly ITC-sensitive (Fig. 1B), its rescue by PS is predicted to be mostly negligible. Thus, although nutrient competition with E suppresses the growth of PS when it is more abundant, SaxA-dependent biocontrol mostly does not occur because E does not need to be rescued to achieve substantial growth rates (Suppl. Fig. 14, Suppl. Fig. 15). Like for K, we experimentally tested the model predictions of PS(KO):E coculture at different ratios and ITC concentrations and find good agreement between the model predictions and the experimental data (Suppl. Fig. 16). Thus, our analysis using rescue, suppression and biocontrol indices shows that the “virulence” factor SaxA can indirectly benefit ITC-sensitive commensals depending on their ITC sensitivity.

### Pathogen and commensal traits shape the outcome of SaxA-mediated ITC degradation

To assess how trait variation influences the virulence role of SaxA, we modified model parameters *in silico* to predict outcomes of SaxA-mediated ITC degradation in different systems (Fig. 8; Suppl. Figs. 17–25). PS displays strong ITC-degrading capacity and low ITC sensitivity (Suppl. Tab. 4). We modeled a variant, PS(KO)*, with tenfold higher ITC sensitivity (θ is tenfold lower, Equation 1, Fig. 1B). Although K suppresses PS* more effectively than PS at high ITC concentrations, the SaxA-dependent biocontrol region is narrower (Fig. 8B vs. 8A) because rescue of K is impaired at low PS*:K ratios, since a larger PS* population is required for ITC degradation (Suppl. Fig. 17B vs. 17A). Conversely, increasing PS resistance by lowering θ tenfold does not change rescue, suppression, or biocontrol patterns (Suppl. Fig. 18; Fig. 7), suggesting a limit to the benefits of ITC resistance for PS and K.

We next examined how ITC-degradation capacity of the pathogen (*λ*, Equation. 3, Fig. 1B) changes the dynamics in the system. Surprisingly, a variant PS** with tenfold higher *λ* (Fig. 8C; Suppl. Fig. 17C) expands the SaxA-dependent biocontrol region, consistent with higher pathogen–commensal competition. Conversely, lowering *λ* contracts the SaxA-dependent biocontrol region (Suppl. Fig. 19).

Cocultures with a commensal K* less sensitive to ITC resembled strain E, showing minimal rescue benefit (Suppl. Fig. 25), while a more ITC-sensitive K* suppressed PS less efficiently (Suppl. Fig. 24). Thus, an optimal balance of ITC sensitivity and nutrient competition appears critical for commensal efficacy in biocontrol. Finally, modifying intrinsic growth rates of both the pathogen and commensal (*μ*, Equation 1, Fig. 1B) also altered the outcomes of the system. Increasing commensal growth or decreasing pathogen growth (twofold changes) expanded the biocontrol region (Suppl. Figs. 22, 21), whereas the opposite shifts contracted it (Suppl. Figs. 23, 20). Together, these simulations provide quantitative predictions for conditions favoring SaxA-mediated commensal rescue and pathogen suppression, highlighting potential strategies for biocontrol in bacterial leaf communities.

## Discussion

Growth and virulence of opportunistic pathogens like *Pseudomonas viridiflava* are influenced by interactions with other microbes on the plant leaf, as well as by features of the leaf environment such as plant metabolites. In turn, the colonization success of commensal bacteria on the leaf may depend on pathogens like *Pseudomonas*. Here, we showed that the toxicity of plant specialized metabolites such as GLS-derived ITCs influences the outcome of pathogen-commensal interactions. Our results demonstrate that in a bacterial community, SaxA-mediated degradation of 4MSOB-ITC can benefit growth of both *Pseudomonas viridiflava* and diverse ITC-susceptible commensals. Thus, SaxA can be considered a public good because it rescues diverse commensals from ITC toxicity. In turn, the rescued commensal strains can potentially reduce the growth of opportunistic pathogens like *Pseudomonas viridiflava* by competing for resources. The balance of this tradeoff depends strongly on factors like the traits of the competing microbes, their relative abundances, and ITC levels. In the plant context, the fact that increasing toxin levels might reduce pathogen growth and virulence while at the same time having negative effects on plant health by reducing competition with pathogen-suppressing commensals, indicates that plant defense strategies should be reevaluated from a broader perspective. There is also a need to rethink the definition and roles of virulence factors, especially those which provide a public benefit to microbial communities.

### Detoxification followed by nutrient competition shapes growth dynamics in toxin-laden environments

GLS-derived breakdown products such as 4MSOB-ITC are important barriers to leaf colonization by non-adapted pathogens and commensals (17,20,35). We previously showed that *Pseudomonas viridiflava* 3D9 (in this study: PS) can rescue the commensal *Plantibacter* sp. 2H11-2 (in this study: G) in an ITC-concentration-dependent manner (23). This is similar to other known microbial public good effects such as antibiotic degradation (36–38). In the previously studied dual system of PS and G we found an ITC-dependent benefit for G from SaxA. Especially at high ITC concentrations (60 µg/mL) growth of G was fully dependent on ITC degradation by PS. In the present study, we included four more commensals. These commensals are sensitive to 4MSOB-ITC to different extents, but for most of them, growth as well depended on ITC degradation by PS in both coculture and a synthetic community context. In turn, we find that PS growth is regulated by nutrient competition with the commensal. We used mathematical modelling to quantitatively interpret our data and to explore its potential ecological implications. In most cases, a simple mathematical model based only on growth, nutrient depletion, ITC suppression and cell death was sufficient to quantitatively explain the observed dynamics of PS-commensal cocultures. Thus, we conclude that these simple interactions dominate in the *in vitro* environment. Moreover, antimicrobial substances are found in many, maybe even most environments. A similar public goods effect was reported in toxic metal working fluids, where the growth of co-cultured bacteria was facilitated by detoxification of toxic metals. Interestingly, in this system competition increased and facilitation decreased as more nutrients were added (39). Taken together, in microbial communities with limited resources and antimicrobial substances, detoxification as a public good generally is expected to give rise to competition. This may have important implications for microbial dynamics in environments like plant leaf surfaces and apoplasts with their diverse specialized metabolites and low nutrient levels. Though our model covered the major dynamics of bacterial interactions in our *in vitro* system, we cannot rule out that the ITC breakdown product 4MSOB-amine may additionally serve as nitrogen source in nutrient-limited conditions like the leaf.

### The tradeoff between SaxA as virulence factor and public good depends on ITC, pathogen and commensal traits

Previously, we had demonstrated an ITC concentration-dependent public good effect of SaxA for the commensal *Plantibacter* sp. G (23). Here, we extend this framework and explore the role of microbial traits such as ITC degradation rate, growth rate and ITC sensitivity in SaxA-mediated microbial interactions. By altering the parameters used to model commensals and PS *in silico* we showed that the outcome of pathogen-commensal interactions is highly context-dependent. We refer to the phenomenon of detoxification followed by competition that suppresses PS as “SaxA-mediated biocontrol”. Our model predicts that this biocontrol increases for pathogens with higher ITC degradation rates, and when PS is outnumbered by commensals. Variable catalytic rates are likely in nature, since *in vitro* studies with diverse SaxA enzymes have shown different activities depending on their phylogenetic origin (18). In addition, SaxA-mediated biocontrol is predicted to decrease if the pathogen is more ITC-sensitive, for example due to fewer or less efficient ITC efflux pumps. It is known that SaxF-like ITC efflux pumps are diverse and differ in their effectiveness in protecting from different ITCs (40). Both ITC degradation and ITC sensitivity of pathogens also differ among the chemically diverse ITCs that occur in diverse Brassicaceae plants (18,40,41). In conclusion, this suggests that the outcome of the pathogen-commensal interaction depends on traits of all parties - plant host, opportunistic pathogen and commensals.

### ITC detoxification might be especially important for commensals when high ITC levels are released from a plant leaf

By degrading ITCs, PS causes a loss of function of these plant defense metabolites, which may promote disease because the plant’s ability to regulate opportunistic pathogens and to shape its microbiome is impaired (42). However, our results suggest that it could alternatively result in SaxA-mediated biocontrol. The suppression of pathogens by nutrient competition is a known principle of microbial biocontrol (43). This study shows that SaxA-mediated biocontrol is more likely when the ITC concentration is above a certain threshold (depending on the commensal’s ITC sensitivity) and the pathogen:commensal ratio is low. *Pseudomonas viridiflava* is a common colonizer of healthy *A thaliana* leaves, where it is present as a small but significant fraction of the microbiome (13). When plants are wounded, for example due to herbivory, ITC levels rapidly increase in leaf tissues (15). Opportunistic pathogens like *Pseudomonas viridiflava* might take advantage of wounded plant tissue and infect the leaf. However, under these conditions according to our model predictions ITC-sensitive commensals like *Janthinobacterium* sp. K would indirectly benefit from SaxA as well and might contribute to suppressing the pathogen. Up to now the ITC concentration to which individual commensal cells are exposed during such an attack has not been quantified. One study measured the bulk apoplastic concentration of 4MSOB-ITC of non-infected *A. thaliana* leaves (∼42 µM = 7.4 µg/mL) (34). These concentrations would be so low that according to our model ITC-sensitive commensals like K would not benefit from SaxA but could suppress PS anyway. However, without spatial information on ITC concentrations in leaves, it is difficult to estimate the exposure of individual bacterial cells to this plant toxin. Some bacterial cells might well be exposed to higher ITC concentrations locally where they might benefit from SaxA-mediated detoxification. In order to assess the relevance of our predictions *in planta*, further experiments and spatially resolved ITC data in leaves are necessary.

### Commensal rescue might indirectly benefit an opportunistic pathogen

A public good resistance mechanism like SaxA will non-selectively rescue any sensitive commensal that is close enough to the degrader strain to benefit from its ITC degradation. Enrichment of diverse bacteria could be beneficial for the host plant, since diverse commensals with a broad palette of resource niches can compete for nutrients with opportunistic pathogens, a phenomenon observed in both root and leaf microbiomes (8,9). Additionally, commensals could suppress pathogenicity by competing for virulence-inducing nutrients like fructose or certain amino acids (44,45). At any rate, our results suggest that SaxA-dependent biocontrol can occur under a variety of conditions, suggesting it could be rather common. Given that other, private resistance mechanisms like ITC efflux pumps are available which would not enrich competitors, an important question is whether PS itself might benefit from rescuing commensals. Leaves are thought to be very resource-poor environments (11,46). Thus, one possibility is that the rescued commensals could provide the opportunistic pathogen different nutrients or vitamins via cross-feeding (47,48) or have specialized adaptations that help the pathogen access host resources (20). This role of SaxA might thereby allow opportunistic pathogens to occupy new niches and become part of a healthy leaf microbiome. The surprising prevalence of ITC degradation probably mediated by SaxA-like enzymes among some of our commensal leaf colonizers additionally underlines that this trait is probably relevant beyond virulence. Therefore, instead of “virulence factor” we might rather call SaxA a “dual-use trait” (49). This concept initially described how virulence traits of human pathogens can emerge in environmental niches outside the human body because they confer advantages also in non-disease situations (50). Our results suggest that SaxA-mediated ITC degradation may shape not only the roles of plant, pathogen and commensals in virulence and in healthy microbiomes but also their evolutionary trajectories. If an opportunistic pathogen like *Pseudomonas viridiflava* is under pressure to cause disease, it will likely have to adapt differently than when it is part of a healthy microbiome. Thus, the context-dependent functions of factors like SaxA - conferring virulence to pathogens on the one hand, but benefiting commensal competitors on the other hand - might contribute to explain why different lineages of opportunistic *Pseudomonas viridiflava* co-exist in healthy *A. thaliana* leaves (13). While uncovering the roles of SaxA in the leaf microbiome will require more research, our results already suggest that it is important to carefully consider the role of “virulence traits” in plant-associated microbiomes.

### Model assumptions, limitations, and ecological relevance

Mathematical modelling played a central role in this study. We showed that a simple model, calibrated on monoculture growth data, can be used to quantitatively interpret interactions that occur in coculture. Furthermore, the model was used to predict scenarios beyond those that were experimentally tested, providing insights into the ecology of plant–pathogen–commensal systems. The natural environment is, of course, expected to be more complex than our simple model. For example, in our model we considered only one or two sequentially consumed nutrients, whereas in plant leaves co-utilization is possible (51,52), along with the secretion of secondary metabolites (53,54). ITC degradation and toxicity were modelled phenomenologically, without explicit representation of their pathways or molecular mechanisms. However, this simple approach adequately described the *in vitro* system. Additionally, our analysis focused mainly on single-species and pairwise cultures, whereas the study of complex microbial communities, especially in a biofilm context such as on leaves, may require consideration of higher-order interactions among multiple bacterial species (55,56). In addition, we did not take account of factors such as pH and temperature shifts (57,58), production of secondary metabolite released in the environment (59), species invasions or migrations (60), and evolutionary dynamics (61). While our model could be extended to account for those cases, our aim here was mostly to study pairwise interactions. Importantly, bacteria on the plant leaf often display non-uniform spatial distributions (62). Capturing such spatial organization would require a more complex, spatially explicit modelling framework, which lies beyond the scope of the present work (63,64). Despite these simplifications, we demonstrated that a minimal phenomenological model incorporating a small number of key factors can successfully explain pairwise interactions in liquid cocultures and generate testable predictions relevant to natural systems.

## Outlook

This study showed how plant pathogen traits that are associated with virulence but function as public good and benefit commensals are likely to also play roles in a balanced, healthy leaf microbiome. The traits of the plant host, opportunistic pathogen and commensal microbes together shape the interplay between the public goods role of the virulence factor SaxA, and nutrient competition, in a way that can potentially benefit plant health. However, ecological interactions in plant leaves are likely to be very different than those observed *in vitro,* due to a variety of factors. Thus, as a next step it will be important to adapt our model to study these interactions *in planta* to understand whether, for example, nutrient competition indeed contributes to a balanced, healthy leaf microbiome containing opportunistic pathogens as well as how virulence is further suppressed. Together with *in planta* studies our results may eventually help in designing more efficient biocontrol agents.

## Funding

KU received funding from the Carl Zeiss Foundation (via the Jena School for Microbial Communication). MTA, KU, MM and RJA were funded by the Deutsche Forschungsgemeinschaft (DFG, German Research Foundation) under Germanýs Excellence

Strategy - EXC 2051 - Project-ID 390713860. JG acknowledges funding by the Max Planck Society.

### Acknowledgements

We thank Dr. Michael Reichelt (Max-Planck-Institute for Chemical Ecology, Jena) for the chemical analysis of 4MSOB-ITC and its breakdown products. We thank René Maskos and Prof. Kirsten Küsel (Friedrich Schiller University Jena) for sequencing our amplicon sequencing libraries on their MiSeq instrument. We thank Rudolph Schlechter and Prof. Mitja Remus-Emsermann (Freie Universität Berlin) for providing the pMRE-Tn7-145 plasmid for fluorescent tagging, and Rebecca Ruiter for assistance with the tagging.

## Supporting information captions

**Supplementary Figure 1: Influence of fluorescent tagging with mScarlet-I on bacterial growth.** At different ITC concentrations the OD_600_ of non-tagged PS or PSKO were compared to mScarlet-I-tagged versions of PS or PSKO, respectively. The results are shown as means (dots) and standard deviations (shaded areas) over time. The mean for PS over time and ITC concentrations was 0.994 (sd=0.006) and for PSKO it was 1.04 (sd=0.02). Hence tagging did not influence PS or PSKO’s growth with different 4MSOB-ITC concentrations.

**Supplementary Figure 2: Controls and normalization of raw reads of the amplicon sequencing experiment. (A)** Raw reads of non-inoculated medium after 24 h incubation. Here only Flavobacteriaceae (*Imtechella halotolerans*, gram-negative standard taxon) and Bacilliaceae (*Allobacillus halotolerans*, gram-positive standard taxon) were amplified. **(B)** Raw read counts of negative controls (NFW was added to the PCRs instead of DNA template) were below 250 reads, and positive controls amplified a community standard (Zymo-mix) as expected. **(C)** Comparison of non-normalized (upper panel, raw read counts) and normalized (lower panel, artificial abundance) abundances of the five SynCom taxa and PS/PSKO in the whole library. The first 35 samples belong to SynCom5 (0-24h, n=5 each timepoint), the next SynCom5+SaxA and the last SynCom5-SaxA. The last three samples show the three inocula (“Inoc.”). Especially gram-positive taxa (*Plantibacter* G, *Rhodococcus* R) were increased by the normalization to the respective gram-positive internal standard taxon.

**Supplementary Figure 3: SynComs grew over time and only PS degraded 4MSOB-ITC.** All five commensals were mixed in equal amounts (SynCom5) and either PS (SynCom5+SaxA) or PSKO (SynCom5-SaxA) were added. **(A)** Measurements of red fluorescence (mScarlet-I) and OD_600_ in each community (n=5) at the sampling timepoints between 0 and 24 h. **(B)** Quantification of 4MSOB-ITC and 4MSOB-amine in all samples after 24 h (n=5).

**Supplementary Figure 4: Validation of the model. (A, B)** After obtaining the values of the parameters by fitting the monocultures, we validated the mathematical model by predicting the dynamics of the total OD_600_ of pairwise cultures of mScarlet-I tagged PS (PSfluo) and PSKO, and mScarlet-I tagged PSKO (PSKOfluo) and PS, with an increasing ITC concentration. The strains were mixed with an initial ratio of 1 (same amount) and the total OD_600_ was measured over 70 h (three replicates, averages and standard deviations as dotted lines and shaded regions, respectively). The model predictions are given as solid lines and are the same for panel A and B, since the two strains have the same set of parameter values. **(C)** Replot of the model prediction of the total OD_600_ for the cocultures in A and B. **(D)** Predicted individual OD_600_ of either only PS, PSfluo, PSKO, or PSKOfluo in the pairwise cultures. The OD_600_ is the same in all cases, because of the symmetry of the model between the two strains in each simulation.

**Supplementary Figure 5: Characteristics of commensal colonizers at 30°C. (A)** Quantification of 4MSOB-ITC and its degradation products 4MSOB-amine and 4MSOB-ITC-GSH conjugate in supernatants of potential SynCom members. A letter code (A to S) was assigned to each strain (see Suppl. Tab. 1). Strains which did not grow in the 5 mL pre-culture within 24 h were not tested. (n=3, no_bac = non-inoculated control). **(B)** Growth curves of potential SynCom members at 30°C in R2A broth. *Janthinobacterium* sp. K does not grow at this temperature, so we shifted to 28°C for the main experiments. Solid lines depict the means; shades illustrate the standard deviations.

**Supplementary Figure 6: Abundance of SynCom taxa at each sampling timepoint.** Normalized abundance of each SynCom taxon over time at 0, 3, 5, 7, 9, 11 and 24 h. Dots show individual samples (n=5 technical replicates), boxes depict the median +/- interquartile range, whiskers show +/- 1.5x interquartile range. Differences between treatments were assessed using t-tests, only significant comparisons (p<0.05) are shown (adjusted p-values, using Bonferroni method).

**Supplementary Figure 7: Fit of PS and PSKO with a single nutrient model to the data. (A)** and **(B)** show the fit of the model (solid lines) with a single nutrient assimilation (Equations 1-3 in Fig. 1B in the main text) to the data (averages and standard deviations over three replicates as dots and shaded regions, respectively) for several ITC concentrations.

**Supplementary Figure 8: Alternative models for commensal E and R.** (**A)** Fit of an alternative model with diauxic shift between two nutrients for the growth of the commensal E. The improvement compared to a single nutrient of Fig. 5B in the main text is not enough to justify a more complex mathematical model with four more parameters. **(B)** Fit of an alternative model with a single nutrient for the commensal R. Here, the single nutrient even fails to catch the trend of the data, therefore a two-nutrient model was used.

**Supplementary Figure 9: R and G form biofilms and/or aggregates which falsify the OD readings.** To check the formation of biofilm and aggregates for the commensal G and R beyond a visual inspection (panel A and B, respectively), we growth them for a longer time (120 h). We verified that their OD dynamics present several complex phases, large oscillations, and high standard deviations, possible indication of inhomogeneous densities in the media induced by bacterial clumps, especially for higher ITC concentrations.

**Supplementary Figure 10: Pairwise predictions of PS(KO) and commensals G, M and R. (A, C, and E)** The model prediction for the total biomass dynamics (OD_600_) of the pairwise culture starting with a same amount of either PS or PSKO and commensal G, M, and R for different ITC concentrations are shown by the solid lines and compared to the experimental data (dots, averages of three technical replicates, standard deviations as shaded region), in panel A, C, and E, respectively. **(B, D, and F)** Separate dynamics of the biomass of either PS or PSKO and the commensal G, M, and R. Solid lines show the OD_600_ of PS or PSKO, and dashed lines the OD_600_ of the commensal. The total biomass is mostly determined by the pathogen, and the commensal biomass reaches a maximum of one fourth of the pathogen biomass.

**Supplementary Figure 11: OD_600_ dynamics of pathogens PS and PSKO growing in the absence or presence of commensal K for several initial PS(KO):K ratios. (A)** OD_600_ dynamics of either PS or PSKO (solid lines, top and bottom row, respectively) and K (dashed lines) for several ITC concentrations and three initial PS(KO):K ratio (1:100, 1:1, 100:1, first, second and third column, respectively). Panel **(B)** shows the same conditions but for monocultures of either PS or PSKO, in the absence of commensal K. Predictions in A and B illustrate how commensal K influences growth of PS or PSKO, which is formalized in the suppression index.

**Supplementary Figure 12: Heatmaps of maximal OD_600_ used to calculate the indices.** Summary of the maximal OD_600_ extracted from the dynamics of the pathogen and commensal biomass as explained in Suppl. Text 3. Panel A shows the maximal OD_600_ attained by the monocultures of PS. PSref indicate the reference initial value of PS (OD_600_ = 0.0025). Panel B shows the maximal OD_600_ of PSKO attained by the pairwise culture of PSKO and K. Panels C and D show the maximal OD_600_ of PS and K attained by the pairwise culture of PS and K. The dashed black line in C and the dot-dashed blue line in D indicate the conditions where the maximal OD_600_ reaches 0.06 (isoline), respectively.

**Supplementary Figure 13: Growth of different initial ratios of PS/PSKO paired with a commensal.** Growth curves of OD_600_ **(A)** and red fluorescence **(B)** of different initial mixtures of PS or PSKO tagged with mScarlet-I mixed with either the commensals E or K, or PSKO as control. (PS(KO):C ratios ranged from 100:1 to 1:100. The total initial OD_600_ was kept at 0.4. The average of three technical replicates is shown (solid line) with standard deviation (shaded area).

**Supplementary Figure 14: Indices for the most ITC-resistant commensal strain E.** Panels A, B, and C: Heat maps of the rescue, suppression, and SaxA-mediated biocontrol index, varying the initial PS(KO):E ratio and for several ITC concentrations. The black dashed and blue dot-dashed lines in C represent where the maximal OD_600_ of PS and E are 0.06, respectively. Areas under the curves in C define the SaxA-mediated biocontrol region where commensal growth benefits more from SaxA than PS, the color illustrates the strength of the potential effect. It is worth noticing that the rescue index does not distinguish between a case when the rescue fails (e.g. ineffective SaxA-dependent degradation of ITC) and a case when there is no need to rescue because the commensal strain presents a low ITC sensitivity (e.g. as for E). To distinguish between these conditions, we introduce in Suppl. Fig. 15 a new index that quantifies the “need-to-be rescued” as ITC-sensitivity index.

**Supplementary Figure 15: ITC-sensitivity index for commensal strain K and E.** Panels A and B show the rescue indices of strains K and E, which were already presented in Fig. 7B (main text) and Suppl. Fig. 14A. Importantly, the rescue index does not provide information on the underlying reason for the rescue outcome. For example, the red region in the rescue index of strain E indicates that E is not rescued in most conditions. However, this is not due to an inefficient SaxA-dependent ITC degradation, but rather to the fact that strain E intrinsically does not require rescue, owing to its low ITC sensitivity. To disentangle these possibilities, we introduce in Panel C the ITC-sensitivity index, which quantifies the sensitivity of each strain to 4MSOB-ITC on a scale from 0 to 1. The index compares the max OD attained by a commensal strain C grown in the presence of PSKO to the same condition but in the absence of ITC (C+PSKO and [ITC]=0). In this representation, green regions denote high ITC sensitivity and thus a potential requirement for rescue, whereas red regions correspond to ITC insensitivity and therefore no need for rescue of the commensal strain. Panel D shows that strain K is highly ITC-sensitive under most conditions, except at low [ITC] and high cell numbers, consistent with the pattern observed in the rescue index. By contrast, Panel E clarifies why the rescue index of strain E shows an apparent lack of rescue: strain E is largely ITC-insensitive across the explored parameter space and therefore does not require rescue. Indeed, E grows robustly even at high [ITC] (see Suppl. Fig. 16).

**Supplementary Figure 16: Pairwise prediction for PS(KO):E at different initial ratios.** Model predictions (solid lines) and experimental measurements (average as dots and shaded regions as standard deviations of three independent replicates) for the OD_600_ of the pairwise cultures PS:E and PSKO:E, in the first and second row, respectively, and for different PS(KO):E ratios (1:100, 1:10, 1:1, 10:1, 100:1).

**Supplementary Figure 17: Indices for a hypothetical pathogen PS* either with increased ITC sensitivity or with increased ITC degradation rate.** Heat maps of the rescue, suppression, and SaxA-mediated biocontrol index for the pathogen PS **(A)**, for hypothetical pathogen PS* with either tenfold higher ITC sensitivity (**B**, corresponding to a tenfold smaller value of the parameter q in the mathematical model, Equation 8 in Suppl. Text 2) or with a tenfold increased ITC degradation rate (**C**, corresponding to tenfold larger value of λ in the mathematical model, Equation 12 of Suppl. Text 2), varying the initial PS(KO)*:K ratio and for several ITC concentrations. The black dashed and blue dot-dashed lines in C represent where the maximal OD_600_ of PS* and K are 0.06, respectively. The white hatched region in the biocontrol index defines the region where the SaxA-mediated biocontrol is maximal and both PS cells are not too abundant, and K cells are present with a sensitive number.

**Supplementary Figure 18: Indices for PS with lower ITC sensitivity.** Panels A, B, and C: Heat maps of the rescue, suppression, and SaxA-mediated biocontrol index for a pathogen PS* obtained by lowering the ITC sensitivity of PS (corresponding to tenfold larger K in the mathematical model, Equation 8 of Suppl. Text 2) and varying the initial PS(KO)*:K ratio and for several ITC concentrations. The black dashed and blue dot-dashed lines in C represent where the maximal OD_600_ of PS* and K are 0.06, respectively. Areas under the curves in C define the SaxA-mediated biocontrol region where commensal growth benefits more from SaxA than PS, the color illustrates the strength of the potential effect.

**Supplementary Figure 19: Indices for PS with lower ITC degradation rate.** Heat maps for the indices, as described in Suppl. Fig. 18, for a pathogen PS* obtained by lowering the ITC degradation rate of PS (corresponding to tenfold smaller λ in the mathematical model, Equation 12 of Suppl. Text 2).

**Supplementary Figure 20: Indices for PS with higher growth rate.** Heat maps for the indices, as described in Suppl. Fig. 18, for a pathogen PS* obtained by increasing the growth rate of PS (corresponding to twofold larger µ_1_ in the mathematical model, Equation 8 of Suppl. Text 2).

**Supplementary Figure 21: Indices for PS with lower growth rate.** Heat maps for the indices, as described in Suppl. Fig. 18, for a pathogen PS* obtained by lowering the growth rate of PS (corresponding to twofold smaller µ_1_ in the mathematical model, Equation 8 of Suppl. Text 2). Potential plant beneficial regions are in the lower left corner below the blue and black lines.

**Supplementary Figure 22: Indices for K with higher growth rate.** Heat maps for the indices, as described in Suppl. Fig. 18, for a pathogen K* obtained by increasing the growth rate of K (corresponding to twofold larger µ_C_ in the mathematical model, Equation 9 of Suppl. Text 2). Potential plant beneficial regions are in the lower left corner below the blue and black lines.

**Supplementary Figure 23: Indices for K with lower growth rate.** Heat maps for the indices, as described in Suppl. Fig. 18, for a pathogen K* obtained by lowering the growth rate of K (corresponding to twofold smaller µ_C_ in the mathematical model, Equation 9 of Suppl. Text 2).

**Supplementary Figure 24: Indices for K when more susceptible to ITC.** Heat maps for the indices, as described in Suppl. Fig. 18, for a pathogen K* obtained by increasing the susceptibility to ITC (corresponding to tenfold smaller K_C_ in the mathematical model, Equation 9 of Suppl. Text 2).

**Supplementary Figure 25: Indices for K when less susceptible to ITC.** Heat maps for the indices, as described in Suppl. Fig. 18, for a pathogen K* obtained by lowering the susceptibility to ITC (corresponding to tenfold larger K_C_ in the mathematical model, Equation 9 of Suppl. Text 2). Since K* is almost insensitive to ITC, it does not need to be rescued by the ITC-degrader PS. For some low ranges of ITC concentration, K* grows better in the presence of PSKO than PS, thus leading to negative values of the rescue index (indicated by the gray regions). Therefore, unlike all the cases here analyzed, the rescue index and therefore the biocontrol index can be negative (gray regions).

## References

1. Finlay BB, Falkow S. Common themes in microbial pathogenicity revisited. Microbiol Mol Biol Rev. 1997 Jun;61(2):136–69.

2. Leveau JHJ. Re-envisioning the plant disease triangle: full integration of the host microbiota and a focal pivot to health outcomes. Annual Review of Phytopathology. 2024 Sep 9;62(1):31–47.

3. Bass D, Stentiford GD, Wang HC, Koskella B, Tyler CR. The pathobiome in animal and plant diseases. Trends in Ecology & Evolution. 2019 Nov;34(11):996–1008.

4. Kettle AJ, Batley J, Benfield AH, Manners JM, Kazan K, Gardiner DM. Degradation of the benzoxazolinone class of phytoalexins is important for virulence of *Fusarium pseudograminearum* towards wheat. Mol Plant Pathol. 2015 Apr 15;16(9):946–62.

5. van den Bosch TJM, Niemi O, Welte CU. Single gene enables plant pathogenic *Pectobacterium* to overcome host-specific chemical defence. Mol Plant Pathol. 2020 Mar;21(3):349–59.

6. Liu X, Keyhani NO, Liu H, Zhang Y, Xia Y, Cao Y. Glyoxal oxidase-mediated detoxification of reactive carbonyl species contributes to virulence, stress tolerance, and development in a pathogenic fungus. PLOS Pathogens. 2024 Jul 30;20(7):e1012431.

7. Ökmen B, Etalo DW, Joosten MHAJ, Bouwmeester HJ, de Vos RCH, Collemare J, et al. Detoxification of α-tomatine by *Cladosporium fulvum* is required for full virulence on tomato. New Phytologist. 2013;198(4):1203–14.

8. Berg M, Koskella B. Nutrient- and Dose-Dependent Microbiome-Mediated Protection against a Plant Pathogen. Curr Biol. 2018 Aug 6;28(15):2487–2492.e3.

9. Yang T, Wei Z, Friman VP, Xu Y, Shen Q, Kowalchuk GA, et al. Resource availability modulates biodiversity-invasion relationships by altering competitive interactions. Environ Microbiol. 2017 Aug;19(8):2984–91.

10. Garcia-Santamarina S, Kuhn M, Devendran S, Maier L, Driessen M, Mateus A, et al. Emergence of community behaviors in the gut microbiota upon drug treatment. Cell. 2024 Oct 31;187(22):6346–6357.e20.

11. Vorholt JA. Microbial life in the phyllosphere. Nat Rev Microbiol. 2012 Dec;10(12):828–40.

12. Bai Y, Müller DB, Srinivas G, Garrido-Oter R, Potthoff E, Rott M, et al. Functional overlap of the *Arabidopsis* leaf and root microbiota. Nature. 2015 Dec;528(7582):364–9.

13. Karasov TL, Almario J, Friedemann C, Ding W, Giolai M, Heavens D, et al. *Arabidopsis thaliana* and *Pseudomonas* pathogens exhibit stable associations over evolutionary timescales. Cell Host Microbe. 2018 Jul 11;24(1):168–179.e4.

14. Halkier BA, Gershenzon J. Biology and biochemistry of glucosinolates. Annu Rev Plant Biol. 2006;57:303–33.

15. Textor S, Gershenzon J. Herbivore induction of the glucosinolate–myrosinase defense system: major trends, biochemical bases and ecological significance. Phytochem Rev. 2009 Jan 1;8(1):149–70.

16. Andersson MX, Nilsson AK, Johansson ON, Boztaş G, Adolfsson LE, Pinosa F, et al. Involvement of the electrophilic isothiocyanate sulforaphane in *Arabidopsis* local defense responses. Plant Physiology. 2015 Jan 1;167(1):251–61.

17. Fan J, Crooks C, Creissen G, Hill L, Fairhurst S, Doerner P, et al. *Pseudomonas sax* genes overcome aliphatic isothiocyanate-mediated non-host resistance in *Arabidopsis*. Science. 2011 Mar 4;331(6021):1185–8.

18. van den Bosch TJM, Tan K, Joachimiak A, Welte CU. Functional profiling and crystal structures of isothiocyanate hydrolases found in gut-associated and plant-pathogenic bacteria. Appl Environ Microbiol. 2018 Jul 15;84(14):e00478–18.

19. Chen J, Ullah C, Reichelt M, Beran F, Yang ZL, Gershenzon J, et al. The phytopathogenic fungus *Sclerotinia sclerotiorum* detoxifies plant glucosinolate hydrolysis products via an isothiocyanate hydrolase. Nat Commun. 2020 Jun 18;11(1):3090.

20. Unger K, Raza SAK, Mayer T, Reichelt M, Stuttmann J, Hielscher A, et al. Glucosinolate structural diversity shapes recruitment of a metabolic network of leaf-associated bacteria. Nat Commun. 2024 Oct 1;15(1):8496.

21. Madsen SR, Olsen CE, Nour-Eldin HH, Halkier BA. Elucidating the role of transport processes in leaf glucosinolate distribution. Plant Physiol. 2014 Nov;166(3):1450–62.

22. Sugiyama R, Li R, Kuwahara A, Nakabayashi R, Sotta N, Mori T, et al. Retrograde sulfur flow from glucosinolates to cysteine in *Arabidopsis thaliana*. Proc Natl Acad Sci U S A. 2021 Jun 1;118(22):e2017890118.

23. Unger K, Ruiter R, Reichelt M, Gershenzon J, Agler MT. *Pseudomonas* virulence factor SaxA detoxifies plant glucosinolate hydrolysis products, rescuing a commensal that suppresses virulence gene expression. bioRxiv. 2025 Apr 25;2025.04.25.650564.

24. Reasoner DJ, Geldreich EE. A new medium for the enumeration and subculture of bacteria from potable water. Appl Environ Microbiol. 1985 Jan;49(1):1–7.

25. Schlechter RO, Jun H, Bernach M, Oso S, Boyd E, Muñoz-Lintz DA, et al. Chromatic bacteria – A broad host-range plasmid and chromosomal insertion toolbox for fluorescent protein expression in bacteria. Front Microbiol. 2018 Dec 12;9:423623.

26. Lundberg DS, Pramoj Na Ayutthaya P, Strauß A, Shirsekar G, Lo WS, Lahaye T, et al. Host-associated microbe PCR (hamPCR) enables convenient measurement of both microbial load and community composition. Elife. 2021 Jul 22;10:e66186.

27. Callahan BJ, McMurdie PJ, Rosen MJ, Han AW, Johnson AJA, Holmes SP. DADA2: High-resolution sample inference from Illumina amplicon data. Nat Methods. 2016 Jul;13(7):581–3.

28. Quast C, Pruesse E, Yilmaz P, Gerken J, Schweer T, Yarza P, et al. The SILVA ribosomal RNA gene database project: improved data processing and web-based tools. Nucleic Acids Res. 2013 Jan;41(Database issue):D590–596.

29. McMurdie PJ, Holmes S. phyloseq: an R package for reproducible interactive analysis and graphics of microbiome census data. PLoS One. 2013;8(4):e61217.

30. Wickham H. ggplot2: Elegant Graphics for Data Analysis [Internet]. Springer-Verlag New York; 2016. Available from: https://ggplot2.tidyverse.org

31. Allen RJ, Waclaw B. Bacterial growth: a statistical physicist’s guide. Rep Prog Phys. 2019 Jan 1;82(1):016601.

32. Regoes RR, Wiuff C, Zappala RM, Garner KN, Baquero F, Levin BR. Pharmacodynamic functions: a multiparameter approach to the design of antibiotic treatment regimens. Antimicrob Agents Chemother. 2004 Oct;48(10):3670–6.

33. Young P. Everything You Wanted to Know About Data Analysis and Fitting but Were Afraid to Ask [Internet]. Cham: Springer International Publishing; 2015 [cited 2025 May 21]. (SpringerBriefs in Physics). Available from: https://link.springer.com/10.1007/978-3-319-19051-8

34. Wang W, Yang J, Zhang J, Liu YX, Tian C, Qu B, et al. An *Arabidopsis* secondary metabolite directly targets expression of the bacterial type III secretion system to inhibit bacterial virulence. Cell Host Microbe. 2020 Apr 8;27(4):601–613.e7.

35. Aires A, Mota VR, Saavedra MJ, Monteiro AA, Simões M, Rosa E a. S, et al. Initial *in vitro* evaluations of the antibacterial activities of glucosinolate enzymatic hydrolysis products against plant pathogenic bacteria. J Appl Microbiol. 2009 Jun;106(6):2096–105.

36. Sorg RA, Lin L, Doorn GS van, Sorg M, Olson J, Nizet V, et al. Collective resistance in microbial communities by intracellular antibiotic deactivation. PLOS Biology. 2016 Dec 27;14(12):e2000631.

37. Domingues IL, Gama JA, Carvalho LM, Dionisio F. Social behaviour involving drug resistance: the role of initial density, initial frequency and population structure in shaping the effect of antibiotic resistance as a public good. Heredity. 2017 Nov;119(5):295–301.

38. Verdon N, Popescu O, Titmuss S, Allen RJ. Habitat fragmentation enhances microbial collective defence. J R Soc Interface. 2025 Feb;22(223):20240611.

39. Piccardi P, Vessman B, Mitri S. Toxicity drives facilitation between 4 bacterial species. Proc Natl Acad Sci U S A. 2019 Aug 6;116(32):15979–84.

40. Russ D, Fitzpatrick CR, Saha C, Law TF, Jones CD, Kliebenstein DJ, et al. An effluent pump family distributed across plant commensal bacteria conditions host- and organ-specific glucosinolate detoxification. Nat Commun. 2025 Jul 1;16(1):5699.

41. Witzel K, Hanschen FS, Schreiner M, Krumbein A, Ruppel S, Grosch R. *Verticillium* suppression is associated with the glucosinolate composition of *Arabidopsis thaliana* leaves. PLOS ONE. 2013 Sep 5;8(9):e71877.

42. Arnault G, Mony C, Vandenkoornhuyse P. Plant microbiota dysbiosis and the Anna Karenina Principle. Trends in Plant Science. 2023 Jan 1;28(1):18–30.

43. Köhl J, Kolnaar R, Ravensberg WJ. Mode of Action of Microbial Biological Control Agents Against Plant Diseases: Relevance Beyond Efficacy. Front Plant Sci. 2019;10:845.

44. Turner SE, Pang YY, O’Malley MR, Weisberg AJ, Fraser VN, Yan Q, et al. A DeoR-type transcription regulator is required for sugar-induced expression of type III secretion-encoding genes in *Pseudomonas syringae* pv. *tomato* DC3000. Mol Plant Microbe Interact. 2020 Mar;33(3):509–18.

45. Yan Q, Rogan CJ, Pang YY, Davis EW, Anderson JC. Ancient co-option of an amino acid ABC transporter locus in *Pseudomonas syringae* for host signal-dependent virulence gene regulation. PLoS Pathog. 2020 Jul 16;16(7):e1008680.

46. Leveau JHJ, Lindow SE. Appetite of an epiphyte: Quantitative monitoring of bacterial sugar consumption in the phyllosphere. Proceedings of the National Academy of Sciences. 2001 Mar 13;98(6):3446–53.

47. Murillo-Roos M, Abdullah HSM, Debbar M, Ueberschaar N, Agler MT. Cross-feeding niches among commensal leaf bacteria are shaped by the interaction of strain-level diversity and resource availability. ISME J. 2022 Sep;16(9):2280–9.

48. Ryback B, Bortfeld-Miller M, Vorholt JA. Metabolic adaptation to vitamin auxotrophy by leaf-associated bacteria. The ISME Journal. 2022 Dec 1;16(12):2712–24.

49. Morris CE, Bardin M, Kinkel LL, Moury B, Nicot PC, Sands DC. Expanding the paradigms of plant pathogen life history and evolution of parasitic fitness beyond agricultural boundaries. Rall GF, editor. PLoS Pathog. 2009 Dec 24;5(12):e1000693.

50. Casadevall A, Steenbergen JN, Nosanchuk JD. “Ready made” virulence and “dual use” virulence factors in pathogenic environmental fungi--the *Cryptococcus neoformans* paradigm. Curr Opin Microbiol. 2003 Aug;6(4):332–7.

51. Schlechter RO, Kear EJ, Bernach M, Remus DM, Remus-Emsermann MNP. Metabolic resource overlap impacts competition among phyllosphere bacteria. ISME J. 2023 Sep;17(9):1445–54.

52. Schäfer M, Pacheco AR, Künzler R, Bortfeld-Miller M, Field CM, Vayena E, et al. Metabolic interaction models recapitulate leaf microbiota ecology. Science. 2023 Jul 7;381(6653):eadf5121.

53. Perrin E, Ghini V, Giovannini M, Di Patti F, Cardazzo B, Carraro L, et al. Diauxie and co-utilization of carbon sources can coexist during bacterial growth in nutritionally complex environments. Nat Commun. 2020 Jun 19;11(1):3135.

54. Gross H, Loper JE. Genomics of secondary metabolite production by Pseudomonas spp. Nat Prod Rep. 2009 Oct 21;26(11):1408–46.

55. Guo X, Boedicker JQ. The Contribution of High-Order Metabolic Interactions to the Global Activity of a Four-Species Microbial Community. PLOS Computational Biology. 2016 Sep 13;12(9):e1005079.

56. Liu W, Røder HL, Madsen JS, Bjarnsholt T, Sørensen SJ, Burmølle M. Interspecific Bacterial Interactions are Reflected in Multispecies Biofilm Spatial Organization. Front Microbiol [Internet]. 2016 Aug 31 [cited 2025 Sep 11];7. Available from: https://www.frontiersin.org/journals/microbiology/articles/10.3389/fmicb.2016.01366/full

57. Ratkowsky DA, Olley J, McMeekin TA, Ball A. Relationship between temperature and growth rate of bacterial cultures. J Bacteriol. 1982 Jan;149(1):1–5.

58. Mougi A. pH Adaptation stabilizes bacterial communities. npj biodivers. 2024 Oct 17;3(1):32.

59. Pacheco AR, Moel M, Segrè D. Costless metabolic secretions as drivers of interspecies interactions in microbial ecosystems. Nat Commun. 2019 Jan 9;10(1):103.

60. Amor DR, Ratzke C, Gore J. Transient invaders can induce shifts between alternative stable states of microbial communities. Science Advances. 2020 Feb 19;6(8):eaay8676.

61. Huang Y, Mukherjee A, Schink S, Benites NC, Basan M. Evolution and stability of complex microbial communities driven by trade-offs. Molecular Systems Biology. 2024 Sep 2;20(9):997–1005.

62. Schlechter RO, Remus-Emsermann MNP. Differential responses of *Methylobacterium* and *Sphingomonas* species to multispecies interactions in the phyllosphere. Environmental Microbiology. 2025;27(1):e70025.

63. Song HS, Cannon WR, Beliaev AS, Konopka A. Mathematical Modeling of Microbial Community Dynamics: A Methodological Review. Processes. 2014 Dec;2(4):711–52.

64. Nagarajan K, Ni C, Lu T. Agent-Based Modeling of Microbial Communities. ACS Synth Biol. 2022 Nov 18;11(11):3564–74.

