## Supplementary Material for "Bacterial degradation of a plant toxin and nutrient competition with commensals trade off to constrain pathogen growth"

[+] contributed equally

Contents:

Supplementary Figures: 1 to 25

Supplementary Tables: 1 to 4

Supplementary References

Supplementary Texts: 1 to 3, attached as separate files

#### Supplementary Figures

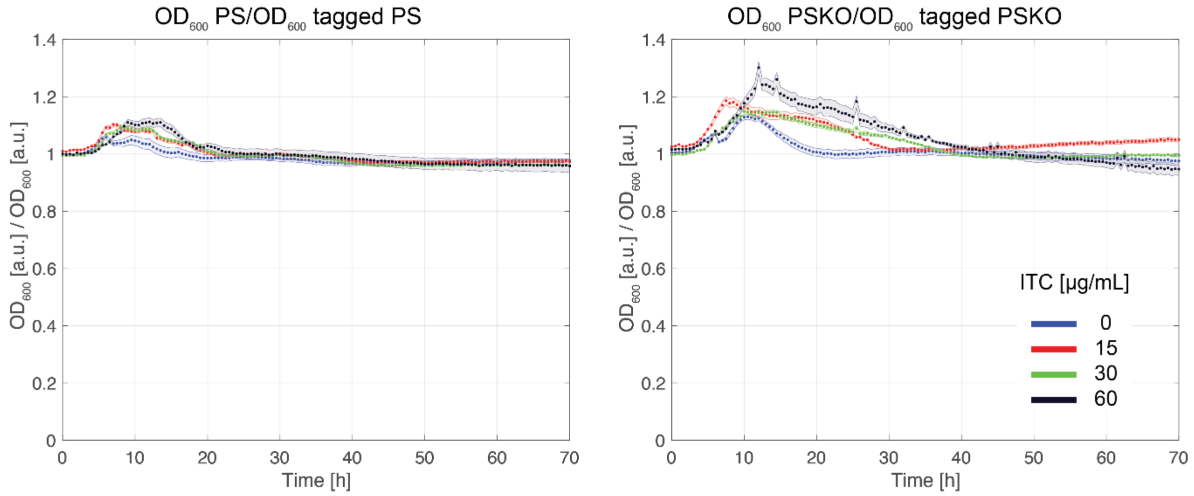


**Supplementary Figure 1: Influence of fluorescent tagging with mScarlet-I on bacterial growth.** At different ITC concentrations the $OD_{600}$ of non-tagged PS or PSKO were compared to mScarlet-I-tagged versions of PS or PSKO, respectively. The results are shown as means (dots) and standard deviations (shaded areas) over time. The mean for PS over time and ITC concentrations was 0.994 (sd=0.006) and for PSKO it was 1.04 (sd=0.02). Hence tagging did not influence PS or PSKO's growth with different 4MSOB-ITC concentrations.

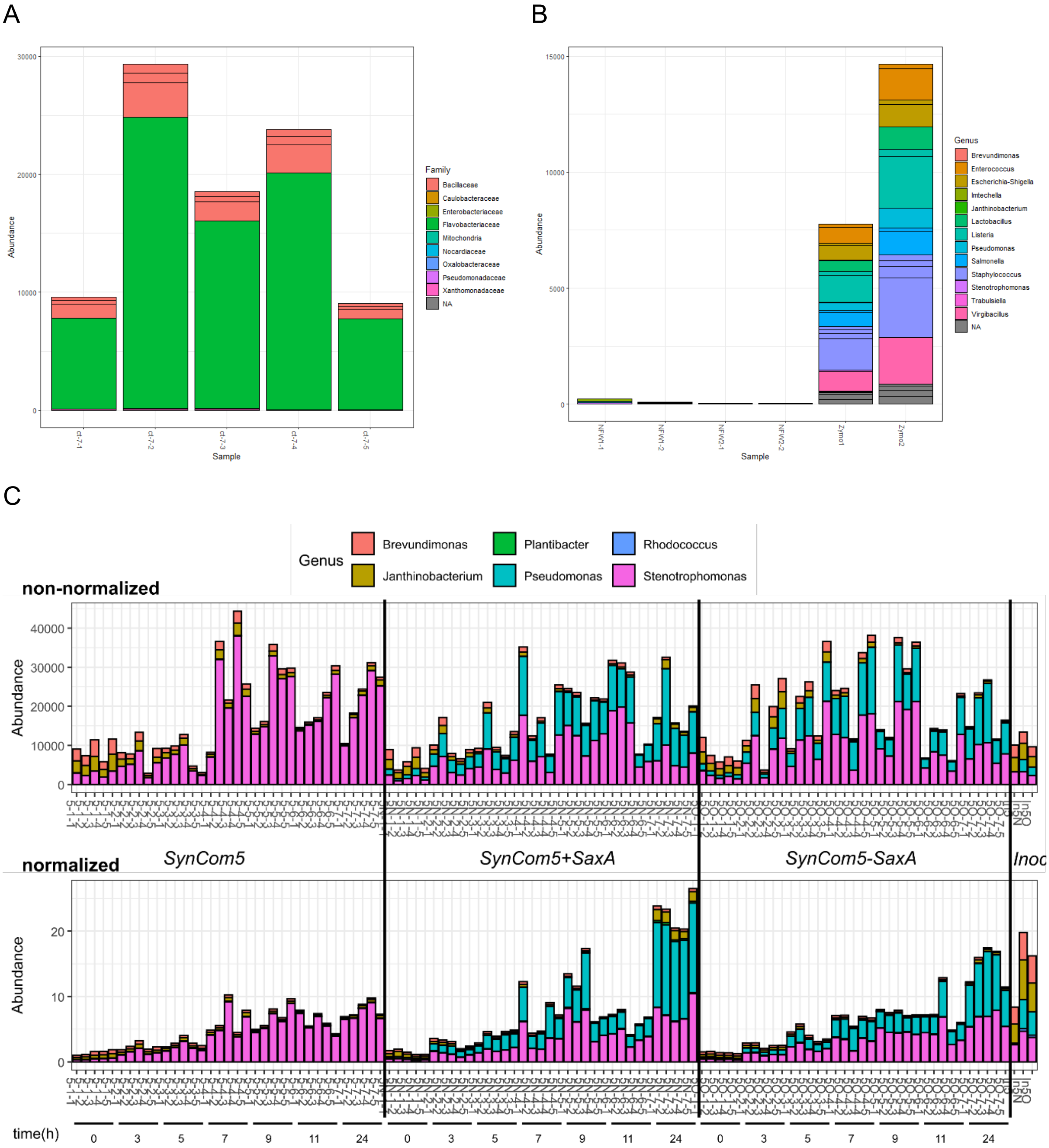


**Supplementary Figure 2: Controls and normalization of raw reads of the amplicon sequencing experiment. (A)** Raw reads of non-inoculated medium after 24 h incubation. Here only Flavobacteriaceae (*Imtechella halotolerans*, gram-negative standard taxon) and Bacilliaceae (*Allobacillus halotolerans*, gram-positive standard taxon) were amplified. **(B)** Raw read counts of negative controls (NFW was added to the PCRs instead of DNA template) were below 250 reads, and positive controls amplified a community standard (Zymo-mix) as expected. **(C)** Comparison of non-normalized (upper panel, raw read counts) and normalized (lower panel, artificial abundance) abundances of the five SynCom taxa and PS/PSKO in the whole library. The first 35 samples belong to SynCom5 (0-24h, n=5 each timepoint), the next SynCom5+SaxA and the last SynCom5-SaxA. The last three samples show the three inocula ("Inoc."). Especially gram-positive taxa (*Plantibacter* G, *Rhodococcus* R) were increased by the normalization to the respective gram-positive internal standard taxon.

A

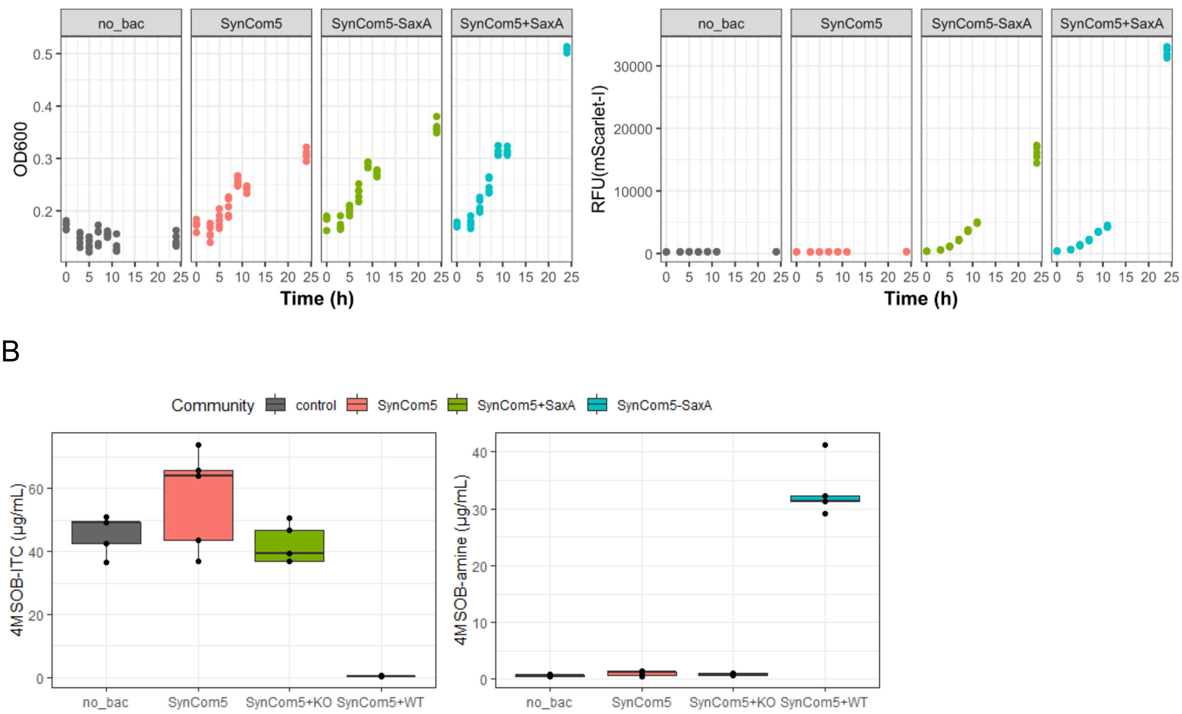


**Supplementary Figure 3: SynComs grew over time and only PS degraded 4MSOB-ITC.** All five commensals were mixed in equal amounts (SynCom5) and either PS (SynCom5+SaxA) or PSKO (SynCom5-SaxA) were added. **(A)** Measurements of red fluorescence (mScarlet-I) and $OD_{600}$ in each community (n=5) at the sampling timepoints between 0 and 24 h. **(B)** Quantification of 4MSOB-ITC and 4MSOB-amine in all samples after 24 h (n=5).

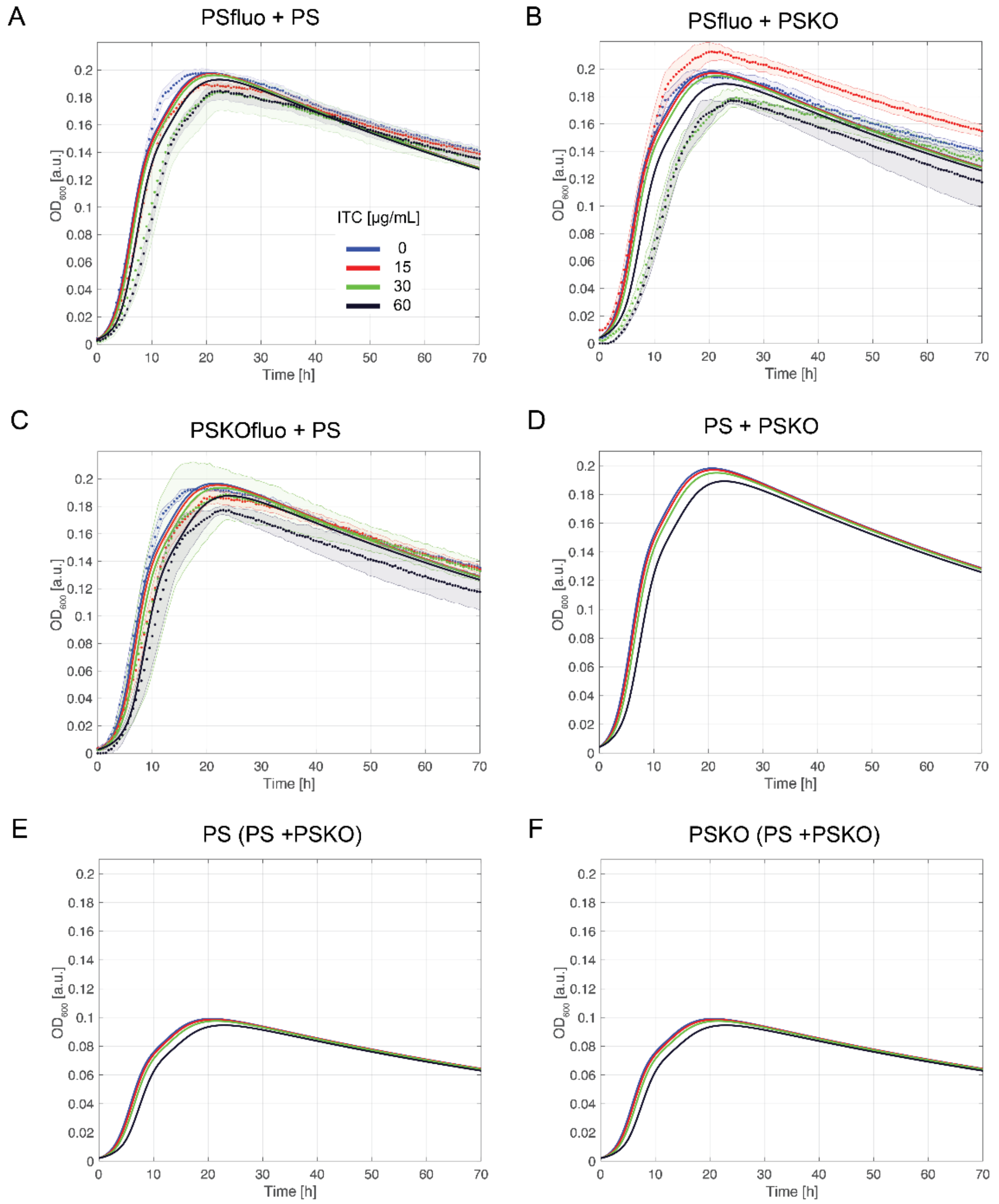


**Supplementary Figure 4: Validation of the model. (A, B)** After obtaining the values of the parameters by fitting the monocultures, we validated the mathematical model by predicting the dynamics of the total $OD_{600}$ of pairwise cultures of mScarlet-I tagged PS (PSfluo) and PSKO, and mScarlet-I tagged PSKO (PSKOfluo) and PS, with an increasing ITC concentration. The strains were mixed with an initial ratio of 1 (same amount) and the total $OD_{600}$ was measured over 70 h (three replicates, averages and standard deviations as dotted lines and shaded regions, respectively). The model predictions are given as solid lines and are the same for panel A and B, since the two strains have the same set of parameter values. **(C)** Replot of the model prediction of the total $OD_{600}$ for the cocultures in A and B. **(D)** Predicted individual $OD_{600}$ of either only PS, PSfluo, PSKO, or PSKOfluo in the pairwise cultures. The $OD_{600}$ is the same in all cases, because of the symmetry of the model between the two strains in each simulation.

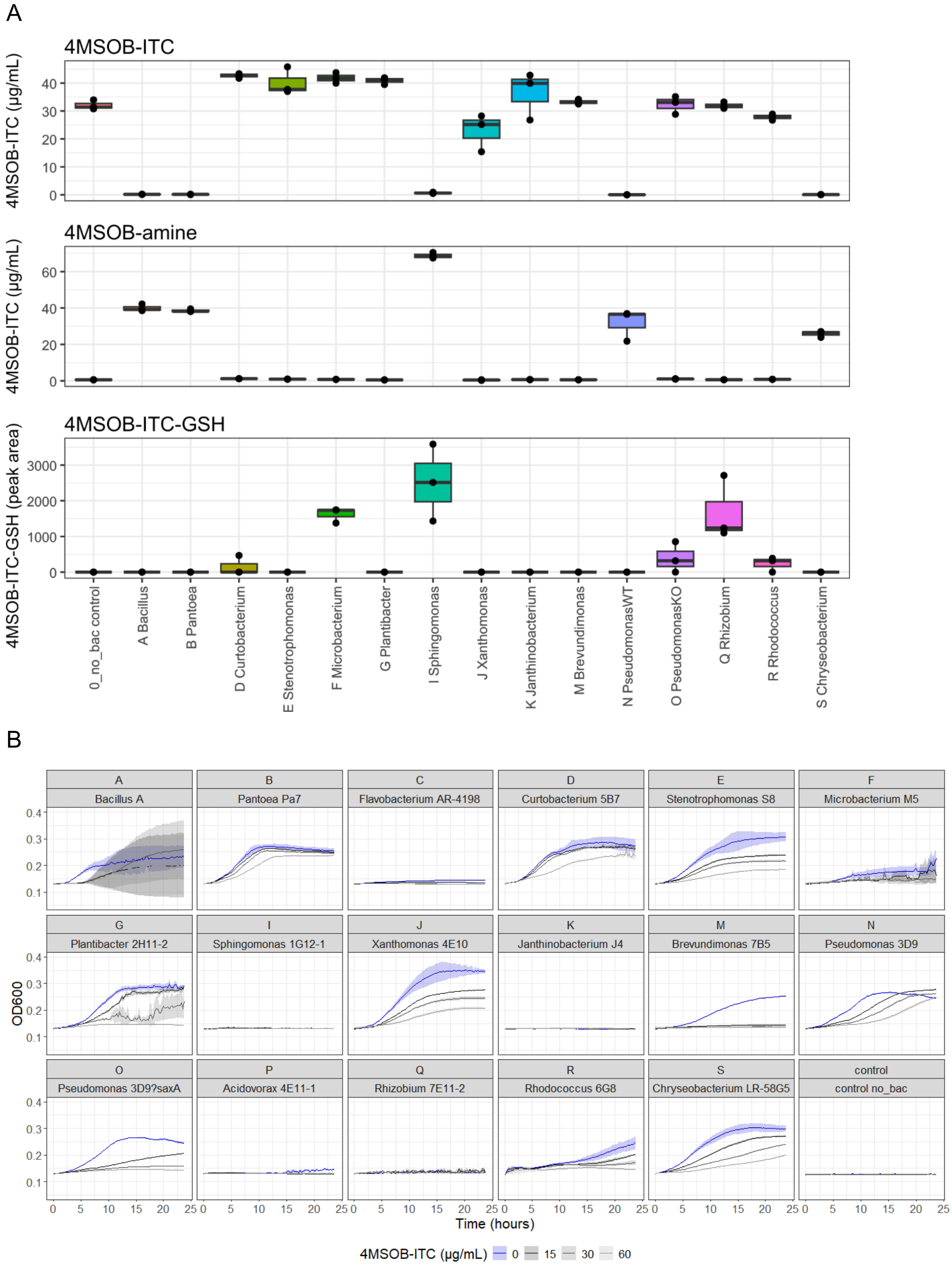


**Supplementary Figure 5: Characteristics of commensal colonizers at 30°C. (A)** Quantification of 4MSOB-ITC and its degradation products 4MSOB-amine and 4MSOB-ITC-GSH conjugate in supernatants of potential SynCom members. A letter code (A to S) was assigned to each strain (see Suppl. Tab. 1). Strains which did not grow in the 5 mL pre-culture within 24 h were not tested. (n=3, no_bac = non-inoculated control). **(B)** Growth curves of potential SynCom members at 30°C in R2A broth. *Janthinobacterium* sp. K does not grow at this temperature, so we shifted to 28°C for the main experiments. Solid lines depict the means; shades illustrate the standard deviations.

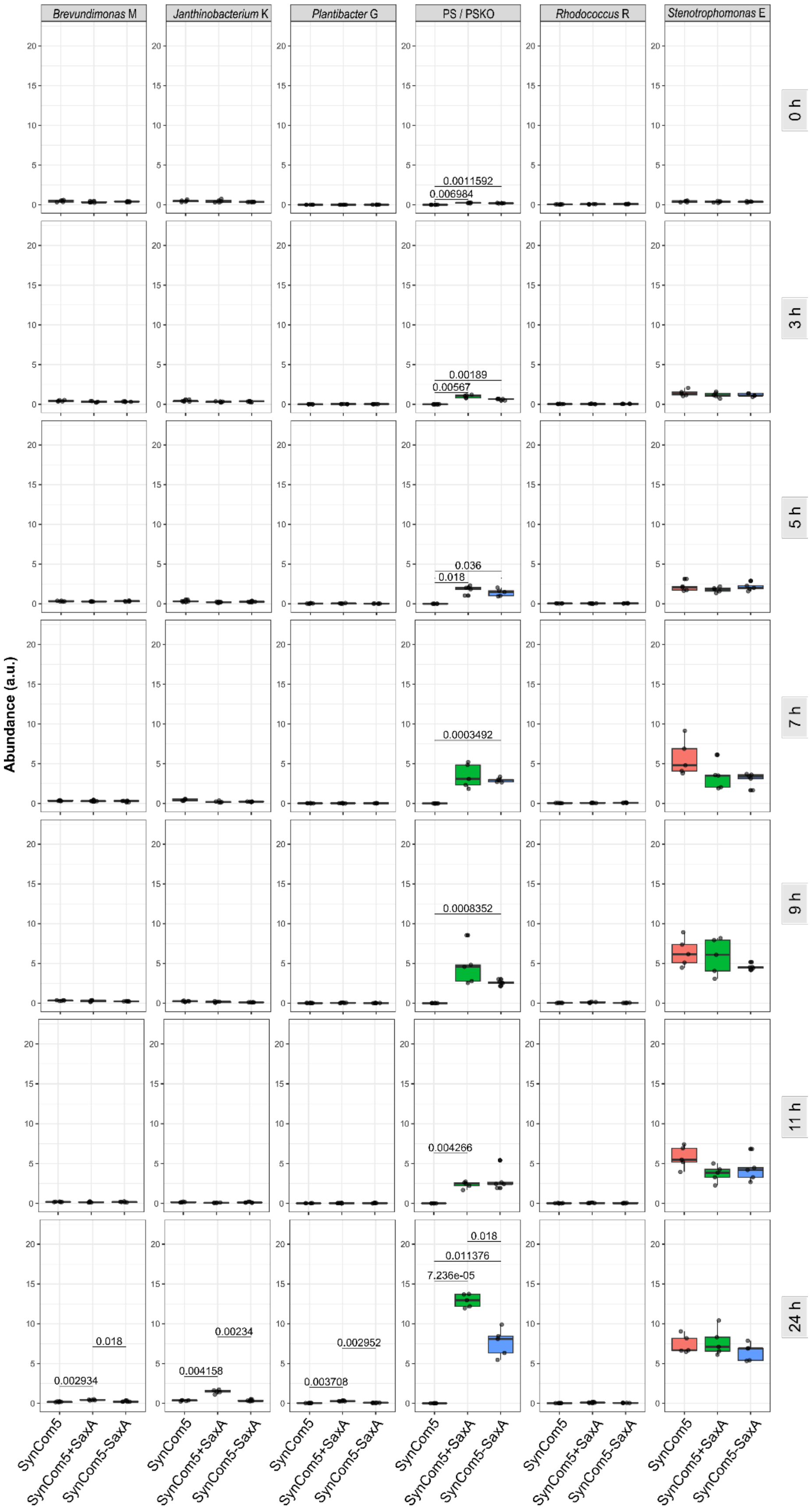


**Supplementary Figure 6: Abundance of SynCom taxa at each sampling timepoint.** Normalized abundance of each SynCom taxon over time at 0, 3, 5, 7, 9, 11 and 24 h. Dots show individual samples (n=5 technical replicates), boxes depict the median +/- interquartile range, whiskers show +/- 1.5x interquartile range. Differences between treatments were assessed using t-tests, only significant comparisons (p<0.05) are shown (adjusted p-values, using Bonferroni method).

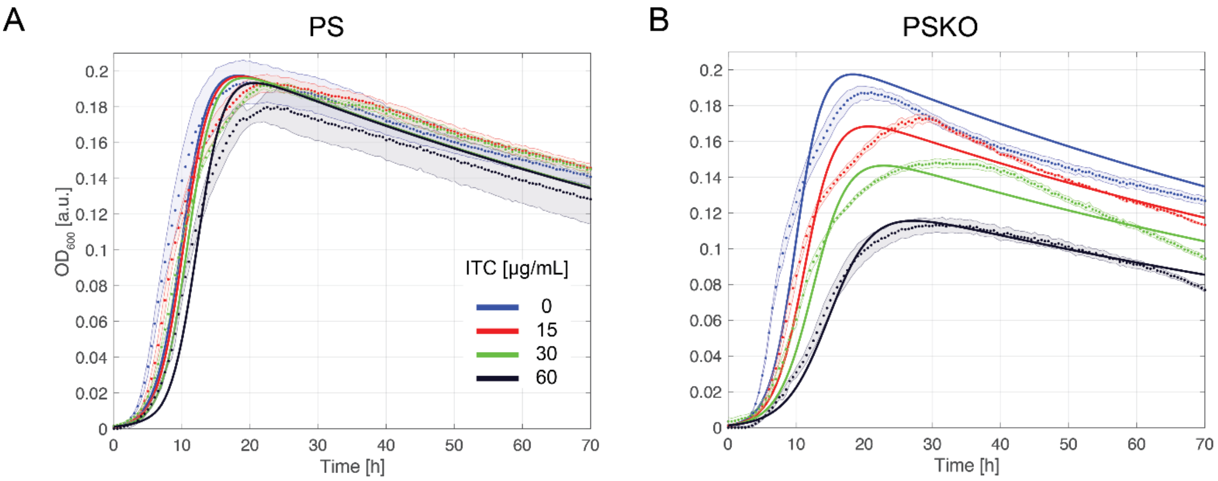


**Supplementary Figure 7: Fit of PS and PSKO with a single nutrient model to the data. (A)** and **(B)** show the fit of the model (solid lines) with a single nutrient assimilation (Equations 1-3 in Fig. 1B in the main text) to the data (averages and standard deviations over three replicates as dots and shaded regions, respectively) for several ITC concentrations.

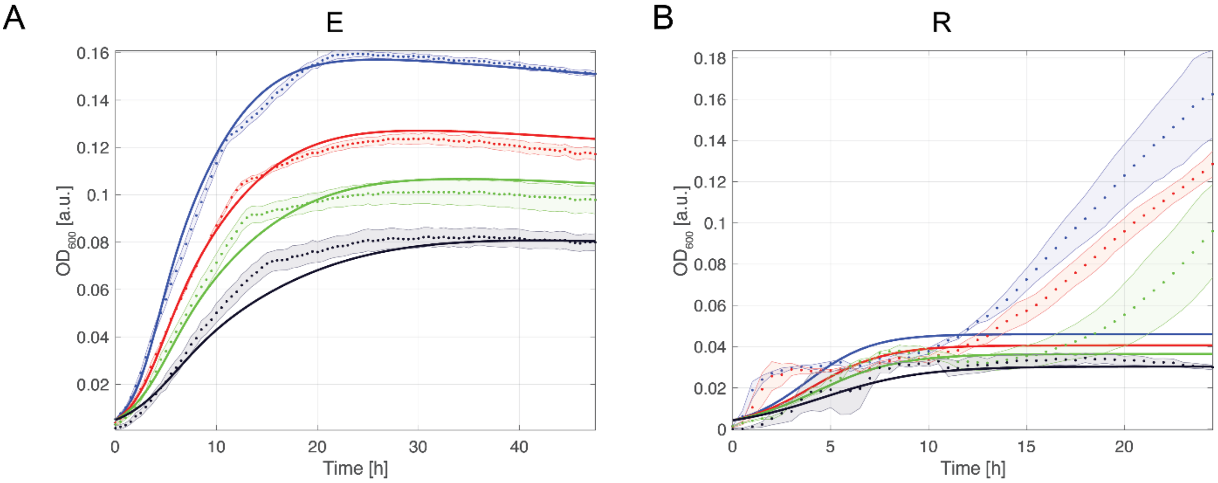


**Supplementary Figure 8: Alternative models for commensal E and R.** (**A)** Fit of an alternative model with diauxic shift between two nutrients for the growth of the commensal E. The improvement compared to a single nutrient of Fig. 5B in the main text is not enough to justify a more complex mathematical model with four more parameters. **(B)** Fit of an alternative model with a single nutrient for the commensal R. Here, the single nutrient even fails to catch the trend of the data, therefore a two-nutrient model was used.

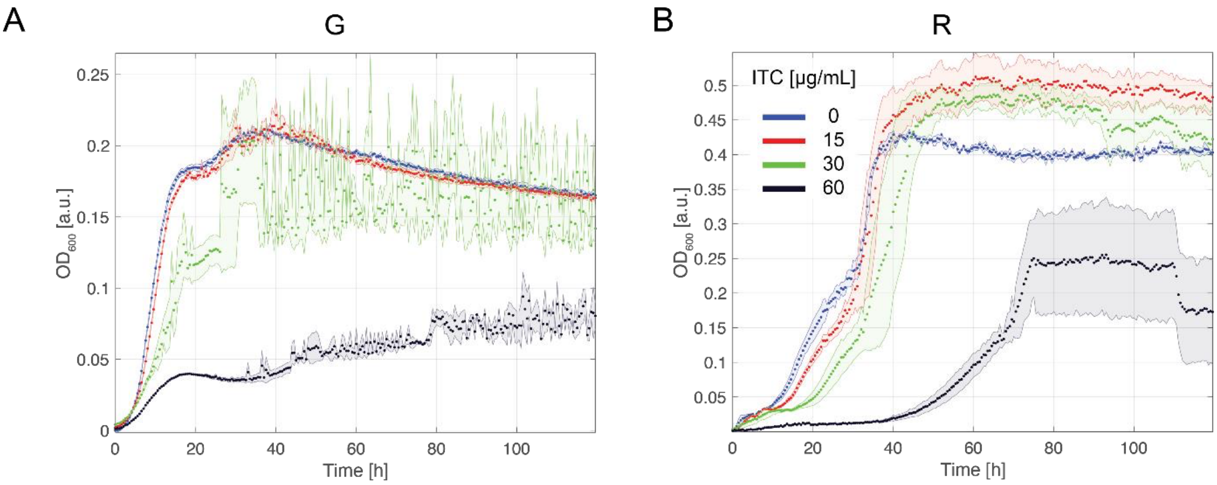


**Supplementary Figure 9: R and G form biofilms and/or aggregates which falsify the OD readings.** To check the formation of biofilm and aggregates for the commensal G and R beyond a visual inspection (panel A and B, respectively), we growth them for a longer time (120 h). We verified that their OD dynamics present several complex phases, large oscillations, and high standard deviations, possible indication of inhomogeneous densities in the media induced by bacterial clumps, especially for higher ITC concentrations.

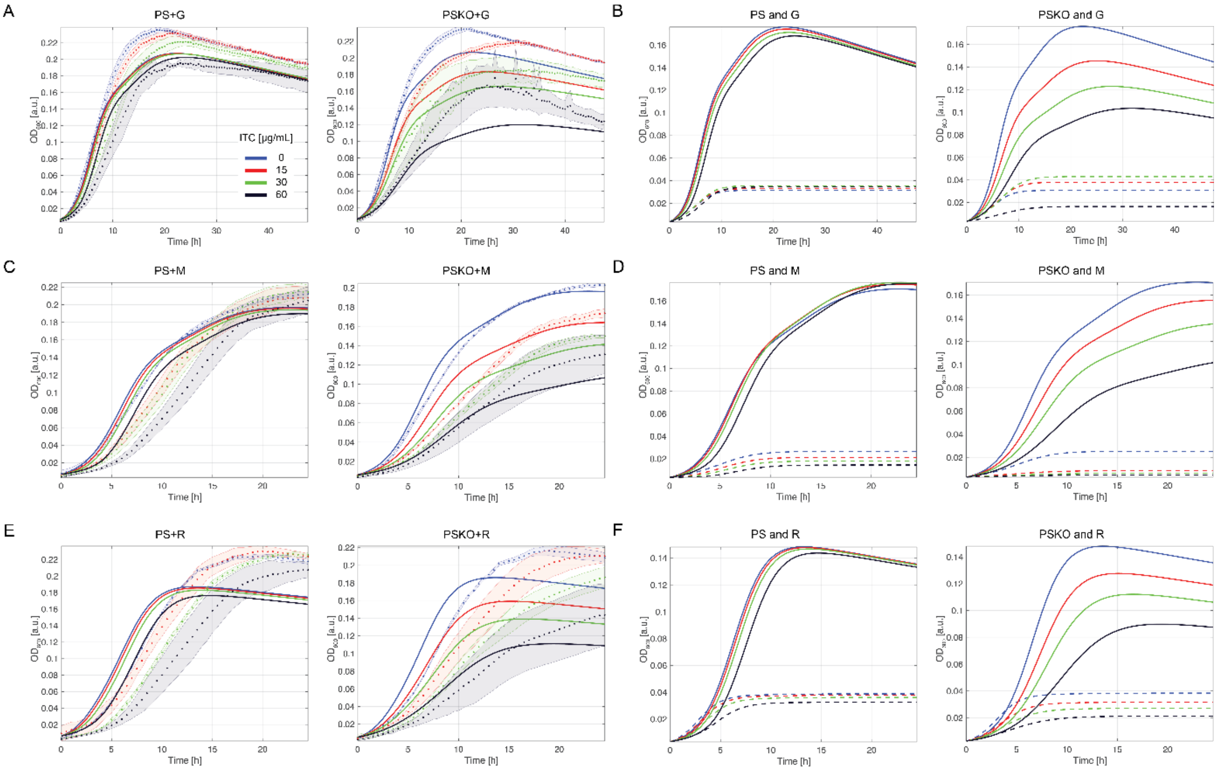


**Supplementary Figure 10: Pairwise predictions of PS(KO) and commensals G, M and R. (A, C, and E)** The model prediction for the total biomass dynamics ($OD_{600}$) of the pairwise culture starting with a same amount of either PS or PSKO and commensal G, M, and R for different ITC concentrations are shown by the solid lines and compared to the experimental data (dots, averages of three technical replicates, standard deviations as shaded region), in panel A, C, and E, respectively. **(B, D, and F)** Separate dynamics of the biomass of either PS or PSKO and the commensal G, M, and R. Solid lines show the $OD_{600}$ of PS or PSKO, and dashed lines the $OD_{600}$ of the commensal. The total biomass is mostly determined by the pathogen, and the commensal biomass reaches a maximum of one fourth of the pathogen biomass.

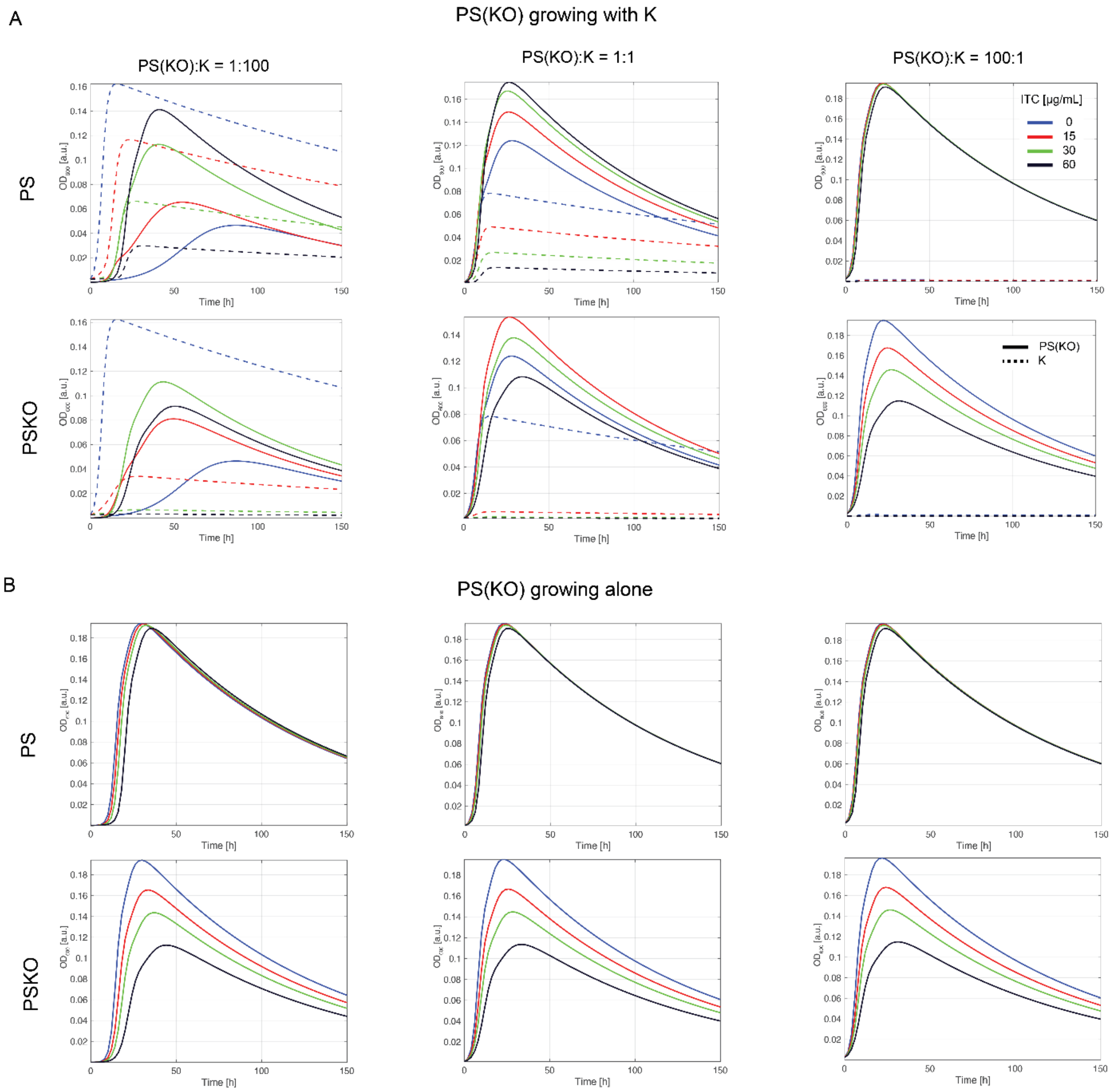


**Supplementary Figure 11: $OD_{600}$ dynamics of pathogens PS and PSKO growing in the absence or presence of commensal K for several initial PS(KO):K ratios. (A)** $OD_{600}$ dynamics of either PS or PSKO (solid lines, top and bottom row, respectively) and K (dashed lines) for several ITC concentrations and three initial PS(KO):K ratio (1:100, 1:1, 100:1, first, second and third column, respectively). Panel **(B)** shows the same conditions but for monocultures of either PS or PSKO, in the absence of commensal K. Predictions in A and B illustrate how commensal K influences growth of PS or PSKO, which is formalized in the suppression index.

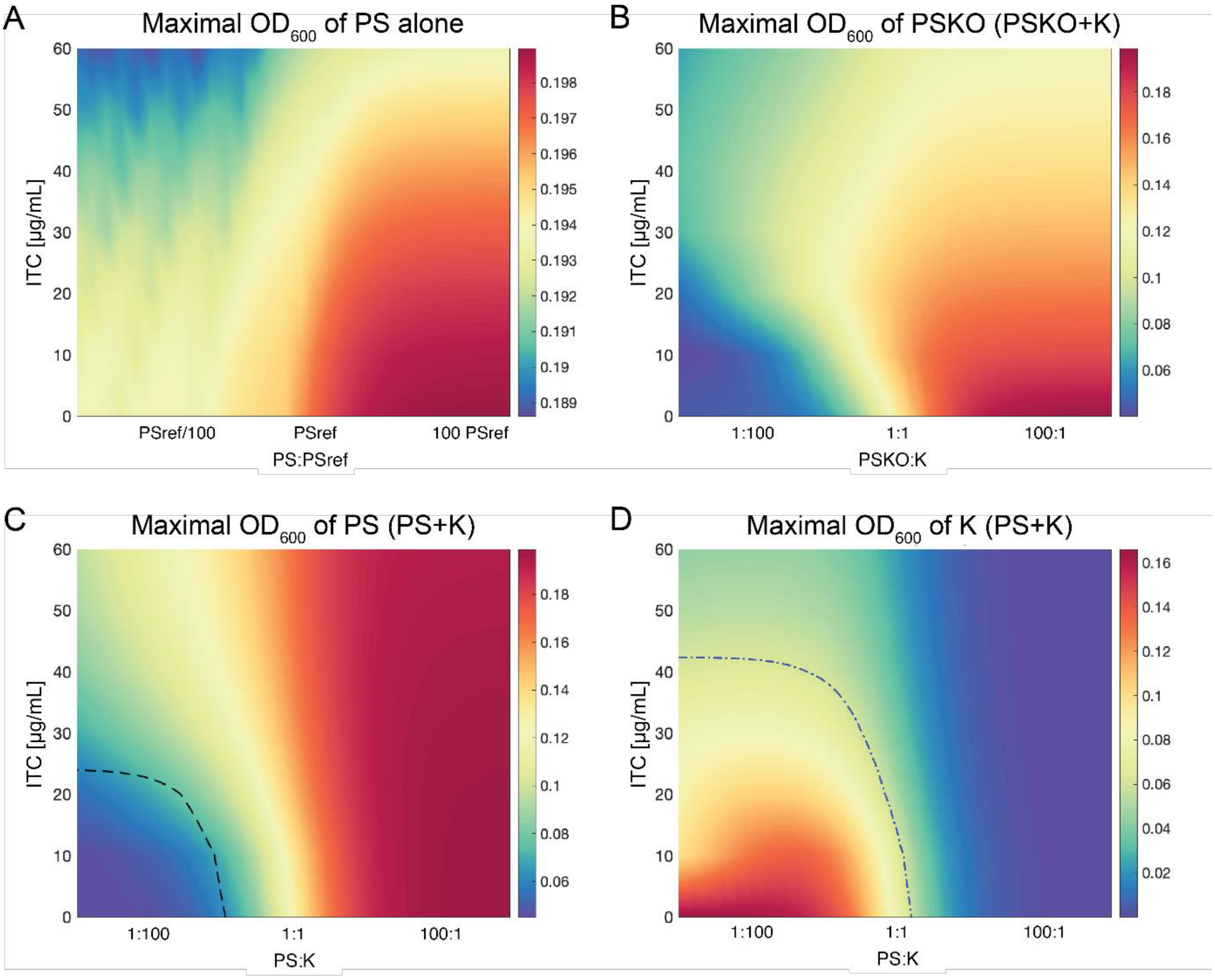


**Supplementary Figure 12: Heatmaps of maximal $OD_{600}$ used to calculate the indices.** Summary of the maximal $OD_{600}$ extracted from the dynamics of the pathogen and commensal biomass as explained in Suppl. Text 3. Panel A shows the maximal $OD_{600}$ attained by the monocultures of PS. PSref indicate the reference initial value of PS ($OD_{600}$ = 0.0025). Panel B shows the maximal $OD_{600}$ of PSKO attained by the pairwise culture of PSKO and K. Panels C and D show the maximal $OD_{600}$ of PS and K attained by the pairwise culture of PS and K. The dashed black line in C and the dot-dashed blue line in D indicate the conditions where the maximal $OD_{600}$ reaches 0.06 (isoline), respectively.

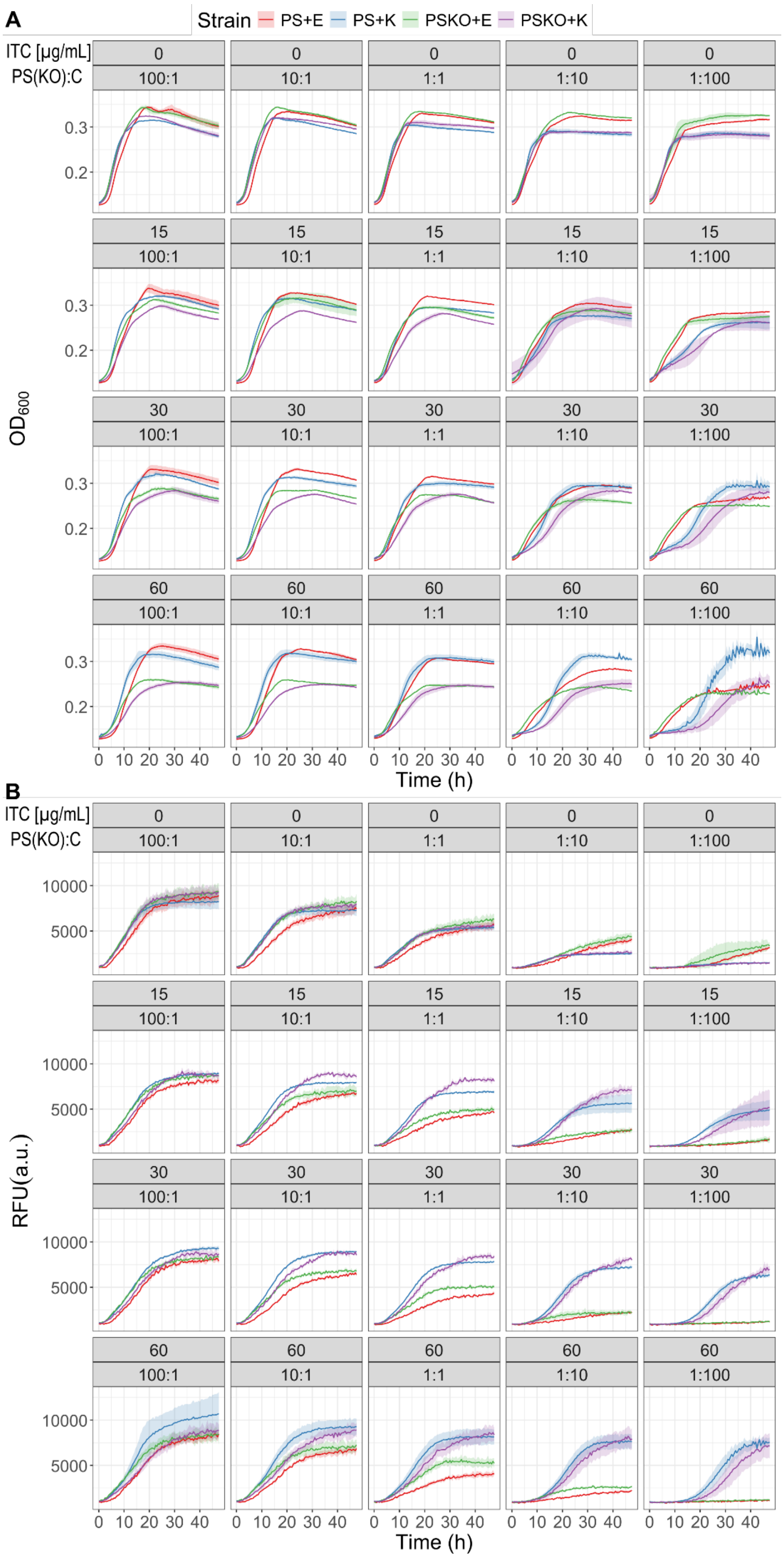


**Supplementary Figure 13: Growth of different initial ratios of PS/PSKO paired with a commensal.** Growth curves of $OD_{600}$ (A) and red fluorescence (B) of different initial mixtures of PS or PSKO tagged with mScarlet-I mixed with either the commensals E or K, or PSKO as control. (PS(KO):C ratios ranged from 100:1 to 1:100. The total initial $OD_{600}$ was kept at 0.4. The average of three technical replicates is shown (solid line) with standard deviation (shaded area).

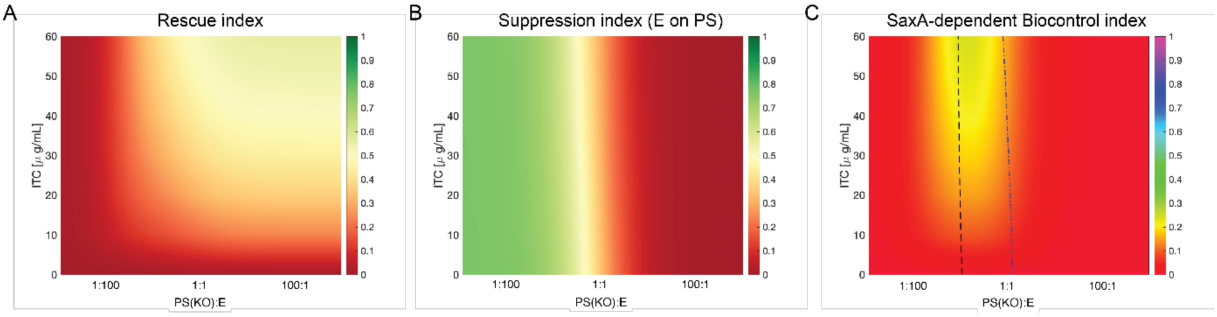


**Supplementary Figure 14: Indices for the most ITC-resistant commensal strain E.** Panels A, B, and C: Heat maps of the rescue, suppression, and SaxA-mediated biocontrol index, varying the initial PS(KO):E ratio and for several ITC concentrations. The black dashed and blue dot-dashed lines in C represent where the maximal $OD_{600}$ of PS and E are 0.06, respectively. Areas under the curves in C define the SaxA-mediated biocontrol region where commensal growth benefits more from SaxA than PS, the color illustrates the strength of the potential effect. It is worth noticing that the rescue index does not distinguish between a case when the rescue fails (e.g. ineffective SaxA-dependent degradation of ITC) and a case when there is no need to rescue because the commensal strain presents a low ITC sensitivity (e.g. as for E). To distinguish between these conditions, we introduce in Suppl. Fig. 15 a new index that quantifies the "need-to-be rescued" as ITC-sensitivity index.

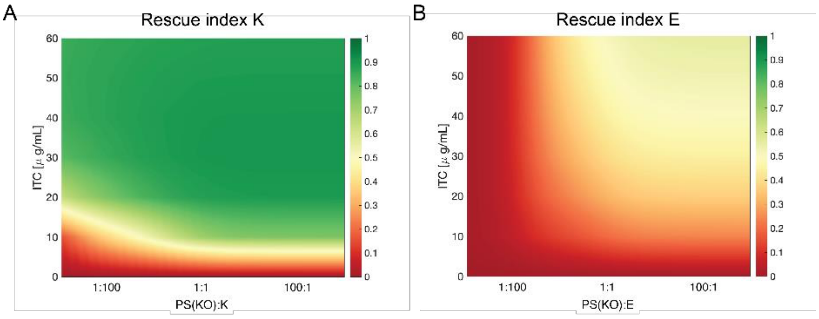


C

$$\text{ITC} - \text{sensitivity index} = 1 - \frac{\text{Max OD}(C \text{ with PSKO})}{\text{Max OD}(C \text{ with PSKO}, [ITC]{=}0)}$$

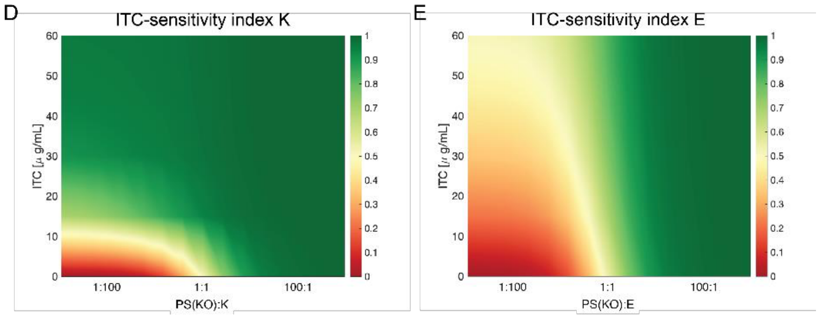


**Supplementary Figure 15: ITC-sensitivity index for commensal strain K and E.** Panels A and B show the rescue indices of strains K and E, which were already presented in Fig. 7B (main text) and Suppl. Fig. 14A. Importantly, the rescue index does not provide information on the underlying reason for the rescue outcome. For example, the red region in the rescue index of strain E indicates that E is not rescued in most conditions. However, this is not due to an inefficient SaxA-dependent ITC degradation, but rather to the fact that strain E intrinsically does not require rescue, owing to its low ITC sensitivity. To disentangle these possibilities, we introduce in Panel C the ITC-sensitivity index, which quantifies the sensitivity of each strain to 4MSOB-ITC on a scale from 0 to 1. The index compares the max OD attained by a commensal strain C grown in the presence of PSKO to the same condition but in the absence of ITC (C+PSKO and [ITC]=0). In this representation, green regions denote high ITC sensitivity and thus a potential requirement for rescue, whereas red regions correspond to ITC insensitivity and therefore no need for rescue of the commensal strain. Panel D shows that strain K is highly ITC-sensitive under most conditions, except at low [ITC] and high cell numbers, consistent with the pattern observed in the rescue index. By contrast, Panel E clarifies why the rescue index of strain E shows an apparent lack of rescue: strain E is largely ITC-insensitive across the explored parameter space and therefore does not require rescue. Indeed, E grows robustly even at high [ITC] (see Suppl. Fig. 16).

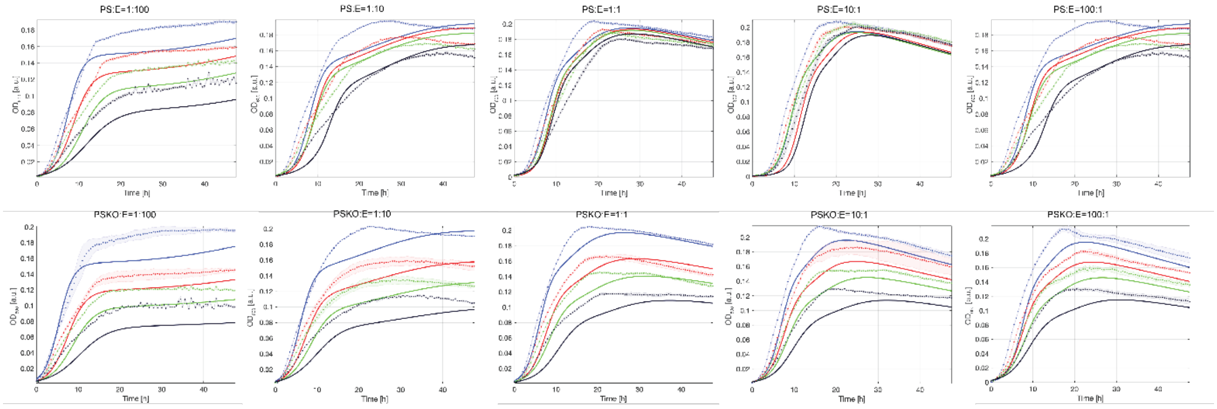

**Supplementary Figure 16: Pairwise prediction for PS(KO):E at different initial ratios.** Model predictions (solid lines) and experimental measurements (average as dots and shaded regions as standard deviations of three independent replicates) for the $OD_{600}$ of the pairwise cultures PS:E and PSKO:E, in the first and second row, respectively, and for different PS(KO):E ratios (1:100, 1:10, 1:1, 10:1, 100:1).

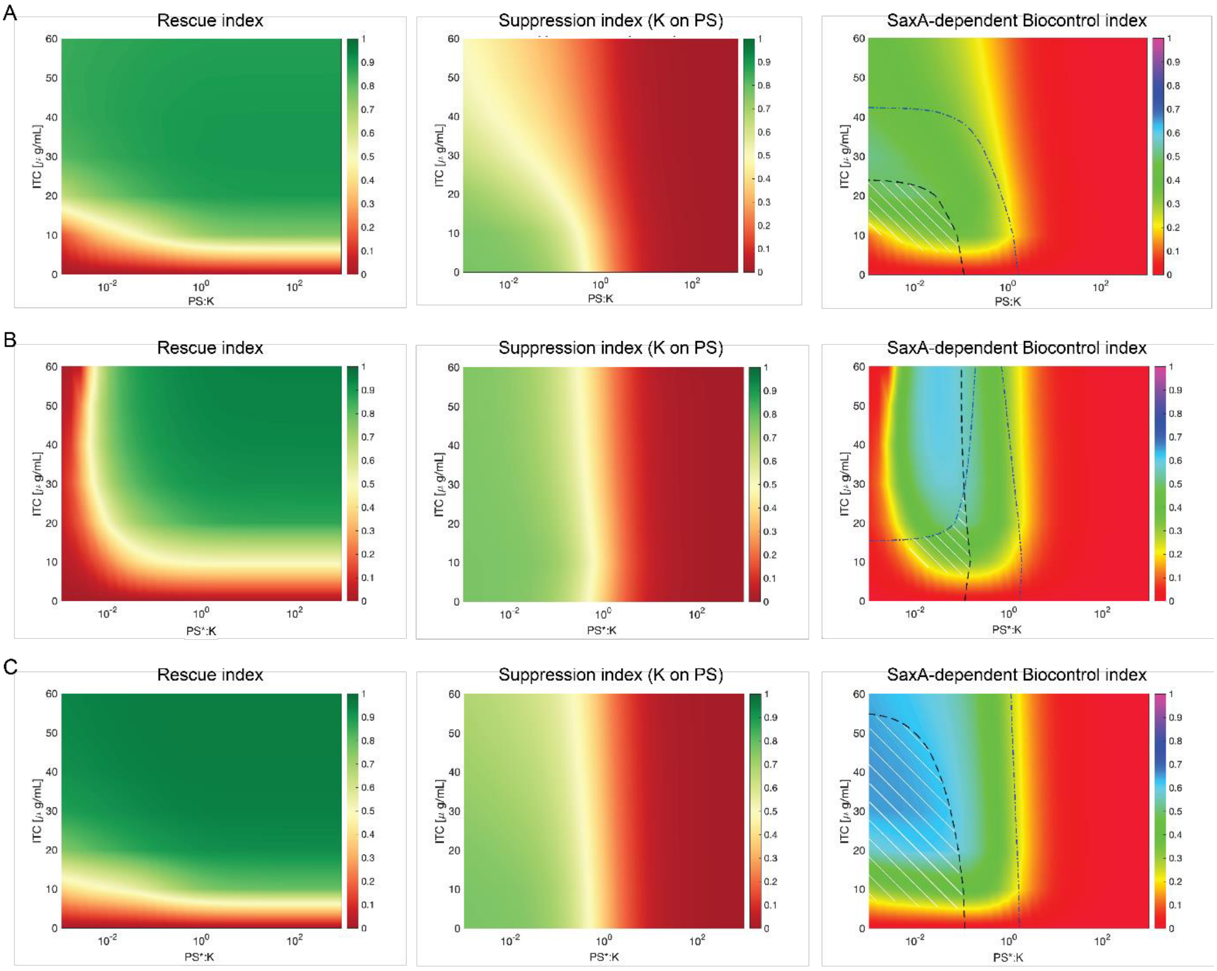


**Supplementary Figure 17: Indices for a hypothetical pathogen PS* either with increased ITC sensitivity or with increased ITC degradation rate.** Heat maps of the rescue, suppression, and SaxA-mediated biocontrol index for the pathogen PS (A), for hypothetical pathogen PS* with either tenfold higher ITC sensitivity (B, corresponding to a tenfold smaller value of the parameter $\theta$ in the mathematical model, Equation 8 in Suppl. Text 2) or with a tenfold increased ITC degradation rate (C, corresponding to tenfold larger value of $\lambda$ in the mathematical model, Equation 12 of Suppl. Text 2), varying the initial PS(KO)*:K ratio and for several ITC concentrations. The black dashed and blue dot-dashed lines in C represent where the maximal $OD_{600}$ of PS* and K are 0.06, respectively. The white hatched region in the biocontrol index defines the region where the SaxA-mediated biocontrol is maximal and both PS cells are not too abundant, and K cells are present with a sensitive number.

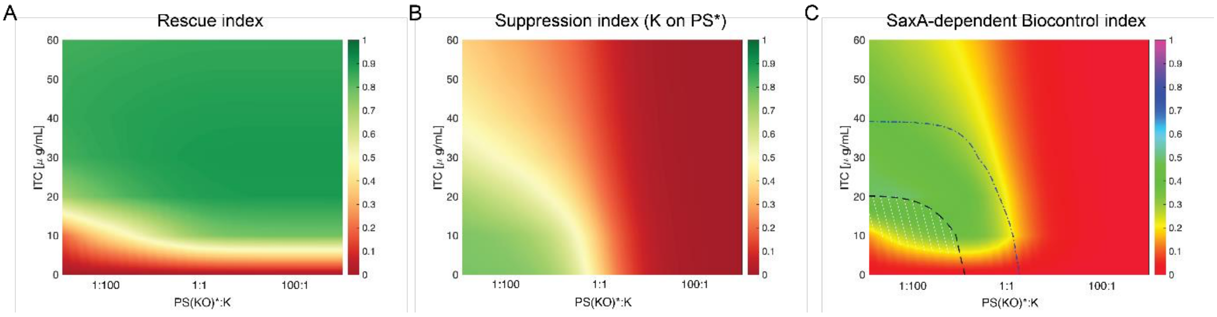


**Supplementary Figure 18: Indices for PS with lower ITC sensitivity.** Panels A, B, and C: Heat maps of the rescue, suppression, and SaxA-mediated biocontrol index for a pathogen PS* obtained by lowering the ITC sensitivity of PS (corresponding to tenfold larger K in the mathematical model, Equation 8 of Suppl. Text 2) and varying the initial PS(KO)*:K ratio and for several ITC concentrations. The black dashed and blue dot-dashed lines in C represent where the maximal $OD_{600}$ of PS* and K are 0.06, respectively. Areas under the curves in C define the SaxA-mediated biocontrol region where commensal growth benefits more from SaxA than PS, the color illustrates the strength of the potential effect.

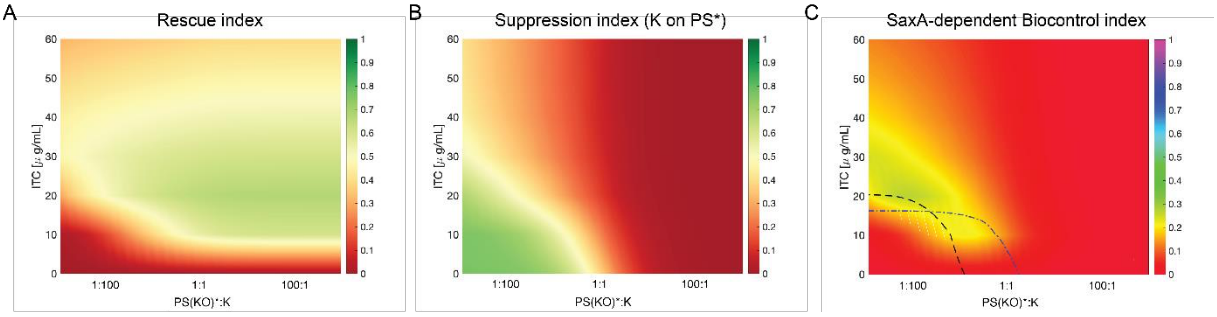


**Supplementary Figure 19: Indices for PS with lower ITC degradation rate.** Heat maps for the indices, as described in Suppl. Fig. 18, for a pathogen PS* obtained by lowering the ITC degradation rate of PS (corresponding to tenfold smaller $\lambda$ in the mathematical model, Equation 12 of Suppl. Text 2).

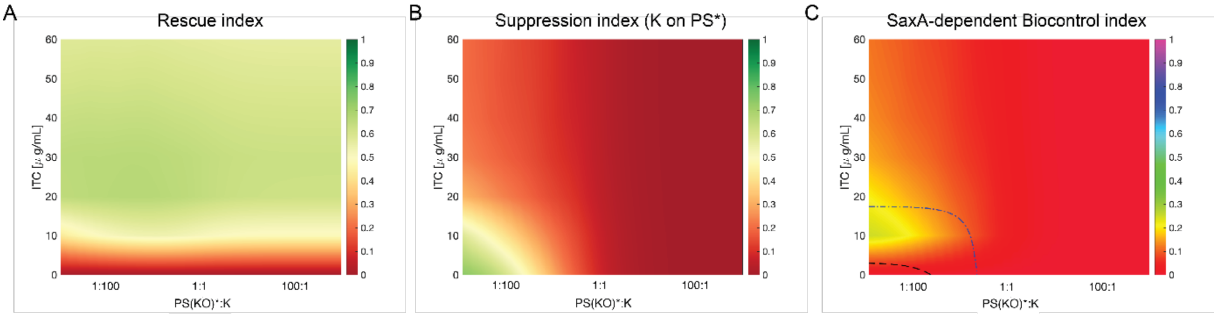


**Supplementary Figure 20: Indices for PS with higher growth rate.** Heat maps for the indices, as described in Suppl. Fig. 18, for a pathogen PS* obtained by increasing the growth rate of PS (corresponding to twofold larger $\mu_1$ in the mathematical model, Equation 8 of Suppl. Text 2).

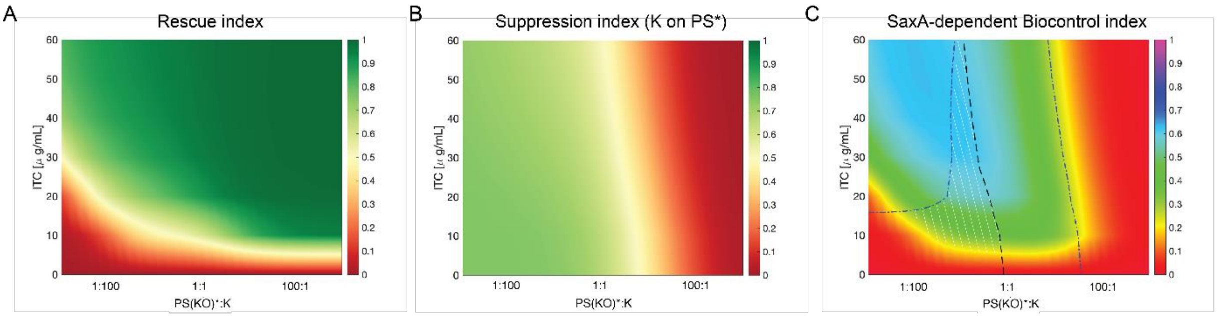


**Supplementary Figure 21: Indices for PS with lower growth rate.** Heat maps for the indices, as described in Suppl. Fig. 18, for a pathogen PS* obtained by lowering the growth rate of PS (corresponding to twofold smaller $\mu_1$ in the mathematical model, Equation 8 of Suppl. Text 2). Potential plant beneficial regions are in the lower left corner below the blue and black lines.

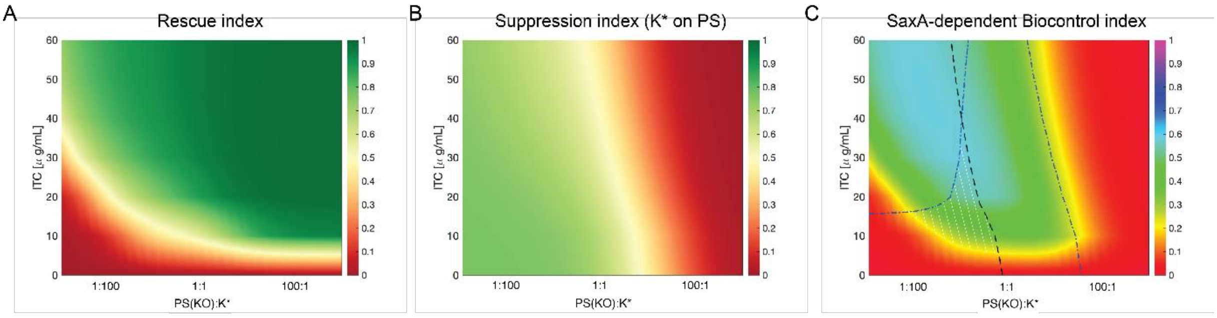


**Supplementary Figure 22: Indices for K with higher growth rate.** Heat maps for the indices, as described in Suppl. Fig. 18, for a pathogen K* obtained by increasing the growth rate of K (corresponding to twofold larger $\mu_C$ in the mathematical model, Equation 9 of Suppl. Text 2). Potential plant beneficial regions are in the lower left corner below the blue and black lines.

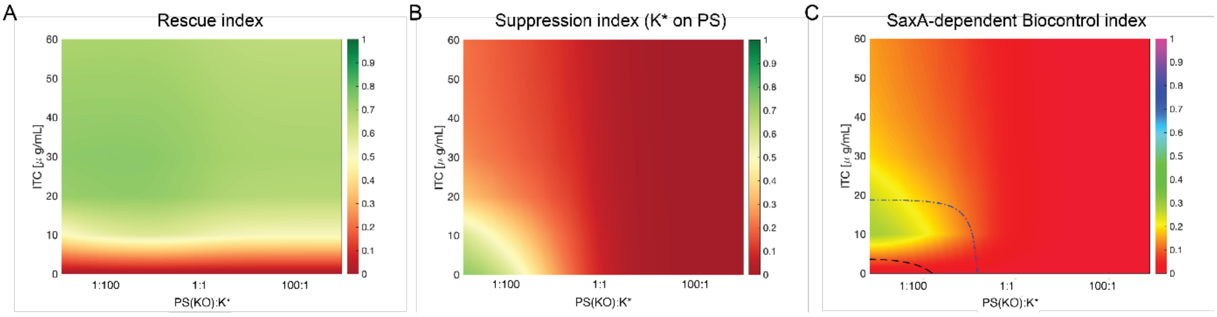


**Supplementary Figure 23: Indices for K with lower growth rate.** Heat maps for the indices, as described in Suppl. Fig. 18, for a pathogen K* obtained by lowering the growth rate of K (corresponding to twofold smaller $\mu_C$ in the mathematical model, Equation 9 of Suppl. Text 2).

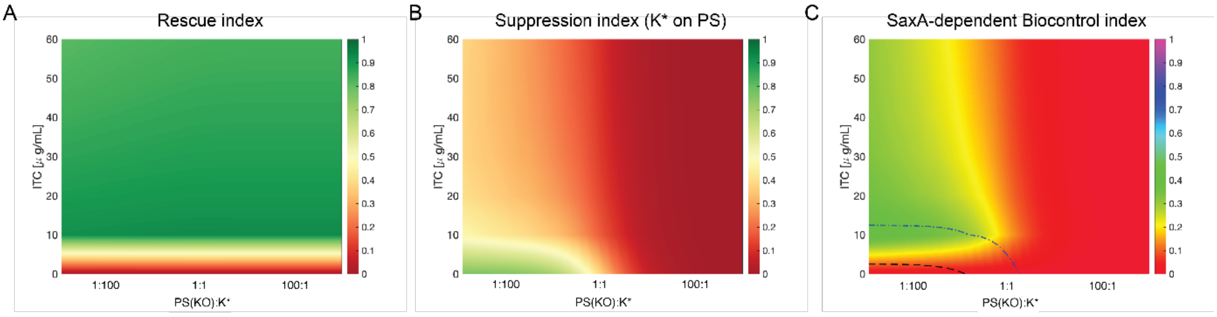


**Supplementary Figure 24: Indices for K when more susceptible to ITC.** Heat maps for the indices, as described in Suppl. Fig. 18, for a pathogen K* obtained by increasing the susceptibility to ITC (corresponding to tenfold smaller $K_C$ in the mathematical model, Equation 9 of Suppl. Text 2).

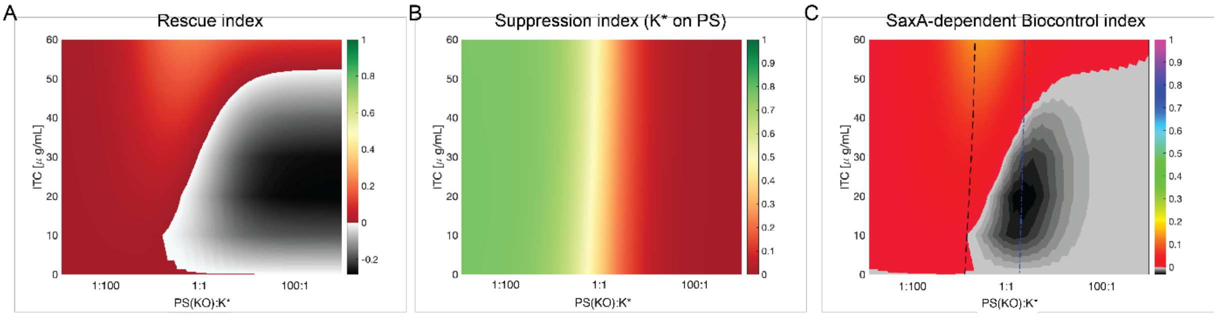


**Supplementary Figure 25: Indices for K when less susceptible to ITC.** Heat maps for the indices, as described in Suppl. Fig. 18, for a pathogen K* obtained by lowering the susceptibility to ITC (corresponding to tenfold larger $K_C$ in the mathematical model, Equation 9 of Suppl. Text 2). Since K* is almost insensitive to ITC, it does not need to be rescued by the ITC-degrader PS. For some low ranges of ITC concentration, K* grows better in the presence of PSKO than PS, thus leading to negative values of the rescue index (indicated by the gray regions). Therefore, unlike all the cases here analyzed, the rescue index and therefore the biocontrol index can be negative (gray regions).

### Supplementary Tables

**Supplementary Table 1: Characteristics of all potential SynCom members.** Bacterial strains were isolated from *A. thaliana* leaves. n.t. = not tested.

| Letter | Phylum | Genus | ID | Reference (origin) | Growth in R2A broth in 24-48 h | 4MSOB-ITC degradation | Chosen for SynCom? | Why not chosen? |
|---|---|---|---|---|---|---|---|---|
| A | Firmicutes | *Bacillus* | A | (Jose *et al.*, 2024) | Yes | Yes | no | Degrades 4MSOB-ITC |
| B | Gammaproteobacteria | *Pantoea* | Pa7 | (Unger *et al.*, 2024) | Yes | Yes | no | Degrades 4MSOB-ITC |
| C | Bacteroidetes | *Flavobacterium* | AR-4198 | (Mayer *et al.*, 2025) | No | n.t. | no | Slow growth |
| D | Actinobacteria | *Curtobacterium* | 5B7 | (Mayer *et al.*, 2025) | Yes | No | no | No more Actinobacteria needed |
| E | Gammaproteobacteria | *Stenotrophomonas* | S8 | (Unger *et al.*, 2024) | Yes | No | yes | |
| F | Actinobacteria | *Microbacterium* | M5 | (Unger *et al.*, 2024) | Yes | No | no | No more Actinobacteria needed |
| G | Actinobacteria | *Plantibacter* | 2H11-2 | (Mayer *et al.*, 2025) | Yes | No | yes | |
| H | Gammaproteobacteria | *Massilia* | 1F9 | (Mayer *et al.*, 2025) | n.t. | n.t. | no | Contaminated glycerol stock |
| I | Alphaproteobacteria | *Sphingomonas* | 1G12-1 | (Mayer *et al.*, 2025) | No | Yes | no | Slow growth, degrades 4MSOB-ITC |
| J | Gammaproteobacteria | *Xanthomonas* | 4E10 | (Mayer *et al.*, 2025) | Yes | No | no | Has sax efflux pumps |
| K | Betaproteobacteria | *Janthinobacterium* | J4 | (Unger *et al.*, 2024) | Yes | No | yes | |
| L | Gammaproteobacteria | *Pseudomonas* | LR-512C6 | (Mayer *et al.*, 2025) | Yes | n.t. | no | not to differentiate from Ps by 16S |
| M | Alphaproteobacteria | *Brevundimonas* | 7B5 | (Mayer *et al.*, 2025) | Yes | No | yes | |
| N | Gammaproteobacteria | *Pseudomonas* | 3D9 | (Mayer *et al.*, 2025) | Yes | Yes | yes | |
| O | Gammaproteobacteria | *Pseudomonas* | 3D9Δ*saxA* | (Unger *et al.*, 2025) | Yes | No | yes | |
| P | Betaproteobacteria | *Acidovorax* | 4E11-1 | (Mayer *et al.*, 2025) | No | n.t. | no | Slow growth |
| Q | Alphaproteobacteria | *Rhizobium* | 7E11-2 | (Mayer *et al.*, 2025) | No | No | no | Slow growth |
| R | Actinobacteria | *Rhodococcus* | 6G8 | (Mayer *et al.*, 2025) | Yes | No | yes | |
| S | Bacteroidetes | *Chryseobacterium* | LR-58G5 | (Mayer *et al.*, 2025) | Yes | Yes | no | Degrades 4MSOB-ITC |

**Supplementary Table 2: Details of the analysis of 4MSOB-ITC, 4MSOB-amine and 4MSOB-ITC-GSH by LC-MS.** Compounds were measured using an Agilent HPLC 1200/API3200 (AB SCIEX) instrument in positive ionisation mode. Abbreviations are: Q1, selected *m/z* of the first quadrupole; Q3, selected *m/z* of the third quadrupole; RT, retention time; DP, declustering potential (V); and CE, collision energy (V).

| Q1 | Q3 | RT (min) | compound | DP | CE |
|---|---|---|---|---|---|
| 136 | 72 | 0.5 | 4MSOB-amine | 26 | 17 |
| 178 | 114 | 2.6 | 4MSOB-ITC | 60 | 13 |
| 485 | 179 | 2.0 | 4MSOB-ITC-GSH | 51 | 29 |

**Supplementary Table 3: Values of the parameters used for generating the plot of Fig. 3 C and D in the main text.** The values of the parameters are used in Equations 1-3 of Fig. 1B to simulate the dynamics of biomass, substrate and ITC.

| Parameter [units] | Value |
|---|---|
| $\mu$ [$h^{-1}$] | 0.056 |
| $\theta$ [a.u.] | 17.11 |
| $\lambda$ [$h^{-1}$] | 0.6 |
| $\theta_S$ [g/mL] | 20 |
| $Y$ [g/(mL a.u.)] | 3.5 |
| $\theta_{ITC}$ [$\mu$g/mL] | 45.46 |
| $\delta$ [$h^{-1}$] | $1{\cdot}10^{-6}$ |

**Supplementary Tables 4: Values of the parameters obtained by fitting the monocultures of pathogens and commensals.** The tables show the values of the parameters obtained by fitting Equations in Suppl. Text 2 to the monocultures of pathogens PS and PSKO, and commensals E, G, K, M, and R.

**PS(KO)**

| Parameter [units] | Mean value | Standard deviation |
|---|---|---|
| $\mu$ [$h^{-1}$] | 0.0325 | 0.0059 |
| $\theta$ [a.u.] | 99.6693 | 0.4736 |
| $\lambda$ [$h^{-1}$] | 0.5349 | 0.0907 |
| $\theta_S$ [g/mL] | 94.8442 | 17.5521 |
| $Y$ [g/(mL a.u.)] | 3.1723 | 0.0270 |
| $\theta_{ITC}$ [$\mu$g/mL] | 5.3960 | 3.7518 |
| $\delta$ [$h^{-1}$] | $2.636 \cdot 10^{-6}$ | $1.0571 \cdot 10^{-9}$ |
| $\mu_2$ [$h^{-1}$] | 0.0108 | 0.0064 |
| $\theta_2$ [a.u.] | 218.23 | 125.97 |
| $Y_2$ [g/(mL a.u.)] | 7.2867 | 0.1328 |
| $\theta_N$ [g/mL] | 0.0152 | 0.0011 |

**E**

| Parameter [units] | Mean value | Standard deviation |
|---|---|---|
| $\mu$ [$h^{-1}$] | 0.0724 | 0.0182 |
| $\theta$ [a.u.] | 53.4802 | 0.4281 |
| $\theta_S$ [g/mL] | 234.37 | 59.07 |
| $Y$ [g/(mL a.u.)] | 3.3846 | 0.0067 |
| $\delta$ [$h^{-1}$] | $1 \cdot 10^{-8}$ | $1.0643 \cdot 10^{-8}$ |

**G**

| Parameter [units] | Mean value | Standard deviation |
|---|---|---|
| $\mu$ [$h^{-1}$] | 0.1001 | 0.1966 |
| $\theta$ [a.u.] | 60.3020 | 0.3973 |
| $\theta_S$ [g/mL] | 496.70 | 976.15 |
| $Y$ [g/(mL a.u.)] | 2.5959 | 0.0035 |
| $\delta$ [$h^{-1}$] | $1 \cdot 10^{-8}$ | $9.4007 \cdot 10^{-9}$ |
| $m$ | 5 | - |

**K**

| Parameter [units] | Mean value | Standard deviation |
|---|---|---|
| $\mu$ [$h^{-1}$] | 0.0300 | 0.0033 |
| $\theta$ [a.u.] | 10.0941 | 0.0520 |
| $\theta_S$ [g/mL] | 88.4704 | 9.9577 |
| $Y$ [g/(mL a.u.)] | 2.9937 | 0.0139 |
| $\delta$ [$h^{-1}$] | $8.7739 \cdot 10^{-7}$ | $1.4925 \cdot 10^{-8}$ |
| $m$ | 2 | 0.02 |

**M**

| Parameter [units] | Mean value | Standard deviation |
|---|---|---|
| $\mu$ [$h^{-1}$] | 0.0698 | 0.0192 |
| $\theta$ [a.u.] | 11.1501 | 0.1883 |
| $\theta_S$ [g/mL] | 367.9310 | 103.6806 |
| $Y$ [g/(mL a.u.)] | 3.8752 | 0.0558 |
| $\delta$ [$h^{-1}$] | $1 \cdot 10^{-8}$ | $2.6746 \cdot 10^{-7}$ |

**R**

| Parameter [units] | Mean value | Standard deviation |
|---|---|---|
| $\mu$ [$h^{-1}$] | 0.0125 | 0.0001 |
| $\theta$ [a.u.] | 61.2687 | 0.1573 |
| $\theta_S$ [g/mL] | 26.7859 | 0.0406 |
| $Y$ [g/(mL a.u.)] | 12.7652 | 0.0442 |
| $\delta$ [$h^{-1}$] | $1 \cdot 10^{-8}$ | $3.6336 \cdot 10^{-8}$ |
| $\mu_2$ [$h^{-1}$] | 0.0147 | 0.0001 |
| $\theta_2$ [a.u.] | 108.93 | 0.3312 |
| $Y_2$ [g/(mL a.u.)] | 0.0117 | 0.0006 |
| $\theta_N$ [g/mL] | 0.0002 | $2.4211 \cdot 10^{-6}$ |

### Supplementary References

**Jose J, Teutloff E, Naseem S, Barth E, Halitschke R, Marz M, Agler MT. 2024.** Immunity and bacterial recruitment in plant leaves are parallel processes whose link shapes sensitivity to temperature stress. *bioRxiv*: 2024.06.10.598336.

**Mayer T, Teutloff E, Unger K, Lehenberger P, Agler MT. 2025.** Deterministic colonization arises early during the transition of soil bacteria to the phyllosphere and is shaped by plant–microbe interactions. *Microbiome* 13: 102.

**Unger K, Raza SAK, Mayer T, Reichelt M, Stuttmann J, Hielscher A, Wittstock U, Gershenzon J, Agler MT. 2024.** Glucosinolate structural diversity shapes recruitment of a metabolic network of leaf-associated bacteria. *Nature Communications* 15: 8496.

**Unger K, Ruiter R, Reichelt M, Gershenzon J, Agler MT. 2025.** *Pseudomonas* virulence factor SaxA detoxifies plant glucosinolate hydrolysis products, rescuing a commensal that suppresses virulence gene expression. *bioRxiv*: 2025.04.25.650564.

**ITC has the effect of modifying the maximal OD and thus the yield coefficient**

Bacterial growth is usually described by the assimilation of substrate $S$ that is transformed into biomass $B$ via an ordinary differential equation system

$$\frac{d[B(t)]}{dt} = \mu \frac{[S(t)]}{[S(t)] + \theta_S}[B(t)] \tag{1}$$

$$\frac{d[S(t)]}{dt} = -Y\frac{d[B(t)]}{dt} \tag{2}$$

where $Y$ represent the yield that converts the units of the substrate (e.g. $g/L$) into biomass units (e.g. $a.u.$ or cell/volume) and expresses how much nutrient amount is necessary to make a unit of biomass. Such equations obey to the mass conservation principle, so that their net sum is zero

$$\frac{d[B(t)]}{dt} + \frac{1}{Y}\frac{d[S(t)]}{dt} = 0 \tag{3}$$

expressing the fact that no substrate is lost in the assimilation into biomass. In the monocultures, we observed that the maximal OD is reduced as a function of the ITC concentration and we hypothesize that this leads to an impaired assimilation of the substrate. Therefore, we modified the ODE system including a loss term (Fig. 1), dependent on the ITC concentration, expressed as the proportion $\alpha$ (a number between 0 and 1) of substrate that is transformed into biomass. The ratio $1-\alpha$ is instead lost into $L$. Accounting for the lost substrate $L$, the ODE system becomes

$$\frac{d[B(t)]}{dt} = -\frac{\alpha}{Y}\frac{d[S(t)]}{dt} \tag{4}$$

$$\frac{d[L(t)]}{dt} = -\frac{1-\alpha}{Y}\frac{d[S(t)]}{dt} \tag{5}$$

$$\frac{d[S(t)]}{dt} = -Y\frac{d[B(t)]}{dt} - Y\frac{d[L(t)]}{dt}. \tag{6}$$

Neglecting the loss term as we are not interested in its dynamics in our system and imposing the Monod equation for the substrate assimilation, we obtain

$$\frac{d[B(t)]}{dt} = \alpha\mu \frac{[S(t)]}{[S(t)] + \theta_S}[B(t)] \tag{7}$$

$$\frac{d[S(t)]}{dt} = -Y\mu \frac{[S(t)]}{[S(t)] + \theta_S}[B(t)]. \tag{8}$$

From Equation 4, we notice the the system converts substrate into biomass via an effective yield

$$y = \frac{Y}{\alpha} \tag{9}$$

that depends on $\alpha$. We suppose then the ratio of nutrient assimilated into biomass $\alpha$ is a saturable function of the ITC concentration

$$\alpha(ITC) = \frac{\theta}{[ITC(t)] + \theta}, \tag{10}$$

so that ITC has the effective results of altering the yield $y$ in Equation 9

$$y = Y\frac{[ITC(t)] + \theta}{\theta} \tag{11}$$

that reduces to the normal yield in the absence of ITC and is smaller than $Y$ in the presence of ITC, expressing the diverting of the nutrient from the biomass pathway. We can finally rewrite the whole system by inserting the definition of $\alpha$ in Equations 7 and 8

$$\frac{d[B(t)]}{dt} = \mu\frac{[S(t)]}{[S(t)] + \theta_S}\frac{\theta}{[ITC(t)] + \theta}[B(t)] \tag{12}$$

$$\frac{d[S(t)]}{dt} = -\mu Y\frac{[S(t)]}{[S(t)] + \theta_S}[B(t)]. \tag{13}$$

Including the possibility to degrade ITC with a rate $\lambda$ by PS and a death rate term $\delta$ lead to the ODE of the main text

$$\frac{d[B(t)]}{dt} = \mu\frac{[S(t)]}{[S(t)] + \theta_S}\frac{\theta}{[ITC(t)] + \theta}[B(t)] - \delta[B(t)] \tag{14}$$

$$\frac{d[S(t)]}{dt} = -\mu Y\frac{[S(t)]}{[S(t)] + \theta_S}[B(t)] \tag{15}$$

$$\frac{d[ITC(t)]}{dt} = -\lambda\frac{[ITC(t)]}{[ITC(t)] + \theta_{ITC}}[B(t)],\ 0. \tag{16}$$

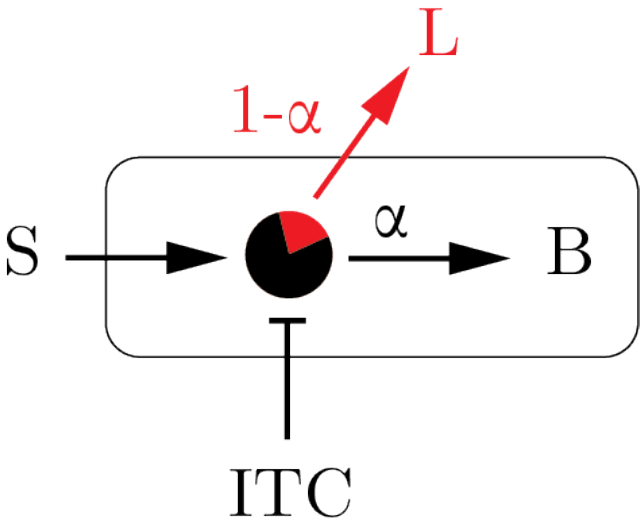


Figure 1: The substrate S is assimilated by the cell and a fraction $\alpha$ is transformed into biomass $B$, while the rest $1-\alpha$ is lost as $L$. ITC has the effect of increasing the coefficient $\alpha$ suppressing the biomass formation.

#### Extension of the model to two nutrients

The dynamics of the OD of the pathogens (PS, PSKO) show a change in the slope between 10 and 20 hours depending on the ITC concentration (Fig. 4 of the main text), probably indicating a switch in the nutrient utilization. To account for it, we modified our original model

$$\frac{d[B(t)]}{dt} = \mu\frac{[S(t)]}{[S(t)]+\theta_S}\frac{\theta}{[ITC(t)]+\theta}[B(t)] - \delta[B(t)] \quad (1)$$

$$\frac{d[S(t)]}{dt} = -\mu Y\frac{[S(t)]}{[S(t)]+\theta_S}[B(t)] \quad (2)$$

$$\frac{d[ITC(t)]}{dt} = -\lambda\frac{[ITC(t)]}{[ITC(t)]+\theta_{ITC}}[B(t)],\ 0 \quad (3)$$

including a diauxic shift from the first nutrient $S_1$ to a second nutrient $S_2$

$$\frac{d[B(t)]}{dt} = \left(\mu_1\frac{[S_1(t)]}{[S_1(t)]+\theta_{S1}} + \mu_2\frac{[S_2(t)]}{[S_2(t)]+\theta_{S2}}\frac{\theta_N}{[S_1(t)]+\theta_N}\right)\frac{\theta}{[ITC(t)]+\theta}[B(t)] - \delta[B(t)] \quad (4)$$

$$\frac{d[S_1(t)]}{dt} = -\mu_1 Y_1\frac{[S_1(t)]}{[S_1(t)]+\theta_{S1}}[B(t)] \quad (5)$$

$$\frac{d[S_2(t)]}{dt} = -\mu_2 Y_2\frac{[S_2(t)]}{[S_2(t)]+\theta_{S2}}\frac{\theta_N}{[S_1(t)]+\theta_N}[B(t)] \quad (6)$$

$$\frac{d[ITC(t)]}{dt} = -\lambda\frac{[ITC(t)]}{[ITC(t)]+\theta_{ITC}}[B(t)],\ 0 \quad (7)$$

where $\mu_i$, $\theta_{Si}$, and $Y_i$ are the maximal growth rate,the Monod constant, and the yield coefficient of the nutrient $i$, and $\theta_N$ the diauxic suppression constant indicating the concentration at which the first nutrient induces half maximal suppression on the assimilation of the second carbon source.

#### Extension of the model to pairwise culture

While growing together in pairwise culture, pathogen and commensal co-utilize the substrate. For simplicity, we suppose that they share the substrate $S_1$. If $B$ and $C$ represent the biomass of the pathogen and commensal, respectively, the system of ODE describing our model becomes

$$\frac{d[B(t)]}{dt} = \left(\mu_1 \frac{[S_1(t)]}{[S_1(t)] + \theta_{S1}} + \mu_2 \frac{[S_2(t)]}{[S_2(t)] + \theta_{S2}} \frac{\theta_N}{[S_1(t)] + \theta_N}\right) \frac{\theta}{[ITC(t)] + \theta}[B(t)] - \delta[B(t)] \tag{8}$$

$$\frac{d[C(t)]}{dt} = \mu_C \frac{[S_1(t)]}{[S_1(t)] + \theta_{SC}} \frac{\theta_C}{[ITC(t)] + \theta_C}[C(t)] - \delta[C(t)] \tag{9}$$

$$\frac{d[S_1(t)]}{dt} = -\mu_1 Y_1 \frac{[S_1(t)]}{[S_1(t)] + \theta_{S1}}[B(t)] - \mu_C Y_C \frac{[S_1(t)]}{[S_1(t)] + \theta_{SC}}[C(t)] \tag{10}$$

$$\frac{d[S_2(t)]}{dt} = -\mu_2 Y_2 \frac{[S_2(t)]}{[S_2(t)] + \theta_{S2}} \frac{\theta_N}{[S_1(t)] + \theta_N}[B(t)] \tag{11}$$

$$\frac{d[ITC(t)]}{dt} = -\lambda \frac{[ITC(t)]}{[ITC(t)] + \theta_{ITC}}[B(t)],\ 0 \tag{12}$$

where we rename the set of parameters with subscript $_C$ for the substrate consumption by the commensal.

**Including a Hill effect of ITC repression**

We noticed that in some cases, such as for the commensal K and G, the ITC repression does not follow the function

$$\frac{\theta}{[ITC(t)] + \theta} \tag{13}$$

but rather a sigmoidal function represented by the introduction of a Hill coefficient $m$

$$\frac{\theta^m}{[ITC(t)]^m + \theta^m} \tag{14}$$

that we use in our model and for fitting the single culture OD dynamics.

#### Development of the rescue, suppression, and biocontrol indexes

In the main text, we defined the rescue, suppression, and SaxA-dependent biocontrol indexes as a function of the maximal OD attained in different conditions of ITC concentrations and pathogen:commensal initial mixing ratio as

$$\text{Rescue index} = 1 - \frac{\text{Max OD}(K \text{ with PSKO})}{\text{Max OD}(K \text{ with PS})} \tag{1}$$

$$\text{Suppression index} = 1 - \frac{\text{Max OD}(PS \text{ with K})}{\text{Max OD}(PS)} \tag{2}$$

$$\text{Biocontrol index} = \text{Rescue index} \cdot \text{Suppression index} \tag{3}$$

that, for the cases discussed in the main text, are numbers ranging from 0 to 1. The rescue index quantifies the conditions where the presence of a SaxA-degrader PS enhance the maximal biomass produced by the commensal compared to the case in the absence of degrader (such as PSKO). The suppression index quantifies the conditions where the maximal biomass attained by the pathogen ITC-degrader PS is repressed by growing in the presence of commensal K, due to nutrient competition. Finally, the SaxA-dependent biocontrol index combines the two previous indexes indicating the conditions when the plant could benefit of a control of the pathogen PS number by the presence of the commensal exploiting the trade-off between the commensal rescue due to sharing the effect of SaxA on ITC degradation and the PS repression because of the nutrient competition with the rescued commensal. To understand better the dynamics of the biomass in different scenario of ITC concentrations and pathogen:commensal initial ratios, we refer to Fig. 12 in the supplementary text, replotted here as upper panel of Fig. 1.

The central panel of Fig. 1A illustrates the case where PS (or PSKO) and K are introduced at equal initial concentrations (PS(KO):K = 1:1). In this condition, the maximal optical density ($OD_{600}$) of K co-cultured with PSKO declines from 0.08 (blue dashed line) to below 0.005 when exposed to ITC concentrations exceeding 15 $\mu$g/mL (red dashed line, lower panel). By contrast, PS, thanks to its ability to degrade ITC, partially rescues K's growth; at 15 $\mu$g/mL ITC, the $OD_{600}$ reaches approximately 0.05 (upper panel, red dashed line). Due to the high ITC-degrading capacity of PS, even a small number of PS cells is sufficient to effectively eliminate ITC and restore K proliferation, as demonstrated in Fig. 1A where the initial PS concentration is reduced by two orders of magnitude relative to K (PS(KO):K = 1:100). In this scenario, the recovered growth of K leads to increased nutrient competition, which in turn suppresses PS proliferation. For instance, at 15 $\mu$g/mL ITC, the maximal $OD_{600}$ of PSKO is 0.08 (solid red line), whereas that of PS drops to 0.06 (lower panel, first column). The impact of nutrient competition becomes even more apparent when comparing PS growth in the presence versus absence of K (Fig. 1B, first rows of panel A vs. panel B). For example, at a PS:K ratio of 1:100 and 60 $\mu$g/mL ITC, PS growth decreases from 0.19 to 0.14 (black solid line), and in the absence of ITC, from 0.19 to 0.05 (blue solid line). Conversely, when the initial PS abundance greatly exceeds that of K (PS(KO):K = 100:1, Fig. 1, rightmost columns), PS monopolizes nutrient resources, resulting in negligible K growth, even when rescued by ITC-degrading PS (dashed lines, upper panel).

We used the dynamics of the biomass for different conditions as presented in Fig. 1 to extract the maximal biomass values (supposed proportional to OD), as summarised in the heat maps of Suppl. Fig. 13. Then, the maximal OD served to compute the values of the indexes for several conditions and summarised in the corresponding heat maps.

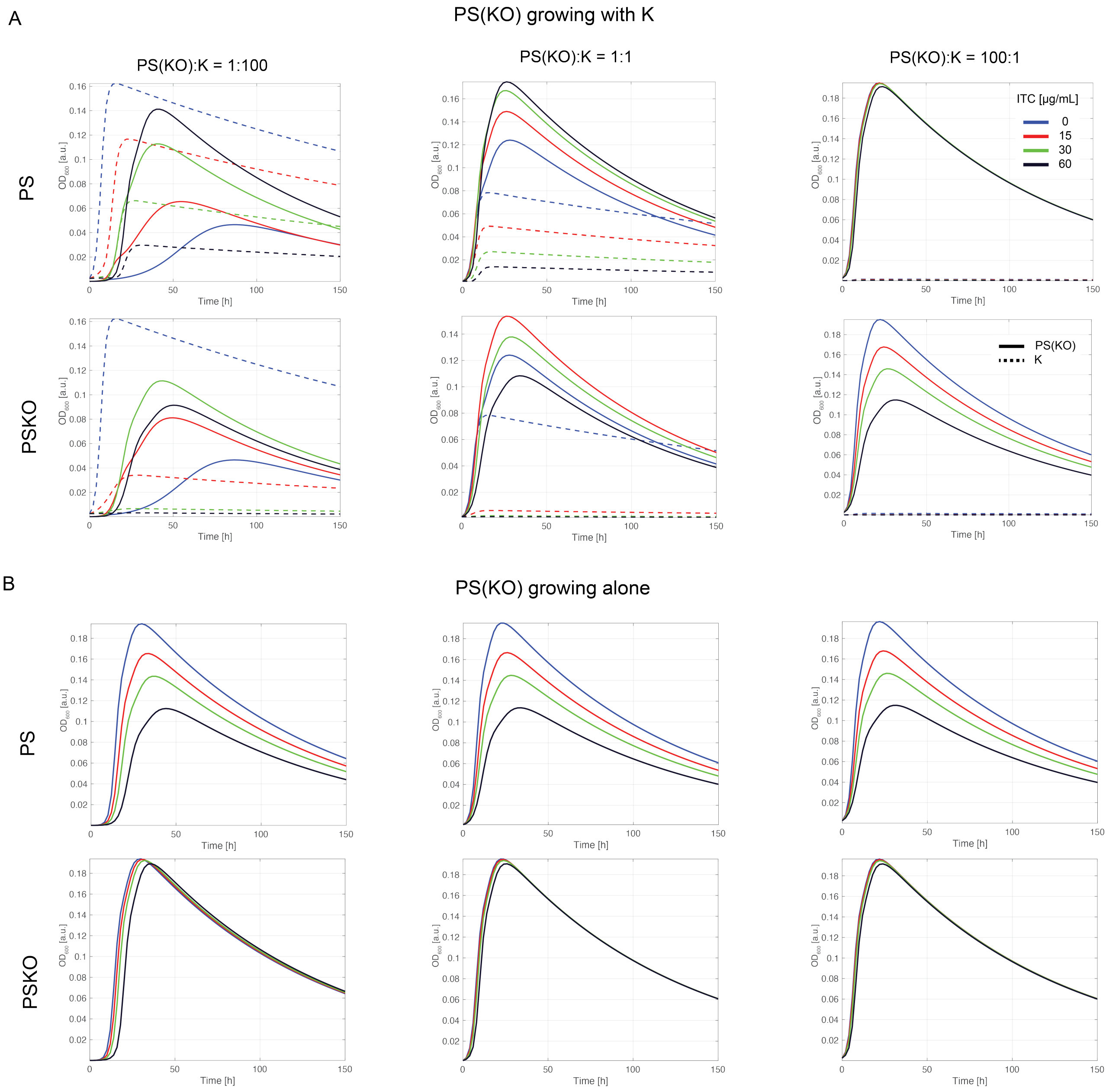


Figure 1: $OD_{600}$ dynamics of pathogens PS and PSKO growing in the absence or presence of commensal K for several initial ratio pathogen:K. (A) OD dynamics of either PS or PSKO (solid lines, top and bottom row, respectively) and K (dashed lines) for several ITC concentrations and three initial pathogen:K ratio (1:100, 1:1, 100:1, first, second and third column, respectively). Panel (B) shows the dynamics of the monocultures of either PS or PSKO, in the absence of commensal K, with the same initial values as in A.
